# Classification of unlabeled observations in Species Distribution Modelling using Point Process Models

**DOI:** 10.1101/651125

**Authors:** Emy Guilbault, Ian Renner, Michael Mahony, Eric Beh

**Affiliations:** School of Mathematical and Physical Sciences, University of Newcastle, Callaghan, NSW, Australia; School of Environmental and Life Sciences, University of Newcastle, Callaghan, NSW, Australia

**Keywords:** Presence-only data, Ecological statistics, Misidentification, Classification, Mixture modelling, EM algorithm, Machine learning

## Abstract

1. Species distribution modelling, which allows users to predict the spatial distribution of species with the use of environmental covariates, has become increasingly popular, with many software platforms providing tools to fit species distribution models. However, the species observations used in species distribution models can have varying levels of quality and can have incomplete information, such as uncertain species identity.
2. In this paper, we develop two algorithms to reclassify observations with unknown species identities which simultaneously predict different species distributions using spatial point processes. We compare the performance of the different algorithms using different initializations and parameters with models fitted using only the observations with known species identity through simulations.
3. We show that performance varies with differences in correlation among species distributions, species abundance, and the proportion of observations with unknown species identities. Additionally, some of the methods developed here outperformed the models that didn’t use the misspecified data.
4. These models represent an helpful and promising tool for opportunistic surveys where misidentification happens or for the distribution of species newly separated in their taxonomy.

## 2 Introduction and background

Species distribution modelling has been a popular topic in ecological statistics over the past decade. Many tools and methods have been developed to provide a means to explore the distributions of species through mapping of suitable environments (Jewell *et al.*, 2007; Peterman *et al.*, 2013; Nezer *et al.*, 2016; Inoue *et al.*, 2017; Schank *et al.*, 2017). Although there are a large number of algorithms and software platforms that can fit species distribution models (SDMs), generalization of these methods and specific applications to real data sets can be tricky (Burnham & Anderson, 2002; Aarts *et al.*, 2012; Guillera-Arroita *et al.*, 2015).

The most common sources of species information used in SDMs are presence-only (PO) and presence-absence (PA) data. PO data only contains information about species presence, in contrast to PA data which records both where species have been found present and where they have not been found (Warton & Shepherd, 2010; Renner *et al.*, 2015). Although PA data is generally of higher quality, it is also less common than PO data because it requires more rigorous planning to visit a set of pre-determined sites. On the other hand, PO data sets are very common, arising from surveys or opportunistic sightings, but they usually have lower quality (van Strien *et al.*, 2013; Ruete & Leynaud, 2015). Point process models (PPMs) are a common tool for fitting SDMs to analyze PO data (Warton & Shepherd, 2010; Mi *et al.*, 2014; Renner *et al.*, 2015) and have been used to fit models for real datasets and simulated data (Baddeley *et al.*, 2006; Illian *et al.*, 2012; Renner & Warton, 2013; Baddeley *et al.*, 2015).

Unreliable or unknown species observation identification is also a main concern in ecology. For example, species records can become confounded when species taxonomy changes (Mahony *et al.*, 2006). Conservation planning efforts depend on clear identification of species and understanding of their distributions and habitat requirements (Franklin, 2013; Guisan *et al.*, 2013). Such concerns are very rarely considered while building SDMs, as people usually clean the data or make some assumptions to avoid such identification problems.

Mixture modelling is a common tool used to represent complex distributions and aims to identify different groups within a dataset while modelling heterogeneity (Martinez, 2015). In communities or groups of individuals/species it is possible to classify or cluster them according to covariate information by using finite mixture modelling (McLachlan & Peel, 2000; Frame & Jammalamadaka, 2007; Dunstan *et al.*, 2013; Fernández-Michelli *et al.*, 2016). One particular application of this approach is to deal with over-dispersed data and to model the different ecological processes at the same time for a single species or for different species in order to classify them (Matthews *et al.*, 2001; Zhang *et al.*, 2004; Tracey *et al.*, 2013).

Machine learning algorithms are also becoming more common in statistical ecology because they can deal with unknown information and recognize some structure in the data (Hastie *et al.*, 2001; Thessen, 2016; Browning *et al.*, 2018). Some algorithms can group observations with similar characteristics (unsupervised learning) and some use separate labeled datasets (supervised learning) or partially labeled data within the studied dataset (semi-supervised learning) to classify the observations (Wendel *et al.*, 2015; Fernández-Michelli *et al.*, 2016; Vo *et al.*, 2018; Zhou, 2018). Some recent publications have applied machine learning algorithms to fit PPMs in a Bayesian framework (Tran, 2017; Vo *et al.*, 2018), but the literature on using machine learning algorithms to fit PPMs is not yet well-developed. Additionally, several R packages have been developed to deal with machine learning procedures (Benaglia *et al.*, 2009; Iovleff, 2018), but none accommodate the intersection of point process modelling with mixture modelling or machine learning algorithms.

In this paper we develop new tools for fitting models to multi-species PO data with partial species identification by combining the PPM framework with mixture modelling and machine learning approaches to accommodate incomplete labelling. These tools implement two algorithms to reclassify the unreliable observations to belong to one of the existing species. The first tool fits mixtures of PPMs to all available data with an Expectation-Maximization (EM) algorithm and uses them to classify the unreliable points. This method will be called *Mixture method*. The second tool employs an iterative technique to fit separate PPMs to points with known labels augmented by some points with unknown labels depending on classification probabilities at each iteration. This method will be hereafter known as the *Loop method*. Using simulations, we compare the performance in classification and prediction for the proposed algorithms to the simple, standard approach of fitting individual PPMs to the points with known species labels only. We found that performance varied based on the choice of initialization and algorithm parameters but some of the methods can outperform the fitting of individual PPMs.

## 3 New modelling methods

### 3.1 Notation

The fitted point process models in our proposed methods make use of a total of *M* + *N* + *Q* locations as follows:

Let *s*_1_ = {*s*_1_, …, *s*_*m*1_}, s_2_ = {*s*_*m*1+1_, …, *s*_*m*1+*m*2_}, …, **s**_*K*_ = {*s*_*M−mK* +1_, …, *s*_*M*_} be vectors that contain all of the observed locations with known species identities 1, 2, …, *K*, respectively. These are represented by the orange, purple, and turquoise dots in Figure 1 for a hypothetical dataset. Let |**s**_1_| = *m*_1_, |**s**_2_| = *m*_2_, …, |**s**_*K*_| = *m*_*K*_ be the number of observed locations with known species identity for each of the *K* species. We collect the *M* = *m*_1_ + *m*_2_ + … + *m*_*K*_ total locations with known species identities of all *K* species in **s** = {**s**_1_, **s**_2_, …, **s**_*K*_}. Let **u** = *{s_M_*_+1_, …, *s*_*M*+*N*_} contain the *N* observed locations with uncertain species identities. These are represented by the black question marks in Figure 1. Let **q** = {*s*_*M+N+1*_, …, *s*_*M+N+Q*_} contain the locations of *Q* quadrature points placed along a regular *c*_1_ × *c*_2_ grid throughout the study region (Figure 1). Each quadrature point is placed at the center of one of *Q* unique rectangular grid cells throughout the study region. Let *c*(*s*) be the grid cell in which location *s* is contained.

**Figure 1:**
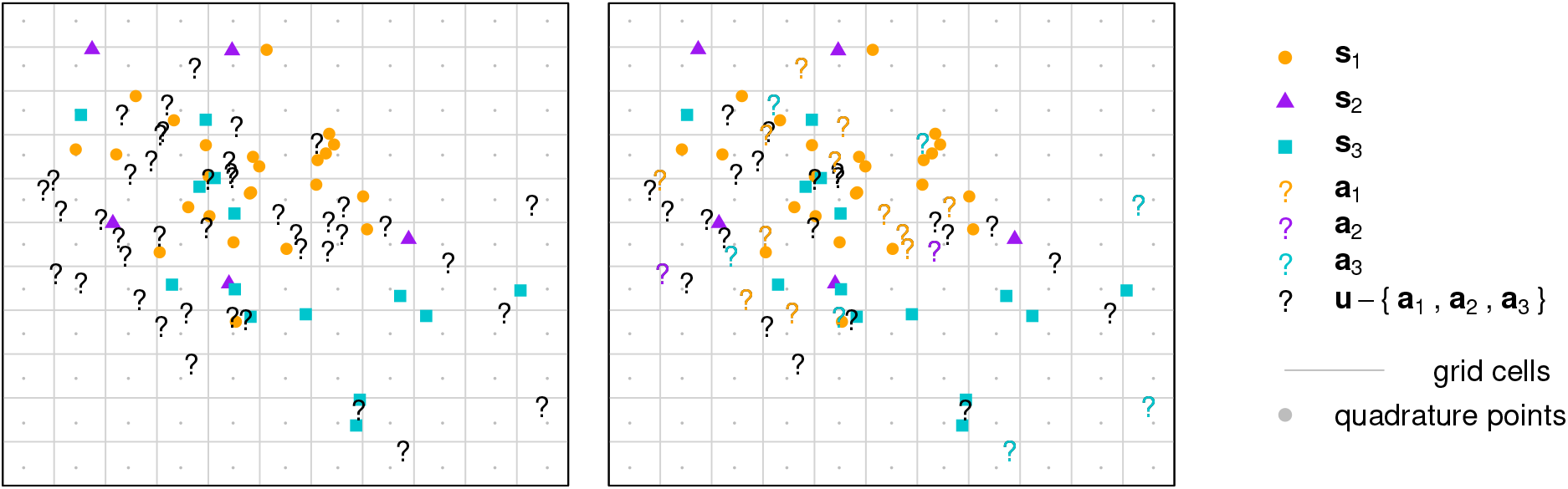
Three illustrative point patterns. The orange, purple, and turquoise colored dots represent locations with known species identity, **s**_1_, **s**_2_, and **s**_3_. The gray dots represent quadrature points **q**, which are spaced evenly along a regular grid such that one quadrature point is at the centre of each rectangular grid cell. The black question marks (left) represent observed locations **u** with uncertain species identity. The locations in **a**_1_ ∈ **u**, **a**_2_ ∈ **u**, and **a**_3_ ∈ **u** which are reclassified as belonging to one of the species are represented by coloured question marks (right).

### 3.2 Loop methods

The three loop algorithms proceed by iterating between steps that augment the vectors of locations with known species identities **s**_1_, **s**_2_, …, **s**_*K*_ with locations **a**_1_ *⊂* **u**, **a**_2_ ⊂ **u**, …, **a**_*K*_ ⊂ **u**, update the quadrature weights, and fit point process models as follows:

1. Fit *K* initial point process models using the vectors of observed locations with known species identity **s**_1_, **s**_2_, …, **s**_*K*_.
2. Compute the predicted intensities 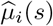 for all *s* ∈ {**s** ∪ **u**} for *i* ∈ {1, …, *K*}.
3. Derive an (*M* + *N*) × *K* matrix of membership probabilities ***ω***, where

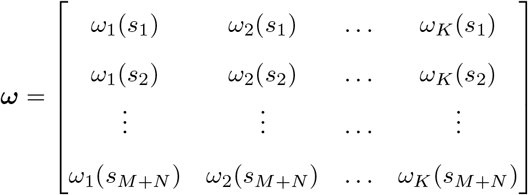 The membership probability of location *s* for species *i* is defined as

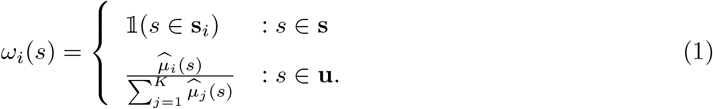 That is, the membership probabilities for the locations with known species identity are 1 for the correct species and 0 otherwise, and for the locations with unknown species identity, they are proportional to the fitted intensities.
4. Define an augmented vector for species *i* as **y**_*i*_ = **s**_*i*_ ∪ **a**_*i*_ for all *i* ∈ {1, …, *K*}. We define **a**_*i*_ as follows:
  - For the **Normal** method, **a**_*i*_ = **u** (left panel of Figure 2).

**Figure 2:**
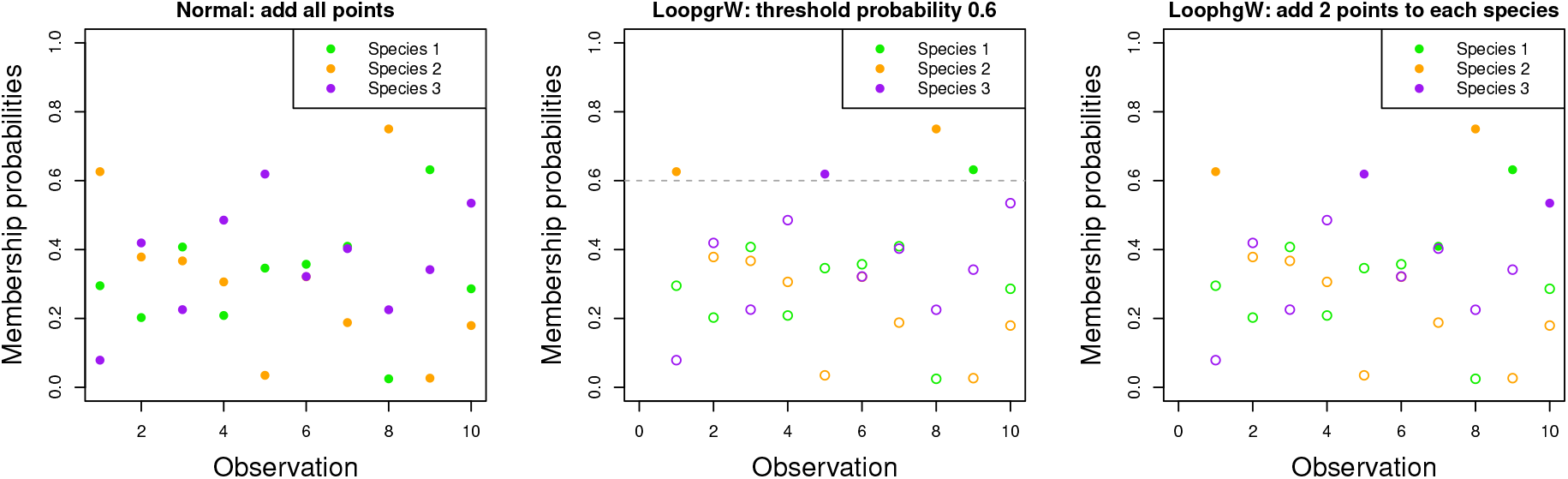
(Left) Normal Loop function. We add all points with unknown species labels to each species, using membership weights that are proportional to the fitted intensities. (Middle) Method Loop grW function. We add all points with membership probabilities greater than a threshold *δ*_max_, then we decreases from that value to a minimum of *δ*_min_ by increments of *δ*_step_. (Right) Method Loop hgW function. We add the *a* points with highest membership probabilities to each species, increasing the number *a* from 1 to 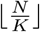.
  - For the **Loop grW** method, 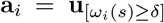, where *δ* is a minimum membership probability threshold that takes the following values successively at each iteration {*δ*_max_, δ_max_ − *δ*_step_, …, *δ*_min_}. That is, the Loop grW method augments the locations with known species identity *i* with the locations with unknown species identity with membership probabilities for species *i* that are higher than the current threshold *δ* (middel panel of Figure 2).
  - For the **Loop hgW** method, 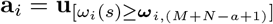, where ***ω***_*i*,(*j*)_ represents the *j*^th^ smallest entry of vector ***ω***_*i*_, the *i*^th^ column of ***ω***, and *a* represents the number of locations to be augmented. We set *a* to be the same integer for all *K* species for some *a* between 1 and 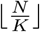 then at each iteration a is increased by one (right panel of Figure 2).
5. Update the quadrature weights for each species. First, assign each location in {**y**_1_, …, **y**_*K*_, **q**} to a grid cell. Then, compute the vector of quadrature weights **w**_*i*_ for all points *t* ∈ {**y**_*i*_ ∪ **q**} as follows:

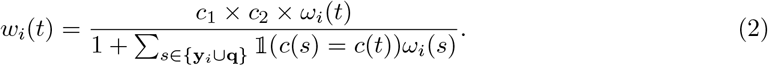 This way of computing quadrature weights is an extension of standard quadrature weight schemes for point process models (Berman & Turner, 1992), in which the weight for location *s* is equal to the area of the grid cell *c*(*s*) that contains *s* divided by the total number of quadrature and observed locations in *c*(*s*). Here, we divide the area of the grid cell by the sum of the membership probabilities of the observed locations in the grid cell (both with and without known species identities) plus 1 (for the one quadrature point in the grid cell).
6. Fit point process models using the augmented vector **y**_*i*_, quadrature points **q** and quadrature weights **w**_*i*_ for all species *i* ∈ {1, …, *K*}.
7. Return to step 2 and stop when we either reach likelihood convergence or we reach a maximum number of iterations that is different depending on the method chosen. Likelihood convergence is determined by:

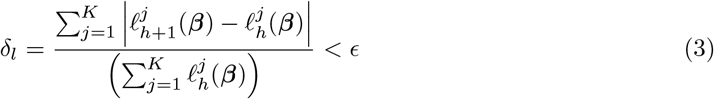

for some choice of *ε*, where 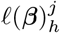 is the fitted log-likelihood for the *j*^th^ species at the *h*^th^ iteration.

The maximum number of iterations varies for the different methods, as follows:

- For the **Normal** method, the maximum number of iterations is set by the user. We set the default number of iterations to be 50.
- For the **Loop grW** method, the maximum number of iterations is determined by the choice of *δ*_max_, *δ*_step_, and *δ*_min_.
- For the **Loop hgW** method, the maximum number of iterations is 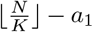, where ⌊*c*⌋ rounds the number *c* down to the nearest integer, and *a*_1_ is the first value of *a* chosen by the user. In the case of decimals numbers, only the floor is considered as the we can’t add more points than available per species.

### 3.3 Mixture of PPMs method

The four mixture algorithms can be fitted by maximizing a log-likelihood function and reclassifying the locations with uncertain identity using an EM algorithm framework. The algorithm proceeds as follows:

1. We initialize the membership probabilities ***ω*** for each location *s* for each species *i* in one of the following ways:

- For the **knn method**, we calculate the distance *d*_*i*_(*s*) of each location *s* to the *k*^th^ nearest neighbor of species *i*, for all *K* species. We calculate the membership probability of location *s* for species *i* using:

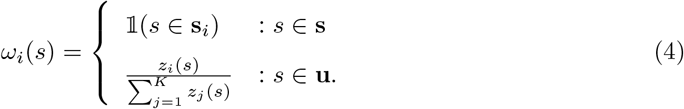

where

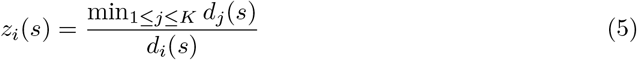
- For the **kmeans method**, we define *ω*_*i*_(*s*) as in (4) but define *z*_*i*_(*s*) as

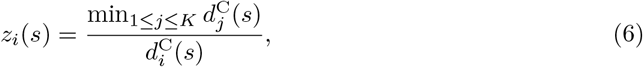

where 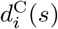 is the distance to the *i*^th^ centroid of the *i*^th^ cluster.
- For the **random method**, we define *ω*_*i*_(*s*) as in (4) and *z*_*i*_(*s*) is drawn randomly from a uniform distribution:

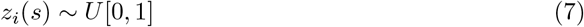
- For the **equal method**, we assign equal membership probabilities for the locations with uncertain identity:

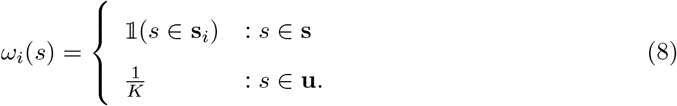 Regardless of the initialization method, the sum of membership probabilities across the all species is equal to 1 for all points.
2. Classify the locations in **u** to belong to one of the *K* species based on the membership probabilities ***ω***.
3. Fit a point process model using a marked point pattern, where each observation *s* has a mark defined by the known or classified identity among the *K* species.
4. Compute the predicted intensities 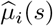 for all *s* ∈ {**s** *∪* **u**} for *i* ∈ {1, …, *K*}.
5. E step: We first get the predicted values of each species at the locations *s* ∈ {**s** *∪* **u**} and calculate the predicted intensity of the mixture of *K* densities using:

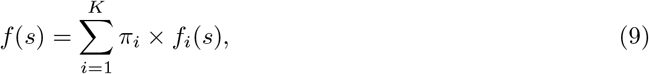

where *f*_*i*_(*s*) is the density at location *s* for the *i*^th^ component and *π*_*i*_ is the mixing proportion or weight of the *i*^th^ species in the mixture.
6. We calculate new membership probabilities for each unknown point of **u** using:

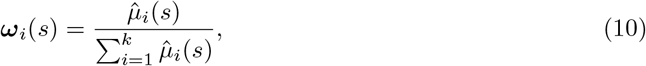

where *µ*_*i*_(*s*) is the intensity of the *i*^th^ species at location *s* ∈ **s**. For the observations **s** with known labels, the membership probabilities are set to 1 for the correct species label and 0 otherwise.
7. M step: Classify the locations in **u** to belong to one of the *K* species. The classification for each point *s* corresponds to the highest membership probability *ω*_*i*_(*s*) for *i* ∈ {1, …, *K*}. We compute each species’ proportion of the whole by summing the membership probabilities for each species across both **s** and **u**.
8. Compute a marked PPM based on the updated classifications and membership probabilities.
9. Calculate the model log likelihood using:

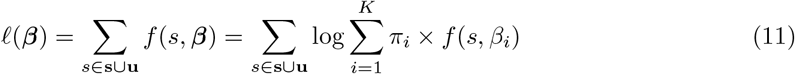
10. Repeat steps 4-9 until we achieve likelihood convergence, defined as follows:

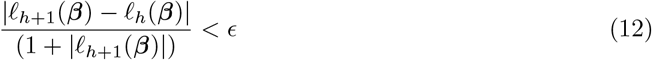

where *l*_*h*_(***β***) is the log-likelihood at the *h*^th^ iteration and ε is a pre-specified tolerance level.

## 4 Simulation framework

### 4.1 Simulation data

To compare the performance of the different algorithms, we simulated patterns **t**_1_, **t**_2_, and **t**_3_ of individuals for three species based on “true” distributions defined by four different predictors. Because performance could varied based on sample size, the correlations *ρ_i,j_* among the species distributions, and the proportion of observations with unknown labels, we consider similar and different low abundances by randomly simulating numbers of points between 20 and 50 for the species as well as the correlation between the true species distributions:

- Case 1: at least two species *i* and *j* have distributions that are highly correlated (|*ρ*_*i,j*_| ≥ 0.85 for some *i, j* ∈ {1, 2, 3})
- Case 2: no two species have highly correlated distributions (|*ρ*_*i,j*_| < 0.45 for all *i, j* ∈ {1, 2, 3})

We chose these values for abundances as they would be small enough such that potential value of adding points with unknown species identities could be investigated, and we chose these cutoffs for correlation to create clearly distinguishable contexts.

We then created locations with unknown labels **u** by hiding uniformly at random a certain proportion of the total observations (20%, 50% and 80%). The locations in **t**_1_, **t**_2_, and **t**_3_ that retained their true species identities therefore became the simulated point patterns **s**_1_, **s**_2_, and **s**_3_ with known species identities. Simulations were conducted using the version 3.4.2 of R (R Core Team, 2017) and used high performance computing to implement 1000 simulations each for different combinations of abundances, correlation among species distributions, and proportions of observations with unknown labels. We also tested different parameters for the knn initialization of the mixture algorithm (the value of *k* neighbors), the Loop grW function (the maximum threshold *δ*_max_, minimum threshold *δ*_min_ and the step size *δ*_step_) and the Loop hgW function (initial number of points added to the point pattern *a*).

### 4.2 Suite of Evaluation tools

We consider various measures of performance for comparing the distributions. For classification methods, misclassification/accuracy analysis is a common measure of performance (Wendel *et al.*, 2015). We choose the highest mixing weight for each observation to determine the labeling when computing accuracy. We also compared the final membership probabilities of the correct labels of each point to 1 (the true weight) with a residual sum of squares (RSS).

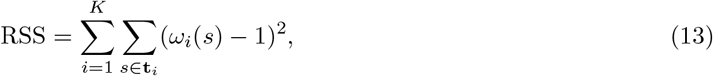

where *ω*_*i*_(*s*) is the final membership probability for location *s* for the correct species *i* computed using the methods outlined in sections 3.2 and 3.3. Considering residual sum of squares (RSS) alone does not provide a reliable comparison because the number of unknown observations can vary, so we consider meanRSS instead to standardize the measure for all fitted models:

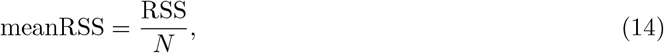

where *N* is the number of observations with uncertain species identities.

We also considered measures that compare the true distribution from which we generate the points to the predicted distributions of the model. We use a sum of correlations between the true and predicted distributions across all species (hereafter referred to as ‘sumcor’) to assess how well the predicted distributions align with the true distributions. We can use various correlation measures such as Pearson’s correlation coefficient, Kendall’s *τ* or Spearman’s *ρ* when computing sumcor.

Another global measure of predictive performance of the intensity estimates is the Integrated Mean Square Error (IMSE) (Swanepoel, 1988; Es, 1997). The function is defined as:

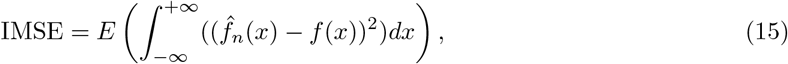

where 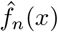 is an estimator of the density function *f* (*x*). We standardized this value by rescaling the intensities to be able to compare each methods even if different number of points are considered and compute the IMSE using the values of the true and predicted intensities at the quadrature points **q**, and sum across the 3 species.

## 5 Results

Here we present the results of the simulations, with more detailed results appearing in the Appendix. In this section, we only present the results from the knn, Lopp grW, Loop hgW and individual PPM methods that displayed the best performances. First, we present the model performances from varying data parameters (abundance, correlation and percentage of hidden labeled data). The individual PPM results will be used as a point of comparison with the other methods as the individual method does not include any of the points with unknown labels. We, then, focus on varying model parameters in the different methods (the value of *k* for knn, the values of *δ*_max_, *δ*_min_ and *δ*_step_ for Loop grW and the value of *a* for Loop hgW). For these results, we set *k* = 1, *δ*_max_ = 0.5, *δ*_min_ = 0.1, *δ*_step_ = 0.1 and *a* = 5 according to the algorithm parameters tests presented in section 5.2. For the performance results, the sumcor methods displayed the result using the Pearson correlation coefficient.

### 5.1 Varying species distributions

#### 5.1.1 Different abundances and correlated distributions

In Figure 3, we consider different low abundances (*m*_1_ = 32, *m*_2_ = 42 and *m*_3_ = 23) and where two distributions are highly correlated. With regard to classification performance, the different modelling methods have similar levels of accuracy, although when comparing meanRSS, the individual and Loop grW methods seem to outperform the other methods, especially as we increase the proportion of hidden observations. With regard to predictive performance, the Loop grW method appears to have the greatest performance when measured by IMSE and sumcor, particularly for 50% and 80% of hidden observations. The Loop hgW method performs comparably to the individual PPM method, although its preformance gets relatively better as we increase the proportion of hidden observations. The knn method has the highest IMSE for 50% and 80% of hidden observations, but it is competitive with the individual PPM and loop hgW method when comparing sumcor. See Tables 1 and 2 in the Appendix for a comparison of means and medians across all of these measures.

**Figure 3:**
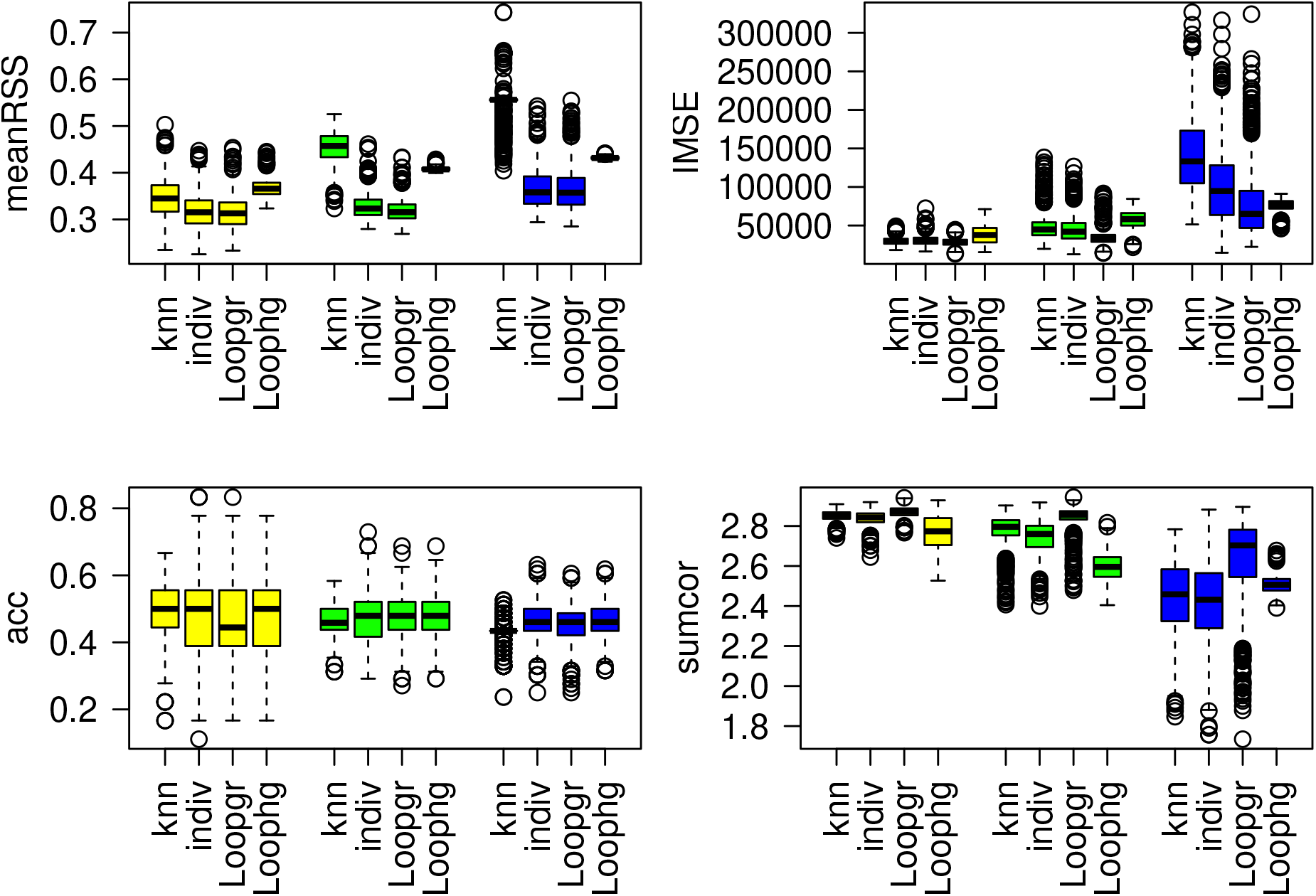
Measures of performance for the knn, individual, Loop grW and Loop hgW methods. Each color boxplot represents a different percentage of hidden observation: in yellow are the performances with 20% of hidden observations, in green with 50% and in blue with 80%. The parameters of abundances and correlation are: *m*_1_ = 32, *m*_2_ = 42, *m*_3_ = 23; *ρ*_1,2_ = 0.85, *ρ*_1,3_ = −0.09, *ρ*_2,3_ = 0.20.

When examining the predicted intensities with 80% of the observations with hidden species identities, the true pattern appears best captured by the Loop grW method (Figure 4), consistent with sumcor. The Loop hgW method tends to overpredict the intensities.

**Figure 4:**
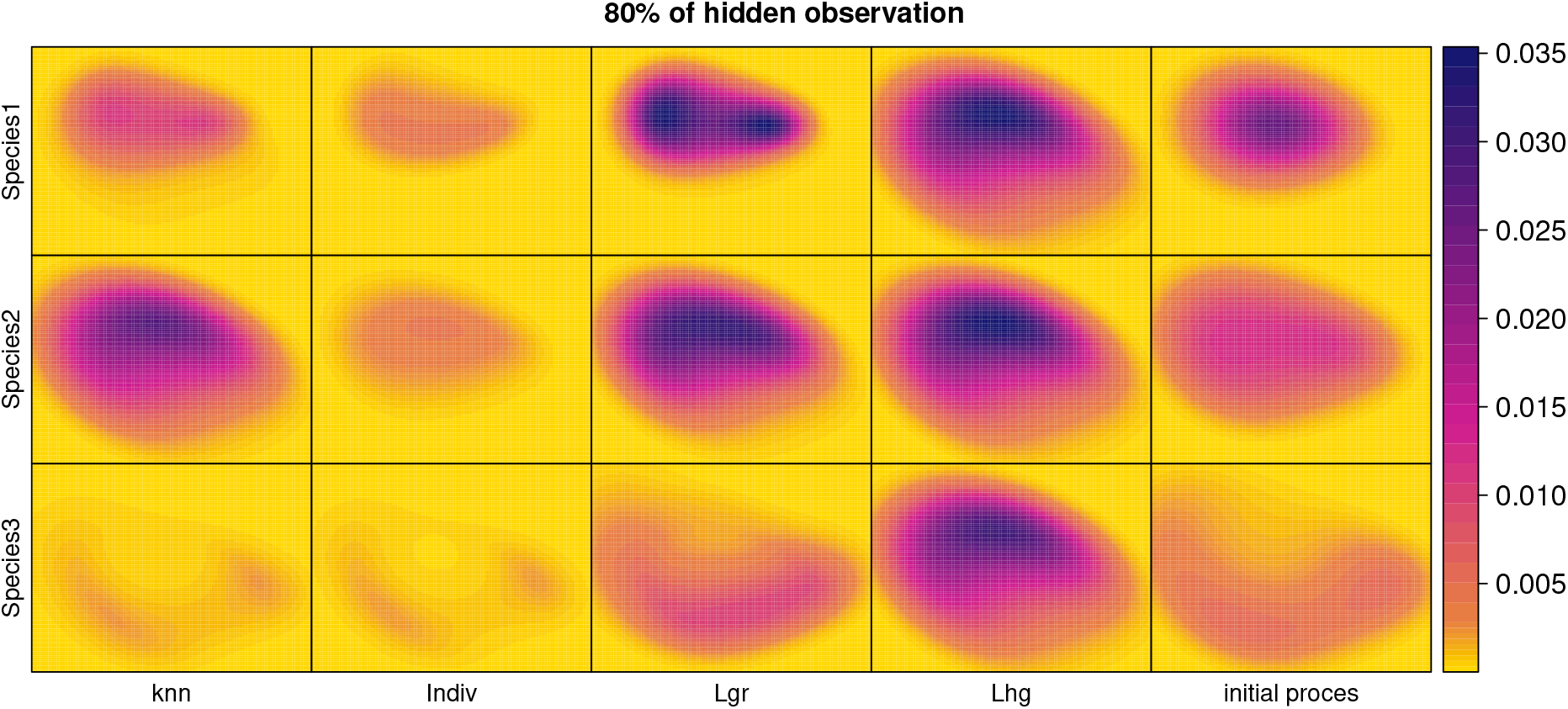
Predicted intensities obtained for the knn, individual, Loop grW and Loop grW methods and the initial intensities from the process with 80% of hidden observations. The parameters of abundances and correlation are: *m*_1_ = 32, *m*_2_ = 42, *m*_3_ = 23; *ρ*_1,2_ = 0.85, *ρ*_1,3_ = *−*0.09, *ρ*_2,3_ = 0.20.

#### 5.1.2 Similar abundances and correlated distributions

In Figure 5, we consider similar abundances (*m*_1_ = 33, *m*_2_ = 34 and *m*_3_ = 35) and where two distributions are highly correlated. With regard to classification performance, the different modelling methods have similar levels of accuracy, except the knn method does relatively poorly with 80% of the observations hidden. The knn method also suffers worse performance as measured by meanRSS at 50% and 80% of hidden observations. Measures of predictive performance are similar to the case with different abundances and correlated distributions. The Loop grW method appears to outperform the others as the proportion of hidden observations increases, with the Loop hgW method competitive with the individual PPM method. The knn method appears to do worse with 80% hidden observations when measured by IMSE. See Tables ?? and ?? in the Appendix for comparisons of means and medians across all of these measures. With 80% hidden observations, the Loop Loop grW method appears to be best aligned with the true intensities, as shown in Figure 6.

**Figure 5:**
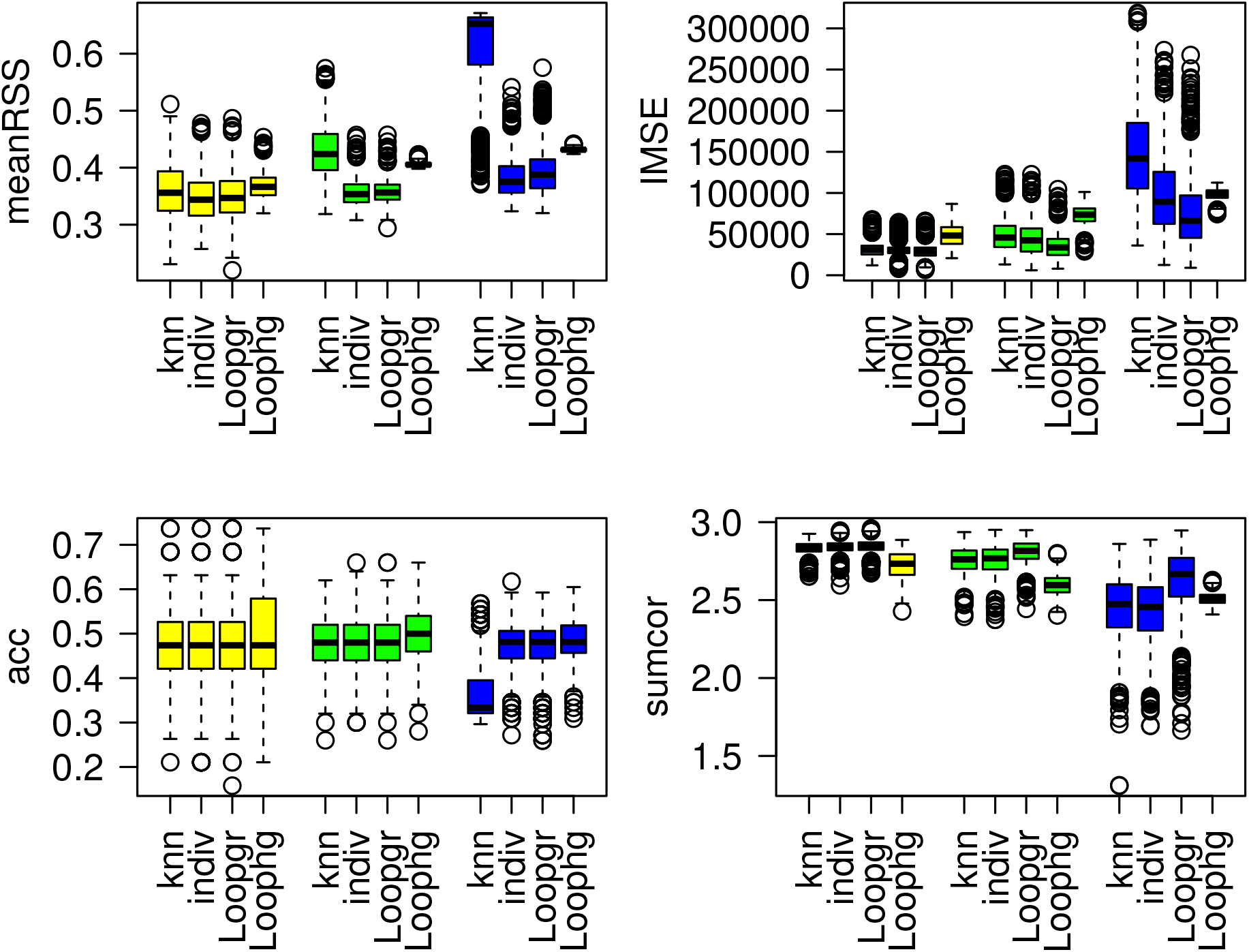
Measures of performance for the knn, individual, Loop grW and Loop hgW methods. Each color represents a different percentage of hidden observations: in yellow are the performances with 20% of hidden observations, in green with 50% and in blue with 80%. The parameters of abundances and correlation are: *m*_1_ = 33, *m*_2_ = 34, *m*_3_ = 35; *ρ*_1,2_ = 0.85, *ρ*_1,3_ = *−*0.09, *ρ*_2,3_ = 0.20.

**Figure 6:**
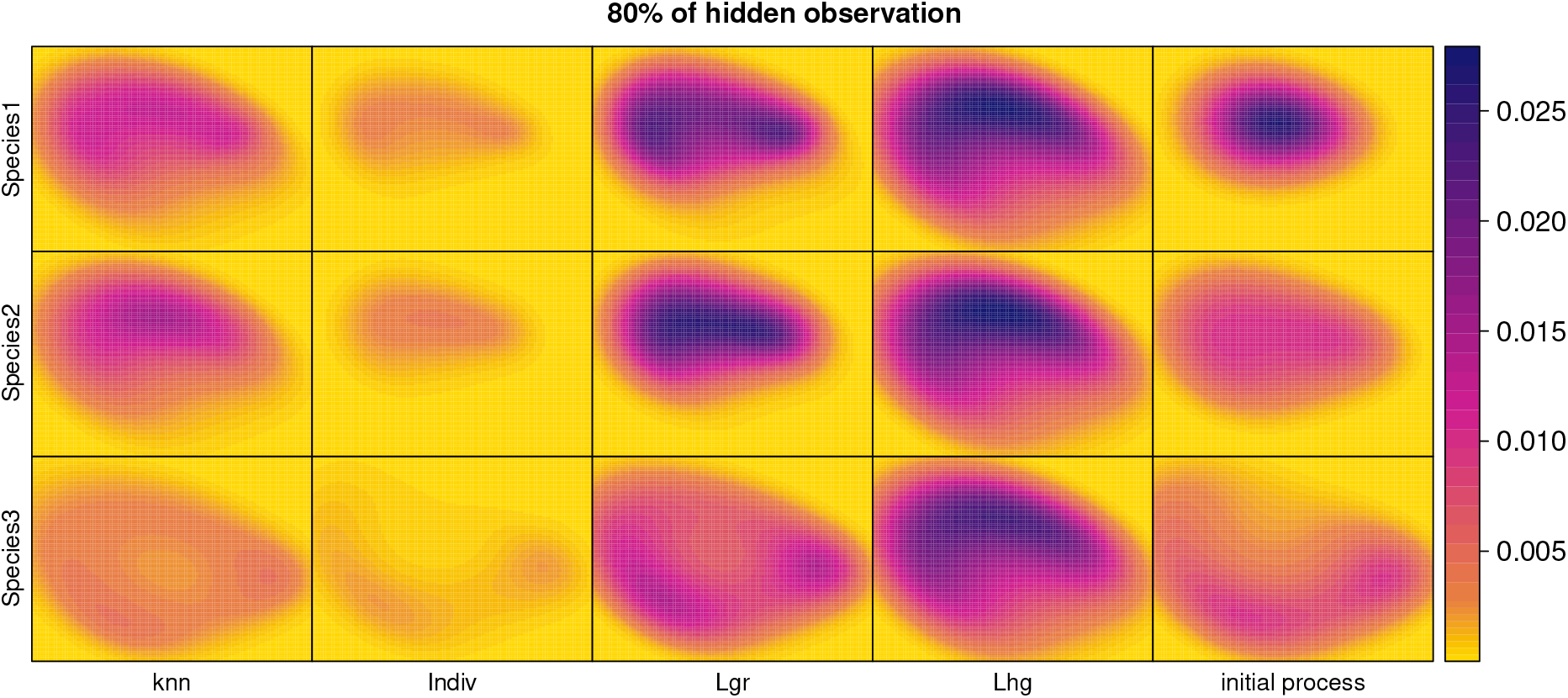
Predicted intensities obtained for the knn, individual, Loop grW and Loop hgW methods and the initial intensities from the process with 80% of hidden observations. The parameters of abundances and correlation are: *m*_1_ = 33, *m*_2_ = 34, *m*_3_ = 35; *ρ*_1,2_ = 0.85, *ρ*_1,3_ = *−*0.09, *ρ*_2,3_ = 0.20.

#### 5.1.3 Different abundances and non correlated distributions

In Figure 7, we consider different abundances (*m*_1_ = 42, *m*_2_ = 31 and *m*_3_ = 25) and where none of the distributions have high correlations. The classification performance and predictive performance comparisons look similar to the case of similar abundances and correlated distributions as shown in Figure 5, with the knn method having the worst classification performance described here at 50% and 80% of hidden observations and the Loop grW method outperforming the others in predictive performance, while the Loop hgW method is competitive with the individual PPM method and the knn method lags behind with IMSE at 80% of hidden observations. Tables 5 and 6 in the Appendix contains the means and medians across all performance measures for this context.

**Figure 7:**
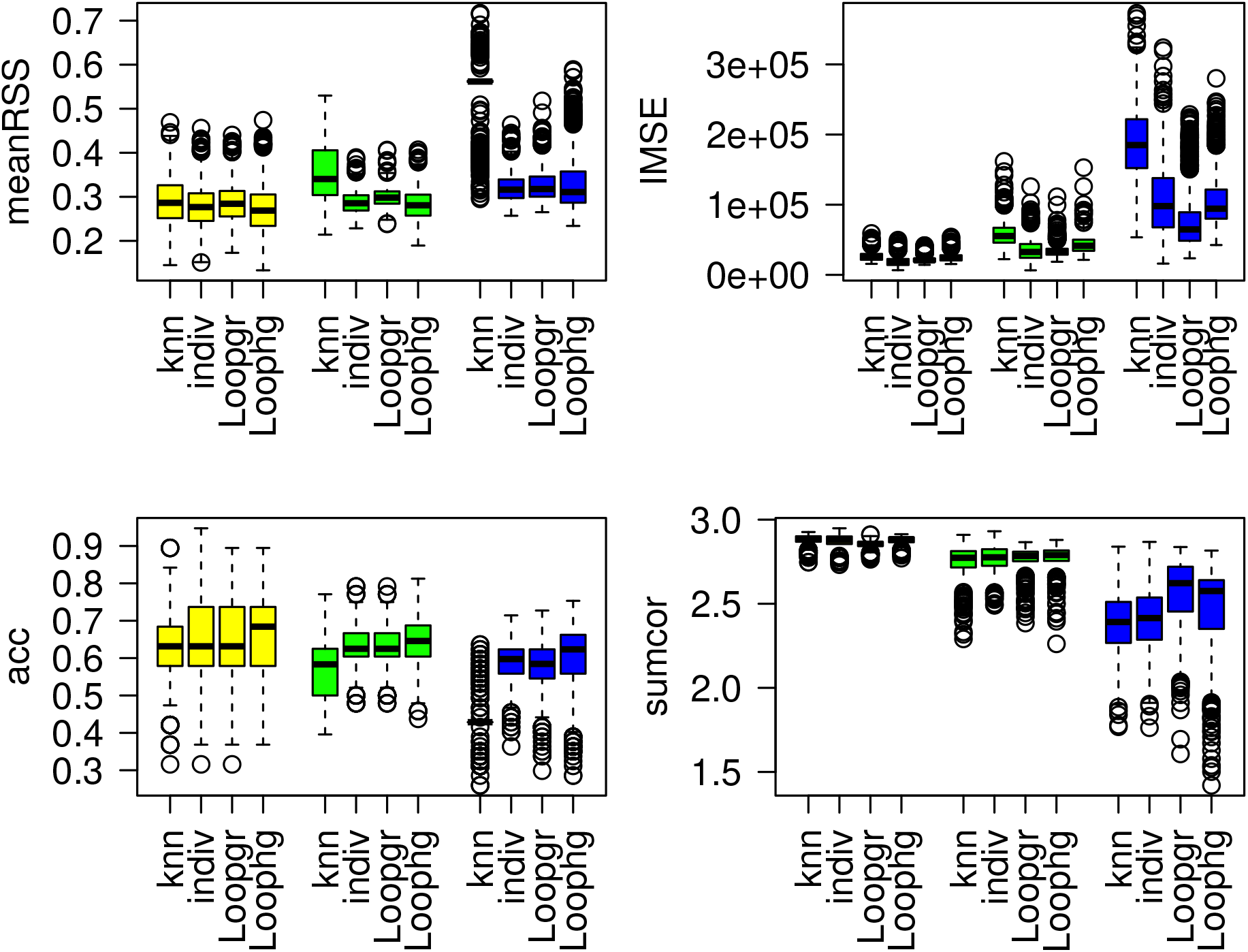
Measures of performance for the knn, individual, Loop grW and Loop hgW methods. Each color represents a different percentage of hidden observations: in yellow are the performances with 20% of hidden observations, in green with 50% and in blue with 80%. The parameters of abundances and correlation are: *m*_1_ = 42, *m*_2_ = 31, *m*_3_ = 25; *ρ*_1,2_ = 0.09, *ρ*_1,3_ = *−*0.42, *ρ*_2,3_ = 0.20.

With 80% of hidden observation as shown in Figure 8, the Loop hgW method for species 1 and 3 and the Loop grW method for species 2 and 3 are the closest to the initial process.

**Figure 8:**
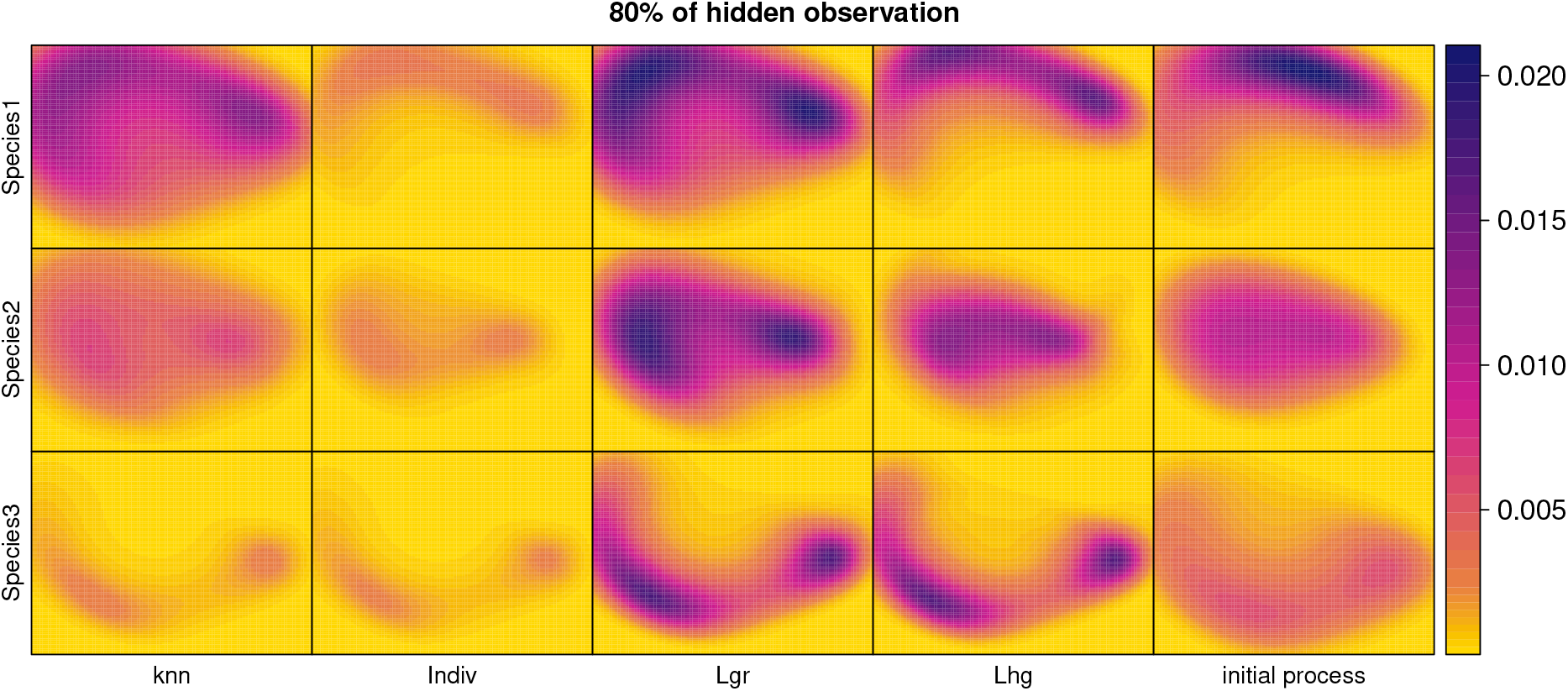
Predicted intensities obtained for the knn, individual, Loop grW and Loop hgW methods and the initial intensities from the process with 80% of hidden observations. The parameters of abundances and correlation are: *m*_1_ = 42, *m*_2_ = 31, *m*_3_ = 25; *ρ*_1,2_ = 0.09, *ρ*_1,3_ = *−*0.42, *ρ*_2,3_ = 0.20.

#### 5.1.4 Similar abundances and non correlated distribution

For similar abundances (*m*_1_ = 39, *m*_2_ = 37, *m*_3_ = 38) and non correlated distributions, we again observe the same trends, as shown in Figure 9: the knn method is the worst method for relabeling performances and the only one not doing as well as the individual method for 50% and 80% of hidden observations. As in previous contexts, the Loop grW method shows the best predictive performance, with the Loop hgW method being competitive with the individual PPM method, and the knn method having higher IMSE than the other methods when 80% of the observations are hidden. Tables 7 and 8 in the Appendix contain the mean and median value for all performance measures.

**Figure 9:**
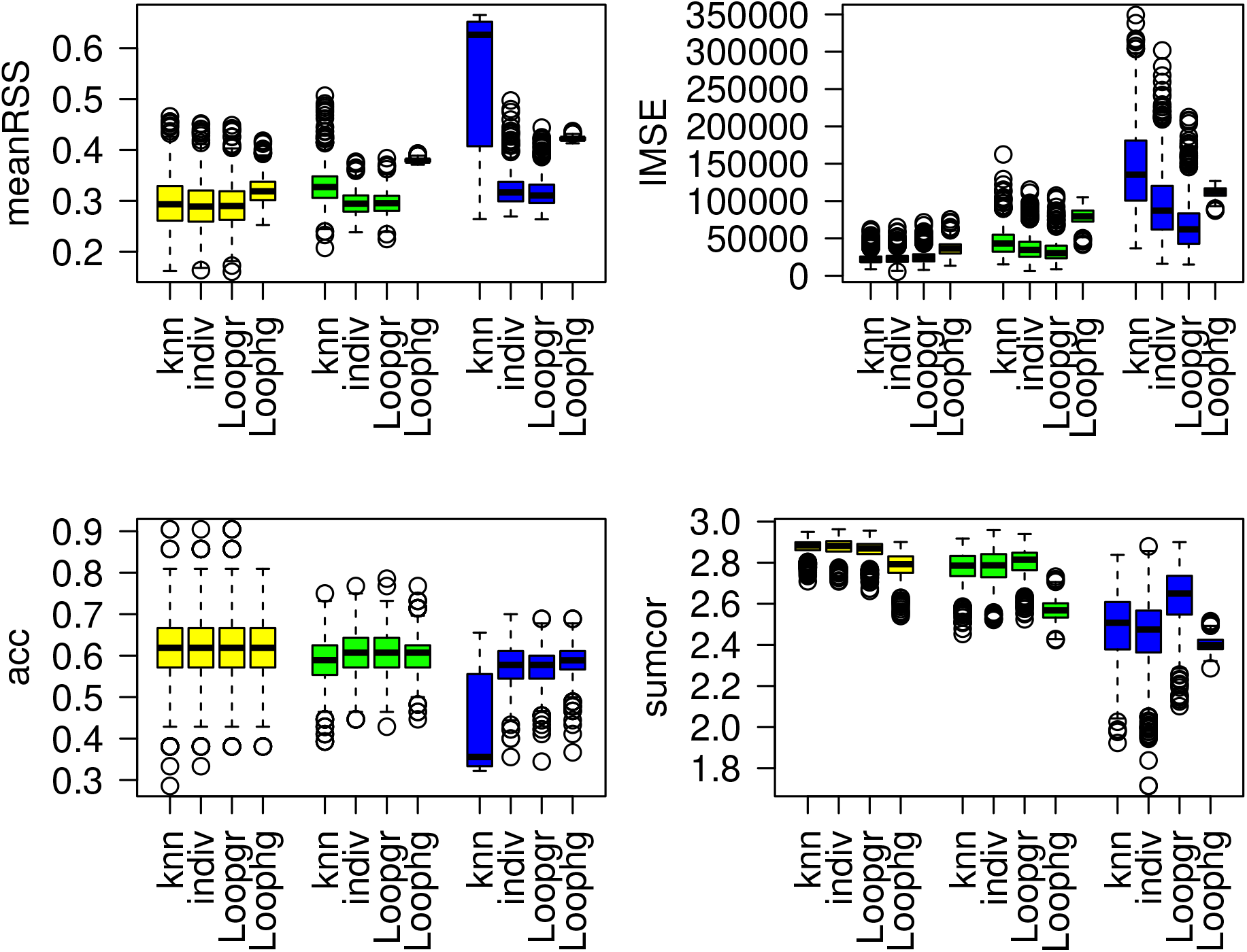
Measures of performance for the knn, individual, Loop grW and Loop grW methods. Each color represents a different proportion of hidden observations: in yellow are the performances with 20% of hidden observations, in green with 50% and in blue with 80%. The parameters of abundances and correlation are: *m*_1_ = 39, *m*_2_ = 37, *m*_3_ = 38; *ρ*_1,2_ = 0.09, *ρ*_1,3_ = *−*0.42, *ρ*_2,3_ = 0.20.

The predicted intensities show the methods LgrW and knn being the closest to the initial process, as shown in Figure 10.

**Figure 10:**
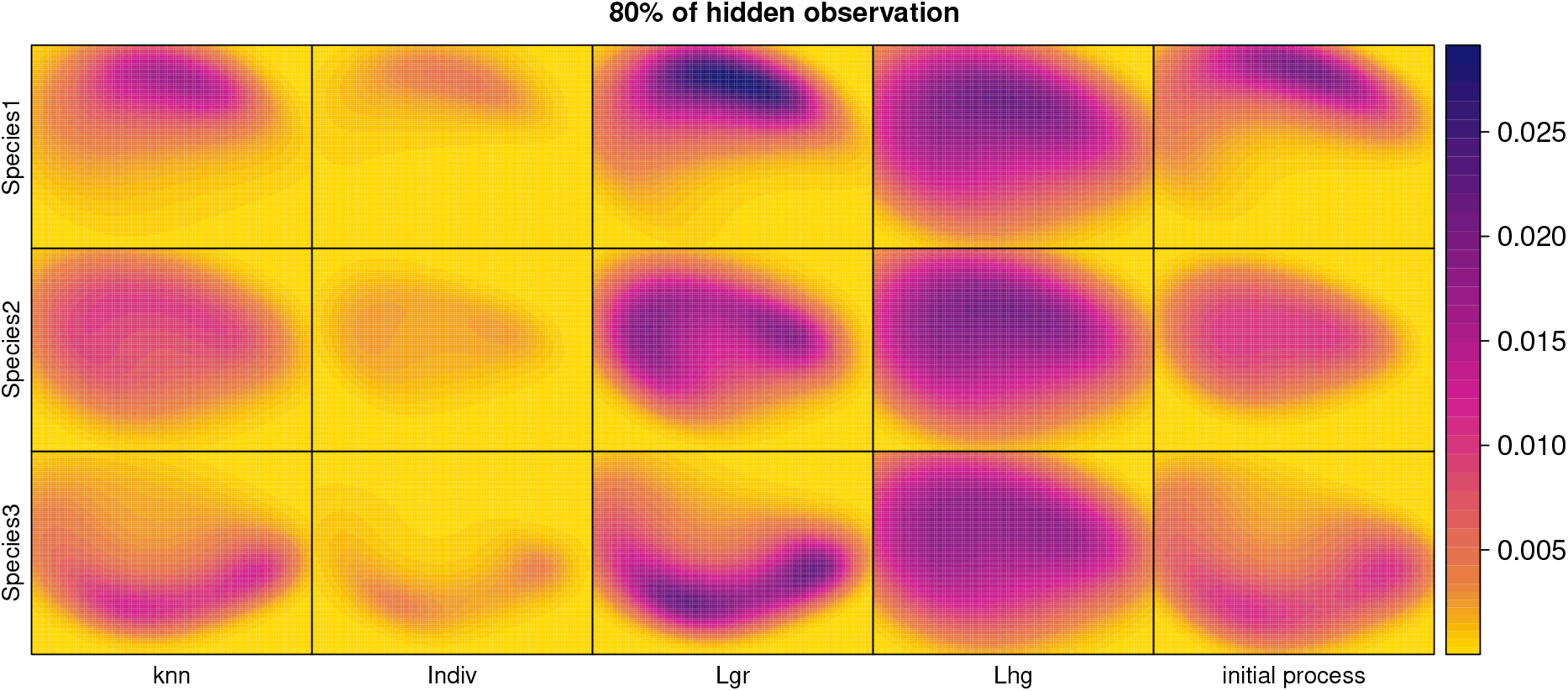
Predicted intensities obtained for the knn, individual, Loopg rW and Loop grW initialization methods and the initial intensities from the process at 80% of hidden observations. The parameters of abundances and correlation are:*m*1=39, *m*2=37, *m*3=38; *ρ*_1*−*2_=0.09, *ρ*_1*−*3_=-0.42, *ρ*_2*−*3_=0.207

### 5.2 Testing algorithm parameters

#### 5.2.1 knn method

We note that when the *k* nearest neighbor value increases (from 1 up to 20), the model performances decrease; Figure 11. It is particularly notable for the performances in prediction where sumcor performances decrease and IMSE performances increase. Also, there is an expected drop in performances as we increase the proportion of observations with unknown species labels.

**Figure 11:**
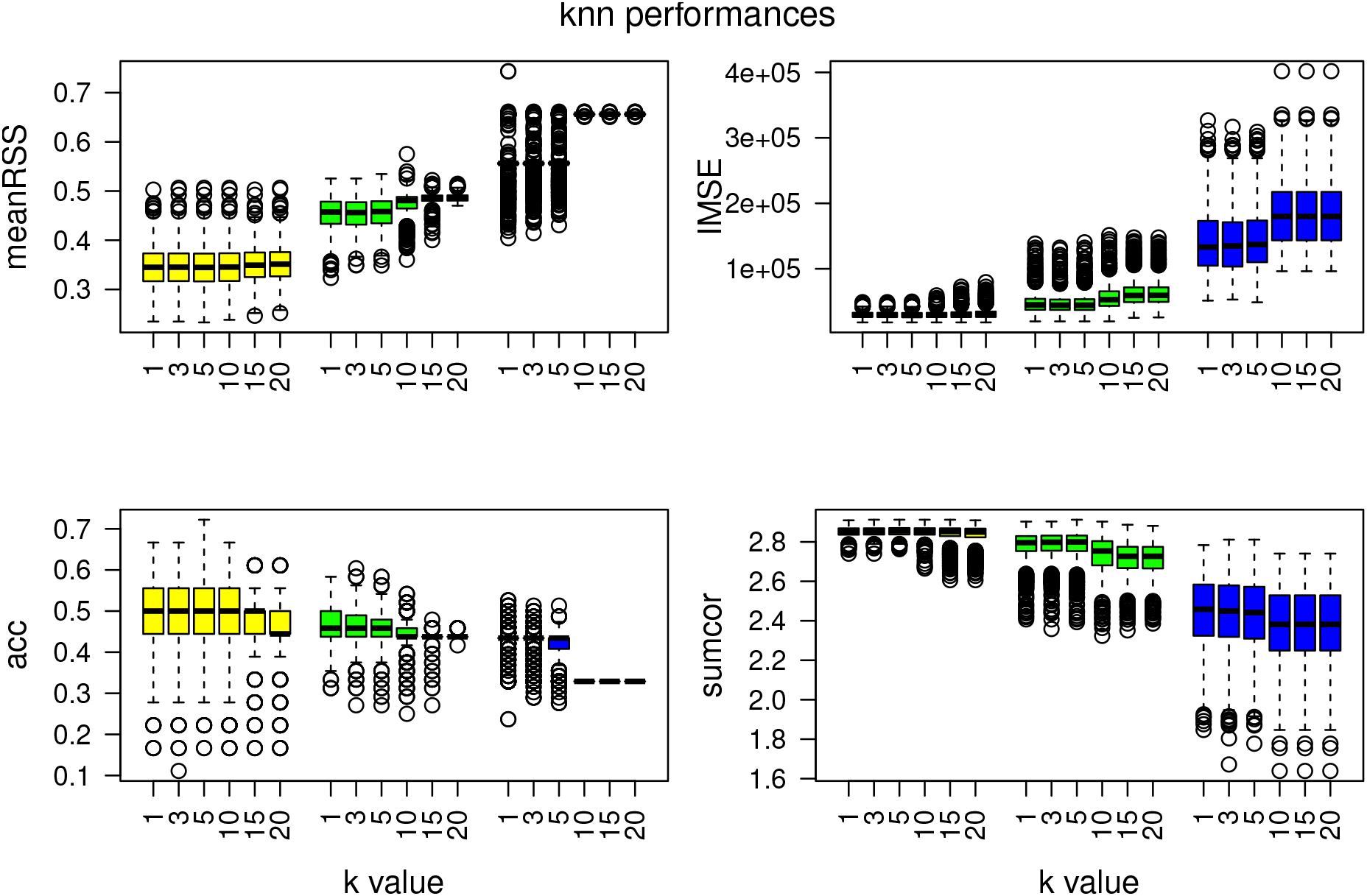
Model performances for the knn method. Each color represents a different percentage of hidden observations: in yellow are the performances with 20% of hidden observations, in green with 50% and in blue with 80%. The parameters of abundances and correlation are: *m*_1_ = 32, *m*_2_ = 42, *m*_3_ = 23; *ρ*_1,2_ = 0.85, *ρ*_1,3_ = *−*0.09, *ρ*_2,3_ = 0.20

#### 5.2.2 Loop grW method

For the Loop grW method we tested different parameters:

1. The initial membership probability threshold *δ*_max_: while this parameter varies from 0.8 to 0.5 in increments of 0.1, the other Loop grW parameters are as follows: *δ*_min_ = 0.1 and *δ*_step_ = 0.1.
2. The final membership probability threshold *δ*_min_: while this parameter varies from 0.1 to 0.7 in increments of 0.2, the other Loop grW parameters are as follows: *δ*_max_ = 0.8 and *δ*_step_ = 0.1.
3. The step size *δ*_step_: while this parameter varies from a minimum of 0.01 to a maximum of 0.2, the other Loop grW parameters are as follows: *δ*_max_ = 0.8 and *δ*_min_ = 0.1.

When we change the value of *δ*_max_, there is very little difference in performance within each proportion of observations with hidden labels, although *δ*_max_ = 0.5 appears to be slightly superior to the other choices for high percentage of hidden observation (Figure 12).

**Figure 12:**
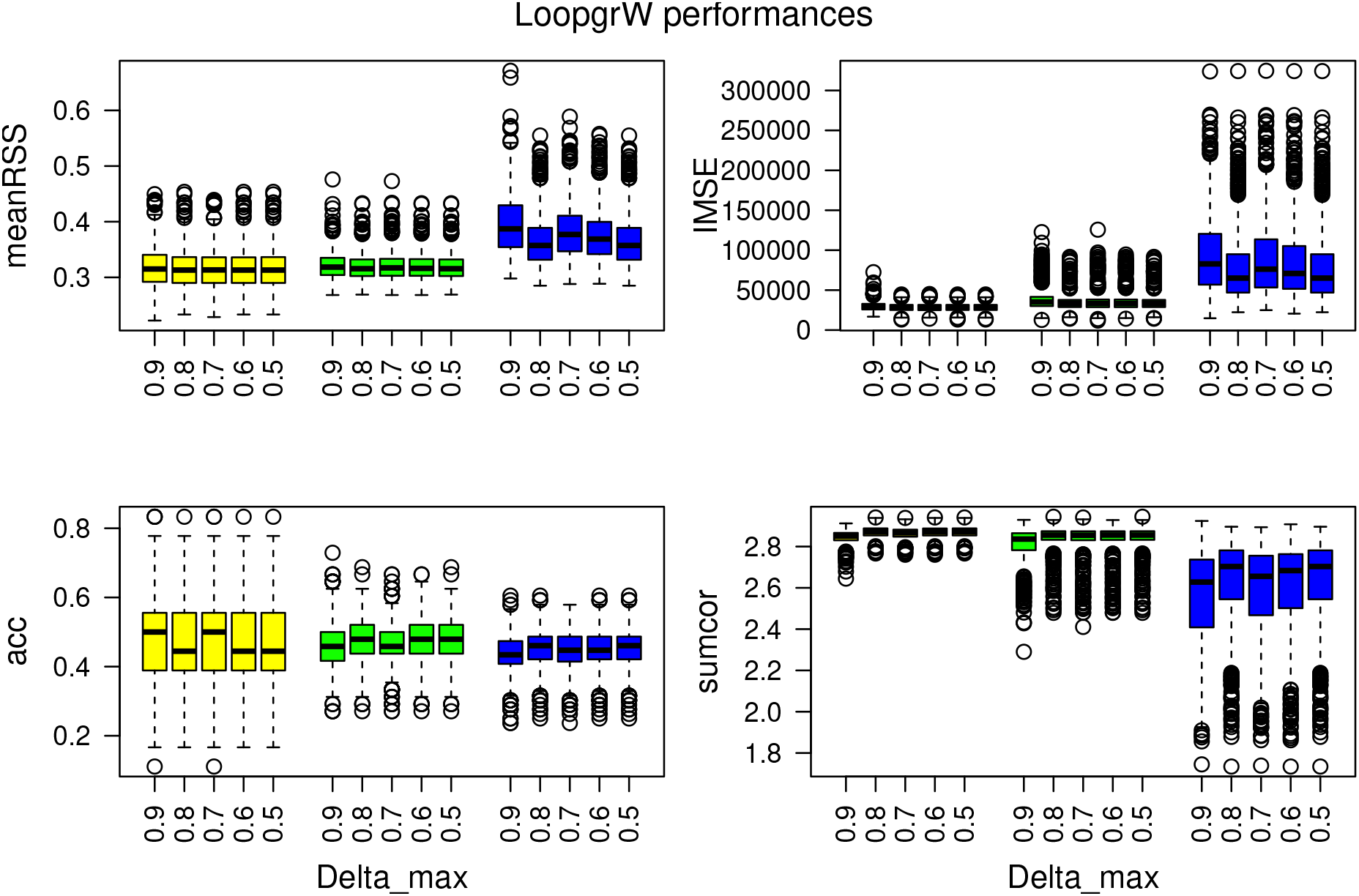
Model performances for the Loop grW method and for different values of *δ*_max_. Each color represents a different proportion of hidden observations: in yellow are the performances with 20% of hidden observations, in green with 50% and in blue with 80%. The parameters of abundances and correlation are: *m*_1_ = 32, *m*_2_ = 42, *m*_3_ = 23; *ρ*_1,2_ = 0.85, *ρ*_1,3_ = *−*0.09, *ρ*_2,3_ = 0.20

When changing *δ*_min_, the classification accuracy is relatively the same (Figure 13). For MeanRSS, IMSE and sumcor, we can observe a curved pattern of performances, where the performances decrease (MeanRSS increases, IMSE increases and sumcor decreases) from *δ*_min_ from 0.1 to 0.5 and then the performances get slightly better (MeanRSS decreases, IMSE decreases and sumcor increases) for *δ*_min_ = 0.7 (Figure 13). *δ*_min_=0.1 displays the better performances.

**Figure 13:**
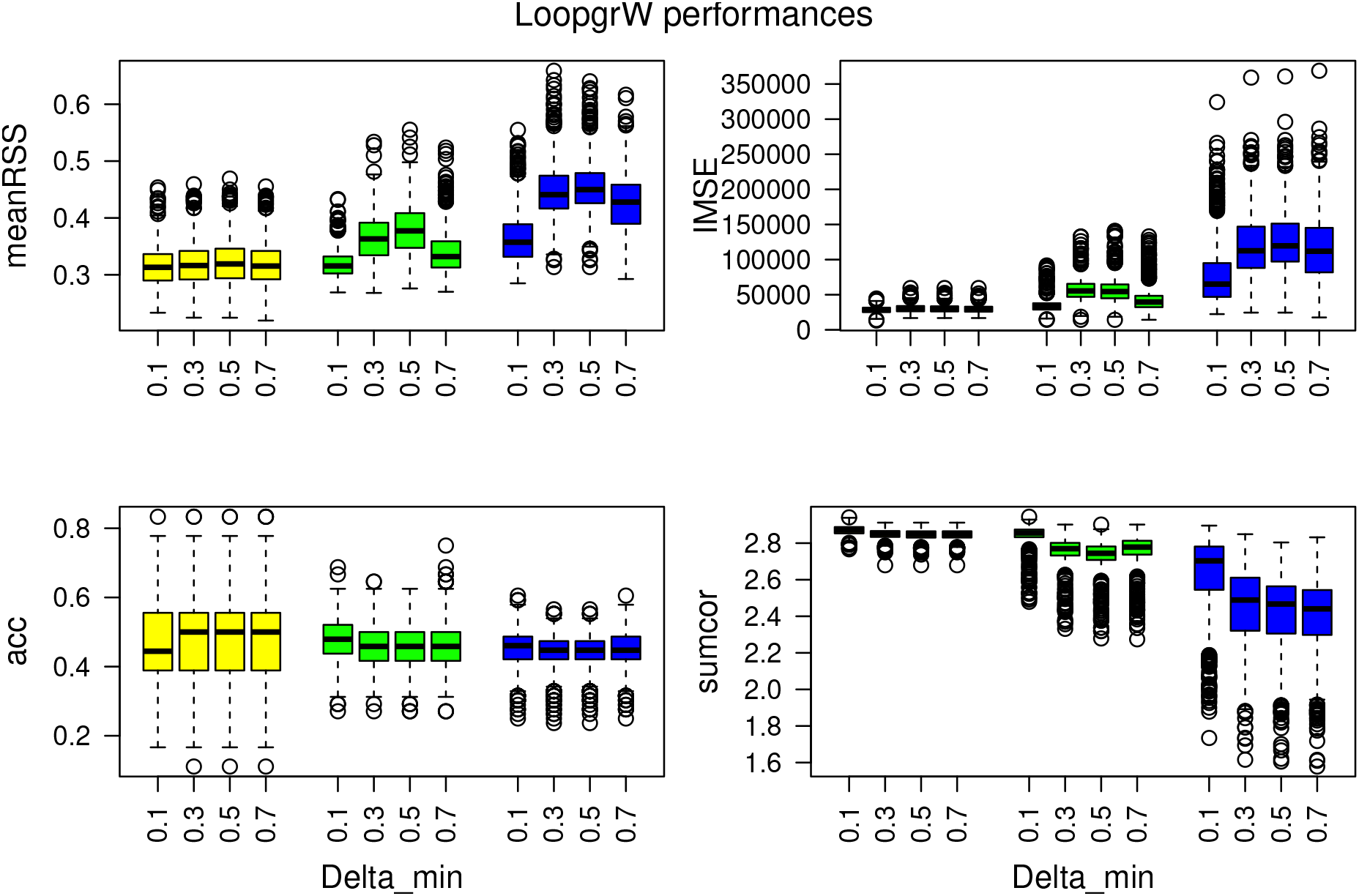
Model performances for the Loop grW method and for different values of *δ*_min_. Each color represents a different proportion of hidden observations: in yellow are the performances with 20% of hidden observations, in green with 50% and in blue with 80%. The parameters of abundances and correlation are: *m*_1_ = 32, *m*_2_ = 42, *m*_3_ = 23; *ρ*_1,2_ = 0.85, *ρ*_1,3_ = −0.09, *ρ*_2,3_ = 0.20

Figure 14 shows different performance measures as we vary *δ*_step_. There do not appear to be major differences in classification performance, although 0.1 appear slightly better for meanRSS. With 50% and 80% of hidden observations, predictive performance display a curve performances where performances get better (IMSE decreases and sumcor increase) from 0.01 till 0.1 and then get worse (IMSE increases and sumcor descreases) from 0.1 to 0.2. *δ*_step_=0.1 displays the best performances accross all measures.

**Figure 14:**
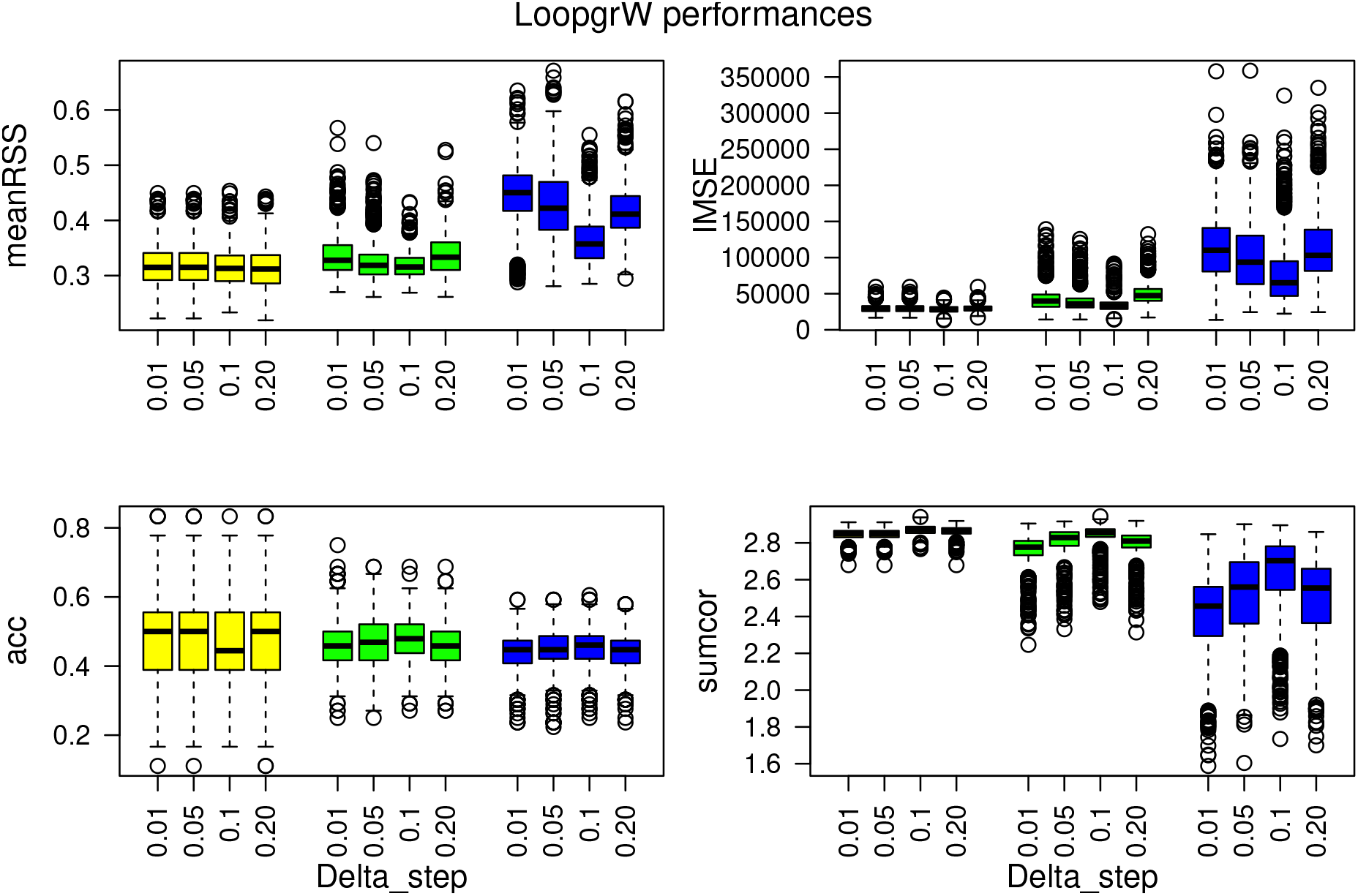
Model performances for the Loop grW method and for different values of weight step. Each color represents a different proportion of hidden observations: in yellow are the performances with 20% of hidden observations, in green with 50% and in blue with 80%. The parameters of abundances and correlation are: *m*_1_ = 32, *m*_2_ = 42, *m*_3_ = 23; *ρ*_1,2_ = 0.85, *ρ*_1,3_ = −0.09, *ρ*_2,3_ = 0.20

#### 5.2.3 Loop hgW method

In the Loop hgW method, we vary the number of points *a* added at each iteration. In Figure 15, we can see that there is no variation in performances when the number of added points *a* increases.

**Figure 15:**
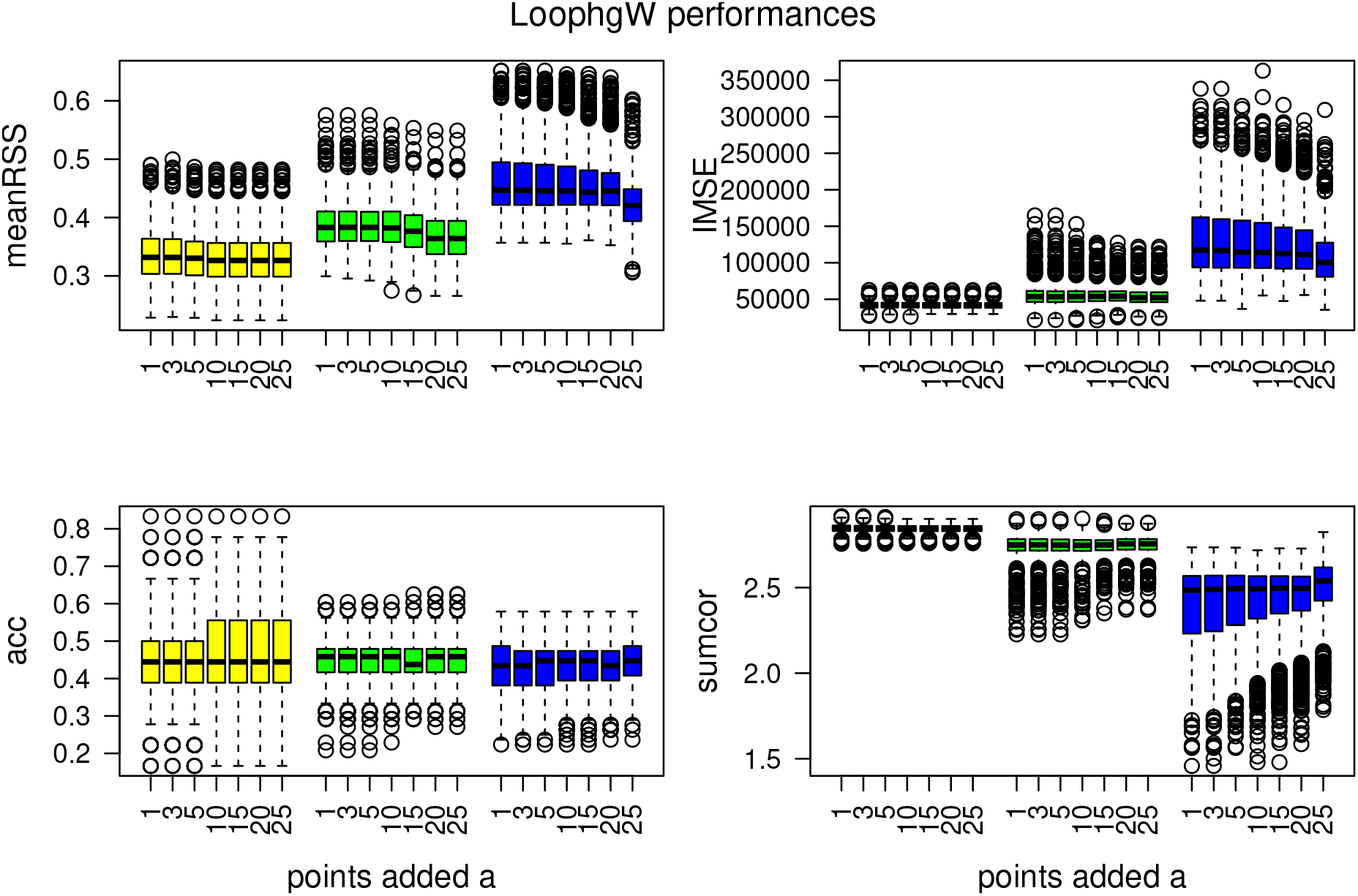
Model performances for the Loop grW method. Each color boxplot represents a different percentage of hidden observations: in yellow are the performances for 20% of hidden observations, in green for 50% and in blue for 80%. The parameters of abundances and correlation are: *m*_1_ = 32, *m*_2_ = 42, *m*_3_ = 23; *ρ*_1,2_ = 0.85, *ρ*_1,3_ = *−*0.09, *ρ*_2,3_ = 0.20

The results for the other combination of abundances and correlation are showed in the Appendix.

## 6 Discussion

In this article, we present a new modelling tool in R that aims to incorporate the observed locations with unknown species identities to improve species distributions. These tools accommodate two ways of reclassifying information using mixture modelling and the machine learning framework with 7 different initialization methods. We tested our algorithms in different contexts where we vary the abundances of our species (similar or different), the correlation between them (two distribution are correlated or none are correlated) and the proportion of unknown species identities (20%, 50% and 80%). The different methods were compared to the individual method which ignores locations with unknown species identities to see whether the proposed algorithms allow us to fit distributions that are closer to the initial processes.

In the results we presented the three best methods. They showed varying performance depending on the aspects of the model and the performance measure considered. The novelty of these tools, makes it difficult to compare to other existing tools that either do not consider point pattern process (Frame & Jammalamadaka, 2007; Frühwirth-Schnatter, 2006; Hui, 2016; Martinez, 2015; Melnykov & Maitra, 2010; Quost & Denœux, 2016), Poisson distributions (Figueirido & Jain, 2002; Hui *et al.*, 2015; Scrucca *et al.*, 2016; Woillez *et al.*, 2012), count data (Benaglia *et al.*, 2009; Iovleff, 2018; Leisch, 2004) or implementation of mixture (Witten, 2011; Wendel *et al.*, 2015) or semi-supervised learning frameworks (Di Zio *et al.*, 2007; Fraley & Raftery, 1998; Jeffries & Pfeiffer, 2001; Taddy & Kottas, 2012).

The other methods (kmeans, random, equal and normal) not presented previously in the results are presented in the Appendix. They show relatively worse performance across all measures, although at times, the normal loop method is competitive with the individual PPM and the Loop hgW methods. We note that this method performs slightly better when the distributions are correlated.

We have noticed differences in performance, that are more significant when we increase the proportion of observations with hidden labels. While at 20% of hidden observations, all methods performed fairly similarly, at 50% and 80% of hidden observations, the loop grW method in particular showed the best predictive performances regardless of differences in abundance and correlation among species distributions. For this method, only the points with the highest membership probabilities are added. We set the maximum and minimum thresholds at *δ*_max_ = 0.5 and *δ*_min_ = 0.1 and a step size of *δ*_step_ = 0.1, but we could expect that performances may be better or worse with different choices of these parameters as shown in the results. These choices appeared to produce superior performances for most measures than other values of these parameters considered. Higher values of *δ*_min_ led to worse performances. This result can be seen as counterintuitive as we can expect that having a smaller interval of weight for example could improve this particular performances. It will in other words reduce the interval of weights and better discriminate the points of uncertain identity. As for *δ*_step_, choosing a value that is too small may lead to iterations where no points are added, while choosing a value that is too large may be too discriminating and does not allow to reclassify the points.

The Loop hgW method did not perform as good as the Loop grW method even if it has been shown to be as good as the individual method in some contexts. For this method, we add initially a certain number of points *a* that is increased at each iteration. While the *a* points with highest membership probabilities are added, these membership probabilities may be small for large values of *a*, and this could explain that this method is not always doing as good as the best method.

Interestingly, the knn method was the best of the four mixture methods tested, outperforming the kmeans, random and equal initialization options. Previous studies using the EM algorithm for classification and clustering data show that such algorithms are highly dependent on the initialization method (Figueirido & Jain, 2002; Melnykov & Maitra, 2010; O’Hagan *et al.*, 2012). Additionally, even very popular methods like kmeans have some drawbacks. Its performance is dependent on overlapping densities and whether the distributions are roughly circular or not. The choice of the centroid is also not consistent and chosen at random for the first calculation (Yoo *et al.*, 2012, 2007; Wu *et al.*, 2008). In our simulations, kmeans, random and equal methods showed very different results and always performed worse than the other methods as well as mainly overestimating (kmeans and random) and underestimating (equal) the predicted intensities compared to the true process.

Despite outperforming the other mixture modelling methods, the knn method was still not competitive with the machine learning methods or the individual PPM method when the proportion of hidden observations are 50% or 80%. However, the knn method was quite consistent in the predicted intensities and showed similar results to the individual method for the sumcor measure at 50% or 80% of hidden observations. Other studies have found that the performance of the knn method is linked to the metric chosen to calculate the nearest neighbor distances and the value of the number *k* of nearest neighbors (Weinberger & Saul, 2009; Guo *et al.*, 2003; Wu *et al.*, 2008).

We tested how the number of neighbors *k* can influence the model and found that for any combination of abundance and correlation, all the measures of performances decrease when the values of *k* increase. It is expected as the neighboring points are further away from one another and could conflate species habitat preferences with differing species abundances, but requiring more neighbor points can also stabilize the distances. The way of choosing the value of *k* by utilizing different distance metrics could also impact the performances as previously noted, but we shall leave this aspect of the analysis for future consideration.

In our simulations, we have considered a relatively general case of point patterns and we only varied species abundance and correlation among distributions in addition to the proportion of observations with hidden information. For real ecological data sets, there are more factors to consider that can influence how a model will perform. First, the abundances tested in the simulation are quite low (20-40 points) and some methods can show convergence issues in this context. While we use the spatstat package (Baddeley *et al.*, 2015) to fit PPMs, we could make use of similar functions in the ppmlasso package (Renner & Warton, 2013) which integrate regularization methods like the lasso penalty that can boost performances with low sample sizes. A related point is that we included all covariates that were used to generate the true point patterns in our models. In real situations, however, we may not have access to the best covariates or know which ones truly determine the species distributions. Applying a lasso penalty to help in variable selection may therefore be provide a natural way forward in this context. Finally, a key reality when dealing with presence-only data is the presence of observer bias, in which sampling effort varies throughout the study region. Some models apply a correction for observer bias in the prediction (Hefley *et al.*, 2013; Lahoz-Monfort *et al.*, 2014; Warton *et al.*, 2013) and our tools would be able to accommodate such improvements.

## 7 Conclusion

The new algorithms presented in this article aim to reclassify observations that have uncertain or unknown labels in order to better predict point pattern distributions. We showed that machine learning based models performed better in a general context than mixture based models no matter the initialization method and also better than the individual PPM method that does not include the points with unknown labels. Our simulations showed encouraging results in this context with good performances in some cases, although there are some improvements to implement in order to make the tools more appropriate for real life data.

## Acknowledgments

Computational resources used in this work were provided by Intersect Australia Ltd.

## Authors’ contributions

EG and IR conceived the ideas and designed methodology; EG and IR built the algorithms; EG analyzed the data; EG and IR led the writing of the manuscript. MM had an overview on the project. MM and EB reviewed the paper. All authors contributed critically to the drafts and gave final approval for publication.

## Data accessibility

Rscript: An example of the scripts used for this paper is available here: functions to use for the test simulation 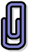 and the script example 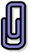.

