## Supplementary files for "Classification of unlabeled observations in Species Distribution Modelling using Point Process Models"

### Appendix

#### 0.1 Complementary plots and results

##### 0.1.1 Tables of performances results

Table 1: Table of prediction performances for all initialization method for the parameters of abundances and correlation: $m1$=32, $m2$=42, $m3$=23; $\rho_{1-2}$=0.85, $\rho_{1-3}$=-0.09, $\rho_{2-3}$=0.20

| | **IMSE20%** | | | | | | | |
|---|---|---|---|---|---|---|---|---|
| | **knn** | **kmeans** | **rand** | **equal** | **indiv** | **norm** | **LoopgrW** | **LoophgW** |
| **mean** | 29873.34 | 30566.7 | 30103.22 | 30823.04 | 30833.03 | 33074.85 | 28473.42 | 38337.59 |
| **median** | 29145.41 | 29401.21 | 29155.54 | 29835.43 | 29747.7 | 32574.67 | 28615.05 | 37671.94 |
| **1stQ** | 26401.45 | 26423.53 | 26397.54 | 26632.7 | 26445.89 | 21545.22 | 25148.72 | 28099.95 |
| **3rdQ** | 29873.34 | 30566.7 | 30103.22 | 30823.04 | 30833.03 | 33074.85 | 28473.42 | 38337.59 |
| | **IMSE50%** | | | | | | | |
| | **knn** | **kmeans** | **rand** | **equal** | **indiv** | **norm** | **LoopgrW** | **LoophgW** |
| **mean** | 47632.35 | 52664.62 | 49666.52 | 84269.4 | 45205.57 | 55993.15 | 34261.84 | 57415.27 |
| **median** | 44845.45 | 47428.91 | 45379.22 | 82418.2 | 42026.26 | 56620.63 | 33132.19 | 58286.63 |
| **1stQ** | 37443.57 | 39007.72 | 37242.49 | 67830.07 | 33147.09 | 49071.91 | 28818.39 | 49986.7 |
| **3rdQ** | 47632.35 | 52664.62 | 49666.52 | 84269.4 | 45205.57 | 55993.15 | 34261.84 | 57415.27 |
| | **IMSE80%** | | | | | | | |
| | **knn** | **kmeans** | **rand** | **equal** | **indiv** | **norm** | **LoopgrW** | **LoophgW** |
| **mean** | 141280.5 | 143874.3 | 149054.2 | 185543.2 | 101548.9 | 67697.39 | 77118.95 | 76162.98 |
| **median** | 133185.2 | 136960.7 | 139548.5 | 179973.8 | 94673.27 | 68406.58 | 65069.8 | 77020.24 |
| **1stQ** | 104787.6 | 106049.5 | 111690.5 | 143565.1 | 63646.31 | 63958.99 | 46908.24 | 71942.22 |
| **3rdQ** | 141280.5 | 143874.3 | 149054.2 | 185543.2 | 101548.9 | 67697.39 | 77118.95 | 76162.98 |
| | **sumcor20%** | | | | | | | |
| | **knn** | **kmeans** | **rand** | **equal** | **indiv** | **norm** | **LoopgrW** | **LoophgW** |
| **mean** | 2.851039 | 2.847275 | 2.849895 | 2.847591 | 2.838309 | 2.768368 | 2.869781 | 2.768368 |
| **median** | 2.853988 | 2.853231 | 2.853619 | 2.852552 | 2.842891 | 2.773598 | 2.871977 | 2.773598 |
| **1stQ** | 2.836368 | 2.834174 | 2.836492 | 2.833477 | 2.81903 | 2.704882 | 2.853768 | 2.704882 |
| **3rdQ** | 2.851039 | 2.847275 | 2.849895 | 2.847591 | 2.838309 | 2.768368 | 2.869781 | 2.768368 |
| | **sumcor50%** | | | | | | | |
| | **knn** | **kmeans** | **rand** | **equal** | **indiv** | **norm** | **LoopgrW** | **LoophgW** |
| **mean** | 2.781201 | 2.752753 | 2.770245 | 2.681062 | 2.742776 | 2.596904 | 2.844039 | 2.596904 |
| **median** | 2.795817 | 2.780585 | 2.78986 | 2.691907 | 2.760229 | 2.595641 | 2.855542 | 2.595641 |
| **1stQ** | 2.753207 | 2.718186 | 2.737616 | 2.629421 | 2.69461 | 2.546021 | 2.83245 | 2.546021 |
| **3rdQ** | 2.781201 | 2.752753 | 2.770245 | 2.681062 | 2.742776 | 2.596904 | 2.844039 | 2.596904 |
| | **sumcor80%** | | | | | | | |
| | **knn** | **kmeans** | **rand** | **equal** | **indiv** | **norm** | **LoopgrW** | **LoophgW** |
| **mean** | 2.440887 | 2.432054 | 2.43764 | 2.375754 | 2.418456 | 2.509087 | 2.639854 | 2.509087 |
| **median** | 2.458626 | 2.462068 | 2.464771 | 2.381614 | 2.431692 | 2.506043 | 2.703239 | 2.506043 |
| **1stQ** | 2.324382 | 2.318554 | 2.322671 | 2.248885 | 2.289263 | 2.477674 | 2.545126 | 2.477674 |
| **3rdQ** | 2.440887 | 2.432054 | 2.43764 | 2.375754 | 2.418456 | 2.509087 | 2.639854 | 2.509087 |

Table 2: Table of relabeling performances for all initialization method for the parameters of abundances and correlation: $m1$=32, $m2$=42, $m3$=23; $\rho_{1-2}$=0.85, $\rho_{1-3}$=-0.09, $\rho_{2-3}$=0.20

| | **MeanRSS20%** | | | | | | | |
|---|---|---|---|---|---|---|---|---|
| | **knn** | **kmeans** | **rand** | **equal** | **indiv** | **norm** | **LoopgrW** | **LoophgW** |
| **mean** | 0.345918 | 0.347046 | 0.345945 | 0.347217 | 0.317614 | 0.367536 | 0.314454 | 0.367536 |
| **median** | 0.344955 | 0.345226 | 0.345374 | 0.345525 | 0.315347 | 0.365854 | 0.31309 | 0.365854 |
| **1stQ** | 0.316702 | 0.317593 | 0.316894 | 0.317426 | 0.291636 | 0.354399 | 0.289945 | 0.354399 |
| **3rdQ** | 0.345918 | 0.347046 | 0.345945 | 0.347217 | 0.317614 | 0.367536 | 0.314454 | 0.367536 |
| | **MeanRSS50%** | | | | | | | |
| | **knn** | **kmeans** | **rand** | **equal** | **indiv** | **norm** | **LoopgrW** | **LoophgW** |
| **mean** | 0.453568 | 0.451534 | 0.454157 | 0.531491 | 0.327572 | 0.408077 | 0.318699 | 0.408077 |
| **median** | 0.457291 | 0.458208 | 0.460002 | 0.530489 | 0.323325 | 0.407274 | 0.315582 | 0.407274 |
| **1stQ** | 0.433407 | 0.427711 | 0.43012 | 0.523974 | 0.309401 | 0.405133 | 0.302372 | 0.405133 |
| **3rdQ** | 0.453568 | 0.451534 | 0.454157 | 0.531491 | 0.327572 | 0.408077 | 0.318699 | 0.408077 |
| | **MeanRSS80%** | | | | | | | |
| | **knn** | **kmeans** | **rand** | **equal** | **indiv** | **norm** | **LoopgrW** | **LoophgW** |
| **mean** | 0.55362 | 0.598744 | 0.59873 | 0.656055 | 0.365143 | 0.431779 | 0.365226 | 0.431779 |
| **median** | 0.556142 | 0.557167 | 0.55731 | 0.656033 | 0.358044 | 0.431575 | 0.357293 | 0.431575 |
| **1stQ** | 0.554679 | 0.555066 | 0.555695 | 0.65512 | 0.333673 | 0.429713 | 0.331644 | 0.429713 |
| **3rdQ** | 0.55362 | 0.598744 | 0.59873 | 0.656055 | 0.365143 | 0.431779 | 0.365226 | 0.431779 |
| | **accmat50%** | | | | | | | |
| | **knn** | **kmeans** | **rand** | **equal** | **indiv** | **norm** | **LoopgrW** | **LoophgW** |
| **mean** | 0.495435 | 0.490333 | 0.496604 | 0.4735 | 0.4785 | 0.473889 | 0.472111 | 0.473889 |
| **median** | 0.5 | 0.5 | 0.5 | 0.5 | 0.5 | 0.5 | 0.444444 | 0.5 |
| **1stQ** | 0.444444 | 0.444444 | 0.444444 | 0.444444 | 0.388889 | 0.388889 | 0.388889 | 0.388889 |
| **3rdQ** | 0.495435 | 0.490333 | 0.496604 | 0.4735 | 0.4785 | 0.473889 | 0.472111 | 0.473889 |
| | **accmatmat50%** | | | | | | | |
| | **knn** | **kmeans** | **rand** | **equal** | **indiv** | **norm** | **LoopgrW** | **LoophgW** |
| **mean** | 0.468229 | 0.454688 | 0.459731 | 0.335 | 0.476417 | 0.472896 | 0.470563 | 0.472896 |
| **median** | 0.458333 | 0.458333 | 0.458333 | 0.333333 | 0.479167 | 0.479167 | 0.479167 | 0.479167 |
| **1stQ** | 0.4375 | 0.4375 | 0.4375 | 0.333333 | 0.416667 | 0.4375 | 0.4375 | 0.4375 |
| **3rdQ** | 0.468229 | 0.454688 | 0.459731 | 0.335 | 0.476417 | 0.472896 | 0.470563 | 0.472896 |
| | **accmatmat80%** | | | | | | | |
| | **knn** | **kmeans** | **rand** | **equal** | **indiv** | **norm** | **LoopgrW** | **LoophgW** |
| **mean** | 0.426829 | 0.377132 | 0.384316 | 0.328947 | 0.466566 | 0.462579 | 0.45254 | 0.462579 |
| **median** | 0.434211 | 0.434211 | 0.434211 | 0.328947 | 0.460526 | 0.460526 | 0.460526 | 0.460526 |
| **1stQ** | 0.434211 | 0.328947 | 0.328947 | 0.328947 | 0.434211 | 0.434211 | 0.421053 | 0.434211 |
| **3rdQ** | 0.468229 | 0.454688 | 0.459731 | 0.335 | 0.476417 | 0.472896 | 0.470563 | 0.472896 |

Table 3: Table of prediction performances for all initialization method for the parameters of abundances and correlation: $m1$=33, $m2$=34, $m3$=35; $\rho_{1-2}$=0.85, $\rho_{1-3}$=-0.09, $\rho_{2-3}$=0.20

| | **IMSE20%** | | | | | | | |
|---|---|---|---|---|---|---|---|---|
| | **knn** | **kmeans** | **rand** | **equal** | **indiv** | **norm** | **LoopgrW** | **LoophgW** |
| **mean** | 31682.4 | 31346.53 | 31211.71 | 28426.21 | 30694.83 | 48906.3 | 29360.53 | 48817.55 |
| **median** | 30702.62 | 30584.4 | 30201.1 | 26726.8 | 30349.27 | 48909.52 | 28870.59 | 48310.83 |
| **1stQ** | 25309.37 | 24285.81 | 24653.1 | 22663.77 | 26983.84 | 36958.04 | 23936.83 | 38132.17 |
| **3rdQ** | 31682.4 | 31346.53 | 31211.71 | 28426.21 | 30694.83 | 48906.3 | 29360.53 | 48817.55 |
| | **IMSE50%** | | | | | | | |
| | **knn** | **kmeans** | **rand** | **equal** | **indiv** | **norm** | **LoopgrW** | **LoophgW** |
| **mean** | 48782.6 | 49806.09 | 48825.12 | 105224.5 | 44405.13 | 67337.22 | 35681.78 | 72785.85 |
| **median** | 45806.18 | 44858.48 | 45040.76 | 103853.1 | 42320.32 | 68005.87 | 33767.29 | 73643.06 |
| **1stQ** | 34346.74 | 33825.66 | 34322.11 | 88807.51 | 28867.55 | 60736.17 | 24564.07 | 65686.65 |
| **3rdQ** | 48782.6 | 49806.09 | 48825.12 | 105224.5 | 44405.13 | 67337.22 | 35681.78 | 72785.85 |
| | **IMSE80%** | | | | | | | |
| | **knn** | **kmeans** | **rand** | **equal** | **indiv** | **norm** | **LoopgrW** | **LoophgW** |
| **mean** | 148675.2 | 147356.9 | 153304.7 | 213997.7 | 96950.5 | 79956.29 | 75765.04 | 98339 |
| **median** | 141853.2 | 135621.7 | 139960 | 208409 | 89415.14 | 80621.03 | 66244.53 | 99143.27 |
| **1stQ** | 105626.9 | 105765.3 | 105988.2 | 181051.7 | 62491.77 | 76432.88 | 45475.13 | 94120.4 |
| **3rdQ** | 148675.2 | 147356.9 | 153304.7 | 213997.7 | 96950.5 | 79956.29 | 75765.04 | 98339 |
| | **sumcor20%** | | | | | | | |
| | **knn** | **kmeans** | **rand** | **equal** | **indiv** | **norm** | **LoopgrW** | **LoophgW** |
| **mean** | 2.831651 | 2.836024 | 2.835977 | 2.864472 | 2.839335 | 2.724069 | 2.845501 | 2.724069 |
| **median** | 2.836267 | 2.840637 | 2.840069 | 2.874718 | 2.844051 | 2.73329 | 2.847182 | 2.73329 |
| **1stQ** | 2.811813 | 2.813422 | 2.816095 | 2.84882 | 2.819519 | 2.661911 | 2.822507 | 2.661911 |
| **3rdQ** | 2.831651 | 2.836024 | 2.835977 | 2.864472 | 2.839335 | 2.724069 | 2.845501 | 2.724069 |
| | **sumcor50%** | | | | | | | |
| | **knn** | **kmeans** | **rand** | **equal** | **indiv** | **norm** | **LoopgrW** | **LoophgW** |
| **mean** | 2.755617 | 2.759118 | 2.763256 | 2.628138 | 2.75412 | 2.595426 | 2.807155 | 2.595426 |
| **median** | 2.761336 | 2.777066 | 2.776934 | 2.641407 | 2.767454 | 2.596093 | 2.81608 | 2.596093 |
| **1stQ** | 2.70087 | 2.707017 | 2.709738 | 2.567773 | 2.697019 | 2.550084 | 2.765208 | 2.550084 |
| **3rdQ** | 2.755617 | 2.759118 | 2.763256 | 2.628138 | 2.75412 | 2.595426 | 2.807155 | 2.595426 |
| | **sumcor80%** | | | | | | | |
| | **knn** | **kmeans** | **rand** | **equal** | **indiv** | **norm** | **LoopgrW** | **LoophgW** |
| **mean** | 2.450128 | 2.444575 | 2.443424 | 2.321052 | 2.432805 | 2.509418 | 2.621491 | 2.509418 |
| **median** | 2.473658 | 2.482951 | 2.467813 | 2.345725 | 2.455511 | 2.507491 | 2.666864 | 2.507491 |
| **1stQ** | 2.325162 | 2.323663 | 2.32243 | 2.210516 | 2.304821 | 2.482758 | 2.522914 | 2.482758 |
| **3rdQ** | 2.450128 | 2.444575 | 2.443424 | 2.321052 | 2.432805 | 2.509418 | 2.621491 | 2.509418 |

Table 4: Table of relabeling performances for all initialization method for the parameters of abundances and correlation: $m1$=33, $m2$=34, $m3$=35; $\rho_{1-2}$=0.85, $\rho_{1-3}$=-0.09, $\rho_{2-3}$=0.20

| | | | | | | | | |
|---|---|---|---|---|---|---|---|---|
| | **MeanRSS20%** | | | | | | | |
| | **knn** | **kmeans** | **rand** | **equal** | **indiv** | **norm** | **LoopgrW** | **LoophgW** |
| **mean** | 0.360506 | 0.36127 | 0.360878 | 0.358587 | 0.347175 | 0.368577 | 0.349602 | 0.368577 |
| **median** | 0.355998 | 0.355642 | 0.355494 | 0.355674 | 0.343864 | 0.366394 | 0.346842 | 0.366394 |
| **1stQ** | 0.324523 | 0.324301 | 0.32438 | 0.323655 | 0.315914 | 0.35173 | 0.321349 | 0.35173 |
| **3rdQ** | 0.360506 | 0.36127 | 0.360878 | 0.358587 | 0.347175 | 0.368577 | 0.349602 | 0.368577 |
| | **MeanRSS50%** | | | | | | | |
| | **knn** | **kmeans** | **rand** | **equal** | **indiv** | **norm** | **LoopgrW** | **LoophgW** |
| **mean** | 0.430069 | 0.444387 | 0.437955 | 0.542182 | 0.356437 | 0.405863 | 0.358256 | 0.405863 |
| **median** | 0.423907 | 0.434345 | 0.431265 | 0.540735 | 0.353543 | 0.405174 | 0.356349 | 0.405174 |
| **1stQ** | 0.395743 | 0.403983 | 0.401017 | 0.53417 | 0.339472 | 0.403063 | 0.344013 | 0.403063 |
| **3rdQ** | 0.430069 | 0.444387 | 0.437955 | 0.542182 | 0.356437 | 0.405863 | 0.358256 | 0.405863 |
| | **MeanRSS80%** | | | | | | | |
| | **knn** | **kmeans** | **rand** | **equal** | **indiv** | **norm** | **LoopgrW** | **LoophgW** |
| **mean** | 0.615044 | 0.631663 | 0.640718 | 0.665938 | 0.38356 | 0.431685 | 0.394776 | 0.431685 |
| **median** | 0.65242 | 0.651876 | 0.653191 | 0.665934 | 0.375222 | 0.431581 | 0.387812 | 0.431581 |
| **1stQ** | 0.580872 | 0.642941 | 0.643302 | 0.664801 | 0.356218 | 0.42946 | 0.364026 | 0.42946 |
| **3rdQ** | 0.615044 | 0.631663 | 0.640718 | 0.665938 | 0.38356 | 0.431685 | 0.394776 | 0.431685 |
| | **accmat20%** | | | | | | | |
| | **knn** | **kmeans** | **rand** | **equal** | **indiv** | **norm** | **LoopgrW** | **LoophgW** |
| **mean** | 0.488316 | 0.486499 | 0.490211 | 0.471261 | 0.468263 | 0.497316 | 0.472895 | 0.497316 |
| **median** | 0.473684 | 0.473684 | 0.473684 | 0.473684 | 0.473684 | 0.473684 | 0.473684 | 0.473684 |
| **1stQ** | 0.421053 | 0.421053 | 0.421053 | 0.421053 | 0.421053 | 0.421053 | 0.421053 | 0.421053 |
| **3rdQ** | 0.488316 | 0.486499 | 0.490211 | 0.471261 | 0.468263 | 0.497316 | 0.472895 | 0.497316 |
| | **accmat50%** | | | | | | | |
| | **knn** | **kmeans** | **rand** | **equal** | **indiv** | **norm** | **LoopgrW** | **LoophgW** |
| **mean** | 0.47152 | 0.455291 | 0.46362 | 0.32334 | 0.47944 | 0.49316 | 0.47916 | 0.49316 |
| **median** | 0.48 | 0.46 | 0.48 | 0.32 | 0.48 | 0.5 | 0.48 | 0.5 |
| **1stQ** | 0.44 | 0.42 | 0.42 | 0.32 | 0.44 | 0.46 | 0.44 | 0.46 |
| **3rdQ** | 0.47152 | 0.455291 | 0.46362 | 0.32334 | 0.47944 | 0.49316 | 0.47916 | 0.49316 |
| | **accmat80%** | | | | | | | |
| | **knn** | **kmeans** | **rand** | **equal** | **indiv** | **norm** | **LoopgrW** | **LoophgW** |
| **mean** | 0.361803 | 0.347469 | 0.342012 | 0.320988 | 0.47384 | 0.482877 | 0.470926 | 0.482877 |
| **median** | 0.333333 | 0.333333 | 0.333333 | 0.320988 | 0.481482 | 0.481482 | 0.481482 | 0.481482 |
| **1stQ** | 0.320988 | 0.320988 | 0.320988 | 0.320988 | 0.444444 | 0.45679 | 0.444444 | 0.45679 |
| **3rdQ** | 0.361803 | 0.347469 | 0.342012 | 0.320988 | 0.47384 | 0.482877 | 0.470926 | 0.482877 |

Table 5: Table of prediction performances for all initialization method for the parameters of abundances and correlation: $m1$=42, $m2$=31 $m3$=25; $\rho_{1-2}$=0.09, $\rho_{1-3}$=-0.42, $\rho_{2-3}$=0.20

| | **IMSE20%** | | | | | | | |
|---|---|---|---|---|---|---|---|---|
| | **knn** | **kmeans** | **rand** | **equal** | **indiv** | **norm** | **LoopgrW** | **LoophgW** |
| **mean** | 26177.01 | 26983.28 | 26592.93 | 28090.51 | 18937.2 | 29416.79 | 21404.66 | 25017.85 |
| **median** | 24579.6 | 25331.95 | 24880.87 | 26483.28 | 17916.56 | 28371.03 | 20694.13 | 23614.66 |
| **1stQ** | 21478.55 | 21939.67 | 21740.9 | 22659.48 | 14100.65 | 20408.61 | 18585.08 | 20564.74 |
| **3rdQ** | 26177.01 | 26983.28 | 26592.93 | 28090.51 | 18937.2 | 29416.79 | 21404.66 | 25017.85 |
| | **IMSE50%** | | | | | | | |
| | **knn** | **kmeans** | **rand** | **equal** | **indiv** | **norm** | **LoopgrW** | **LoophgW** |
| **mean** | 57807.34 | 64296.63 | 60948.44 | 81001.52 | 35471.48 | 53828.55 | 33905.75 | 43210.86 |
| **median** | 55372.88 | 60847.89 | 57639.76 | 79095.56 | 32584.02 | 54051.5 | 32548.78 | 41347.14 |
| **1stQ** | 45832.9 | 48870.65 | 47640.62 | 65799.91 | 24250.53 | 47145.08 | 28726.78 | 34239.99 |
| **3rdQ** | 57807.34 | 64296.63 | 60948.44 | 81001.52 | 35471.48 | 53828.55 | 33905.75 | 43210.86 |
| | **IMSE80%** | | | | | | | |
| | **knn** | **kmeans** | **rand** | **equal** | **indiv** | **norm** | **LoopgrW** | **LoophgW** |
| **mean** | 189799.2 | 182154.1 | 182646.4 | 195116.4 | 106020.5 | 68313.49 | 76166.26 | 106692.4 |
| **median** | 185000.4 | 178741.6 | 179340.2 | 188804.8 | 97995 | 68499.31 | 64749.66 | 94257.43 |
| **1stQ** | 152182.1 | 143859.2 | 144231.6 | 160146.4 | 67785.04 | 64578.99 | 48636.5 | 80049.25 |
| **3rdQ** | 189799.2 | 182154.1 | 182646.4 | 195116.4 | 106020.5 | 68313.49 | 76166.26 | 106692.4 |
| | **sumcor20%** | | | | | | | |
| | **knn** | **kmeans** | **rand** | **equal** | **indiv** | **norm** | **LoopgrW** | **LoophgW** |
| **mean** | 2.881285 | 2.876016 | 2.879242 | 2.874181 | 2.874752 | 2.718791 | 2.854127 | 2.877883 |
| **median** | 2.887516 | 2.884683 | 2.886344 | 2.880487 | 2.879329 | 2.741419 | 2.856694 | 2.883765 |
| **1stQ** | 2.868792 | 2.863323 | 2.865718 | 2.858865 | 2.854883 | 2.642441 | 2.842936 | 2.865685 |
| **3rdQ** | 2.881285 | 2.876016 | 2.879242 | 2.874181 | 2.874752 | 2.718791 | 2.854127 | 2.877883 |
| | **sumcor50%** | | | | | | | |
| | **knn** | **kmeans** | **rand** | **equal** | **indiv** | **norm** | **LoopgrW** | **LoophgW** |
| **mean** | 2.7594 | 2.717874 | 2.73986 | 2.708711 | 2.770196 | 2.467983 | 2.772992 | 2.77945 |
| **median** | 2.773326 | 2.742529 | 2.762842 | 2.715311 | 2.776119 | 2.467739 | 2.784116 | 2.789557 |
| **1stQ** | 2.716048 | 2.665272 | 2.695086 | 2.662646 | 2.725504 | 2.39264 | 2.75288 | 2.754554 |
| **3rdQ** | 2.7594 | 2.717874 | 2.73986 | 2.708711 | 2.770196 | 2.467983 | 2.772992 | 2.77945 |
| | **sumcor80%** | | | | | | | |
| | **knn** | **kmeans** | **rand** | **equal** | **indiv** | **norm** | **LoopgrW** | **LoophgW** |
| **mean** | 2.380468 | 2.358902 | 2.375611 | 2.373889 | 2.404616 | 2.306063 | 2.565392 | 2.481949 |
| **median** | 2.391397 | 2.379346 | 2.386725 | 2.386999 | 2.414627 | 2.301897 | 2.622205 | 2.575522 |
| **1stQ** | 2.267055 | 2.249227 | 2.261969 | 2.25445 | 2.286175 | 2.256923 | 2.453887 | 2.350692 |
| **3rdQ** | 2.380468 | 2.358902 | 2.375611 | 2.373889 | 2.404616 | 2.306063 | 2.565392 | 2.481949 |

Table 6: Table of relabeling performances for all initialization method for the parameters of abundances and correlation: $m1$=42, $m2$=31, $m3$=25; $\rho_{1-2}$=0.09, $\rho_{1-3}$=-0.42, $\rho_{2-3}$=0.20

| | **MeanRSS20%** | | | | | | | |
|---|---|---|---|---|---|---|---|---|
| | **knn** | **kmeans** | **rand** | **equal** | **indiv** | **norm** | **LoopgrW** | **LoophgW** |
| **mean** | 0.289192 | 0.291884 | 0.290793 | 0.299145 | 0.279257 | 0.324109 | 0.287092 | 0.27127 |
| **median** | 0.286336 | 0.28885 | 0.288383 | 0.297714 | 0.276741 | 0.321944 | 0.284232 | 0.268738 |
| **1stQ** | 0.251579 | 0.256319 | 0.253501 | 0.263332 | 0.245303 | 0.305604 | 0.255509 | 0.234031 |
| **3rdQ** | 0.289192 | 0.291884 | 0.290793 | 0.299145 | 0.279257 | 0.324109 | 0.287092 | 0.27127 |
| | **MeanRSS50%** | | | | | | | |
| | **knn** | **kmeans** | **rand** | **equal** | **indiv** | **norm** | **LoopgrW** | **LoophgW** |
| **mean** | 0.353492 | 0.384262 | 0.374061 | 0.471672 | 0.287333 | 0.382842 | 0.298988 | 0.282575 |
| **median** | 0.340381 | 0.38215 | 0.370224 | 0.470474 | 0.285447 | 0.382285 | 0.298218 | 0.280772 |
| **1stQ** | 0.303968 | 0.329134 | 0.321185 | 0.465662 | 0.268877 | 0.379657 | 0.284445 | 0.257346 |
| **3rdQ** | 0.353492 | 0.384262 | 0.374061 | 0.471672 | 0.287333 | 0.382842 | 0.298988 | 0.282575 |
| | **MeanRSS80%** | | | | | | | |
| | **knn** | **kmeans** | **rand** | **equal** | **indiv** | **norm** | **LoopgrW** | **LoophgW** |
| **mean** | 0.562753 | 0.6101 | 0.602795 | 0.56176 | 0.321531 | 0.422909 | 0.326881 | 0.330608 |
| **median** | 0.561774 | 0.564835 | 0.562819 | 0.561701 | 0.316331 | 0.42272 | 0.317651 | 0.310806 |
| **1stQ** | 0.560625 | 0.561526 | 0.561261 | 0.560917 | 0.29715 | 0.420528 | 0.300258 | 0.28666 |
| **3rdQ** | 0.562753 | 0.6101 | 0.602795 | 0.56176 | 0.321531 | 0.422909 | 0.326881 | 0.330608 |
| | **accmat20%** | | | | | | | |
| | **knn** | **kmeans** | **rand** | **equal** | **indiv** | **norm** | **LoopgrW** | **LoophgW** |
| **mean** | 0.630842 | 0.62279 | 0.624895 | 0.60176 | 0.647579 | 0.655895 | 0.650947 | 0.657895 |
| **median** | 0.631579 | 0.631579 | 0.631579 | 0.578947 | 0.631579 | 0.684211 | 0.631579 | 0.684211 |
| **1stQ** | 0.578947 | 0.578947 | 0.578947 | 0.526316 | 0.578947 | 0.578947 | 0.578947 | 0.578947 |
| **3rdQ** | 0.630842 | 0.62279 | 0.624895 | 0.60176 | 0.647579 | 0.655895 | 0.650947 | 0.657895 |
| | **accmat50%** | | | | | | | |
| | **knn** | **kmeans** | **rand** | **equal** | **indiv** | **norm** | **LoopgrW** | **LoophgW** |
| **mean** | 0.566563 | 0.524608 | 0.540958 | 0.437667 | 0.634167 | 0.643688 | 0.633896 | 0.647563 |
| **median** | 0.583333 | 0.520833 | 0.541667 | 0.4375 | 0.625 | 0.645833 | 0.625 | 0.645833 |
| **1stQ** | 0.5 | 0.458333 | 0.479167 | 0.4375 | 0.604167 | 0.604167 | 0.604167 | 0.604167 |
| **3rdQ** | 0.566563 | 0.524608 | 0.540958 | 0.437667 | 0.634167 | 0.643688 | 0.633896 | 0.647563 |
| | **accmat80%** | | | | | | | |
| | **knn** | **kmeans** | **rand** | **equal** | **indiv** | **norm** | **LoopgrW** | **LoophgW** |
| **mean** | 0.41987 | 0.368546 | 0.378961 | 0.428571 | 0.588727 | 0.59987 | 0.577662 | 0.596987 |
| **median** | 0.428571 | 0.376623 | 0.428571 | 0.428571 | 0.597403 | 0.61039 | 0.584416 | 0.623377 |
| **1stQ** | 0.428571 | 0.311688 | 0.311688 | 0.428571 | 0.558442 | 0.571429 | 0.545455 | 0.558442 |
| **3rdQ** | 0.41987 | 0.368546 | 0.378961 | 0.428571 | 0.588727 | 0.59987 | 0.577662 | 0.596987 |

Table 7: Table of prediction performances for all initialization method for the parameters of abundances and correlation: $m1$=39, $m2$=37, $m3$=38; $\rho_{1-2}$=0.09, $\rho_{1-3}$=-0.42, $\rho_{2-3}$=0.20

| | **IMSE20%** | | | | | | | |
|---|---|---|---|---|---|---|---|---|
| | **knn** | **kmeans** | **rand** | **equal** | **indiv** | **norm** | **LoopgrW** | **LoophgW** |
| **mean** | 23262.15 | 22927.19 | 22581.12 | 23803.51 | 23084.38 | 42860.15 | 24887.36 | 36467.82 |
| **median** | 20869.05 | 20597.34 | 20454.91 | 21125.99 | 21818.36 | 42615.25 | 23549.12 | 36353.09 |
| **1stQ** | 18281.93 | 17993.35 | 18022.88 | 18409.4 | 18546.44 | 34983.66 | 19276.34 | 29762 |
| **3rdQ** | 23262.15 | 22927.19 | 22581.12 | 23803.51 | 23084.38 | 42860.15 | 24887.36 | 36467.82 |
| | **IMSE50%** | | | | | | | |
| | **knn** | **kmeans** | **rand** | **equal** | **indiv** | **norm** | **LoopgrW** | **LoophgW** |
| **mean** | 45582.75 | 48804.23 | 44512.35 | 83619.05 | 37438.17 | 69556.75 | 33132.95 | 79528.63 |
| **median** | 43251.61 | 46083.1 | 41657.62 | 80243.78 | 34651.06 | 69530.29 | 30161.47 | 79592.6 |
| **1stQ** | 32229.8 | 33856.28 | 31113.64 | 69641.78 | 25518.74 | 63080.32 | 23424.88 | 72503.96 |
| **3rdQ** | 45582.75 | 48804.23 | 44512.35 | 83619.05 | 37438.17 | 69556.75 | 33132.95 | 79528.63 |
| | **IMSE80%** | | | | | | | |
| | **knn** | **kmeans** | **rand** | **equal** | **indiv** | **norm** | **LoopgrW** | **LoophgW** |
| **mean** | 144828.2 | 148873 | 153726.2 | 172657.9 | 95463.53 | 83480.48 | 67576.17 | 111728.3 |
| **median** | 135307.4 | 141527.2 | 148165.5 | 166194 | 87133.37 | 84077.29 | 62222.48 | 112477.4 |
| **1stQ** | 100631.9 | 109052.2 | 110632.6 | 142021.1 | 61959.71 | 79488.58 | 42845.14 | 107127.3 |
| **3rdQ** | 144828.2 | 148873 | 153726.2 | 172657.9 | 95463.53 | 83480.48 | 67576.17 | 111728.3 |
| | **sumcor20%** | | | | | | | |
| | **knn** | **kmeans** | **rand** | **equal** | **indiv** | **norm** | **LoopgrW** | **LoophgW** |
| **mean** | 2.875633 | 2.874768 | 2.877136 | 2.873493 | 2.877069 | 2.786146 | 2.862084 | 2.786146 |
| **median** | 2.884375 | 2.884464 | 2.884818 | 2.883278 | 2.881602 | 2.79256 | 2.869095 | 2.79256 |
| **1stQ** | 2.861326 | 2.863673 | 2.865306 | 2.859006 | 2.854155 | 2.750797 | 2.842232 | 2.750797 |
| **3rdQ** | 2.875633 | 2.874768 | 2.877136 | 2.873493 | 2.877069 | 2.786146 | 2.862084 | 2.786146 |
| | **sumcor50%** | | | | | | | |
| | **knn** | **kmeans** | **rand** | **equal** | **indiv** | **norm** | **LoopgrW** | **LoophgW** |
| **mean** | 2.776699 | 2.738335 | 2.767012 | 2.688479 | 2.779698 | 2.567736 | 2.801945 | 2.567736 |
| **median** | 2.785394 | 2.766723 | 2.784681 | 2.697484 | 2.786913 | 2.567764 | 2.814532 | 2.567764 |
| **1stQ** | 2.734451 | 2.680041 | 2.722882 | 2.646931 | 2.729348 | 2.533011 | 2.763007 | 2.533011 |
| **3rdQ** | 2.776699 | 2.738335 | 2.767012 | 2.688479 | 2.779698 | 2.567736 | 2.801945 | 2.567736 |
| | **sumcor80%** | | | | | | | |
| | **knn** | **kmeans** | **rand** | **equal** | **indiv** | **norm** | **LoopgrW** | **LoophgW** |
| **mean** | 2.487917 | 2.410227 | 2.451776 | 2.392965 | 2.45751 | 2.402248 | 2.633134 | 2.402248 |
| **median** | 2.507393 | 2.45376 | 2.470099 | 2.39809 | 2.474588 | 2.398593 | 2.649849 | 2.398593 |
| **1stQ** | 2.378056 | 2.323527 | 2.355395 | 2.311409 | 2.363292 | 2.379373 | 2.547662 | 2.379373 |
| **3rdQ** | 2.487917 | 2.410227 | 2.451776 | 2.392965 | 2.45751 | 2.402248 | 2.633134 | 2.402248 |

Table 8: Table of relabeling performances for all initialization method for the parameters of abundances and correlation: $m1$=39, $m2$=37, $m3$=38; $\rho_{1-2}$=0.09, $\rho_{1-3}$=-0.42, $\rho_{2-3}$=0.20

| | | | | | | | | |
|---|---|---|---|---|---|---|---|---|
| | **MeanRSS20%** | | | | | | | |
| | **knn** | **kmeans** | **rand** | **equal** | **indiv** | **norm** | **LoopgrW** | **LoophgW** |
| **mean** | 0.295202 | 0.29564 | 0.294835 | 0.297617 | 0.290555 | 0.320686 | 0.291255 | 0.320686 |
| **median** | 0.293148 | 0.293974 | 0.29327 | 0.294764 | 0.288679 | 0.318697 | 0.289933 | 0.318697 |
| **1stQ** | 0.261123 | 0.261179 | 0.261172 | 0.263076 | 0.259109 | 0.301153 | 0.262641 | 0.301153 |
| **3rdQ** | 0.295202 | 0.29564 | 0.294835 | 0.297617 | 0.290555 | 0.320686 | 0.291255 | 0.320686 |
| | **MeanRSS50%** | | | | | | | |
| | **knn** | **kmeans** | **rand** | **equal** | **indiv** | **norm** | **LoopgrW** | **LoophgW** |
| **mean** | 0.329141 | 0.362157 | 0.338572 | 0.506527 | 0.296188 | 0.379503 | 0.295784 | 0.379503 |
| **median** | 0.327113 | 0.352799 | 0.336062 | 0.507514 | 0.294482 | 0.37901 | 0.295456 | 0.37901 |
| **1stQ** | 0.305748 | 0.326411 | 0.313927 | 0.502052 | 0.278857 | 0.376903 | 0.280101 | 0.376903 |
| **3rdQ** | 0.329141 | 0.362157 | 0.338572 | 0.506527 | 0.296188 | 0.379503 | 0.295784 | 0.379503 |
| | **MeanRSS80%** | | | | | | | |
| | **knn** | **kmeans** | **rand** | **equal** | **indiv** | **norm** | **LoopgrW** | **LoophgW** |
| **mean** | 0.53173 | 0.593943 | 0.589696 | 0.639538 | 0.321745 | 0.422136 | 0.316495 | 0.422136 |
| **median** | 0.626145 | 0.64276 | 0.641813 | 0.639392 | 0.316651 | 0.421867 | 0.310521 | 0.421867 |
| **1stQ** | 0.407638 | 0.507771 | 0.502303 | 0.637981 | 0.299136 | 0.419555 | 0.295843 | 0.419555 |
| **3rdQ** | 0.53173 | 0.593943 | 0.589696 | 0.639538 | 0.321745 | 0.422136 | 0.316495 | 0.422136 |
| | **accmat20%** | | | | | | | |
| | **knn** | **kmeans** | **rand** | **equal** | **indiv** | **norm** | **LoopgrW** | **LoophgW** |
| **mean** | 0.61 | 0.607191 | 0.609181 | 0.602286 | 0.618429 | 0.609571 | 0.623238 | 0.609571 |
| **median** | 0.619048 | 0.619048 | 0.619048 | 0.619048 | 0.619048 | 0.619048 | 0.619048 | 0.619048 |
| **1stQ** | 0.571429 | 0.571429 | 0.571429 | 0.571429 | 0.571429 | 0.571429 | 0.571429 | 0.571429 |
| **3rdQ** | 0.61 | 0.607191 | 0.609181 | 0.602286 | 0.618429 | 0.609571 | 0.623238 | 0.609571 |
| | **accmat50%** | | | | | | | |
| | **knn** | **kmeans** | **rand** | **equal** | **indiv** | **norm** | **LoopgrW** | **LoophgW** |
| **mean** | 0.586748 | 0.532804 | 0.574071 | 0.343857 | 0.603018 | 0.600482 | 0.603696 | 0.600482 |
| **median** | 0.589286 | 0.553571 | 0.589286 | 0.339286 | 0.607143 | 0.607143 | 0.607143 | 0.607143 |
| **1stQ** | 0.553571 | 0.482143 | 0.535714 | 0.339286 | 0.571429 | 0.571429 | 0.571429 | 0.571429 |
| **3rdQ** | 0.586748 | 0.532804 | 0.574071 | 0.343857 | 0.603018 | 0.600482 | 0.603696 | 0.600482 |
| | **accmat80%** | | | | | | | |
| | **knn** | **kmeans** | **rand** | **equal** | **indiv** | **norm** | **LoopgrW** | **LoophgW** |
| **mean** | 0.4377 | 0.370927 | 0.384311 | 0.344444 | 0.573133 | 0.586611 | 0.570711 | 0.586611 |
| **median** | 0.355556 | 0.333333 | 0.344444 | 0.344444 | 0.577778 | 0.588889 | 0.577778 | 0.588889 |
| **1stQ** | 0.333333 | 0.322222 | 0.322222 | 0.344444 | 0.544444 | 0.566667 | 0.544444 | 0.566667 |
| **3rdQ** | 0.4377 | 0.370927 | 0.384311 | 0.344444 | 0.573133 | 0.586611 | 0.570711 | 0.586611 |

##### 0.1.2 Results for the data varying parameters

In this section, details of the 20 and 50% hidden observation are presented. For most of the combination of abundances and correlations, for low percentage of hidden observation (20%), the methods show consistent result with the initial process and show similar results as the individual method. We can notice that when the distribution are less correlated the prediction are closer to the initial process Figures 5, 5. For 50% of hidden observation, knn method (except for the combination different abundances and two distribution are highly correlated), the Loop gr method displayed better results than the individual method. It can be noted that the Loop gr method tend to slightly overestimate the predictions when distribution are not highly correlated, Figures 3, 4.

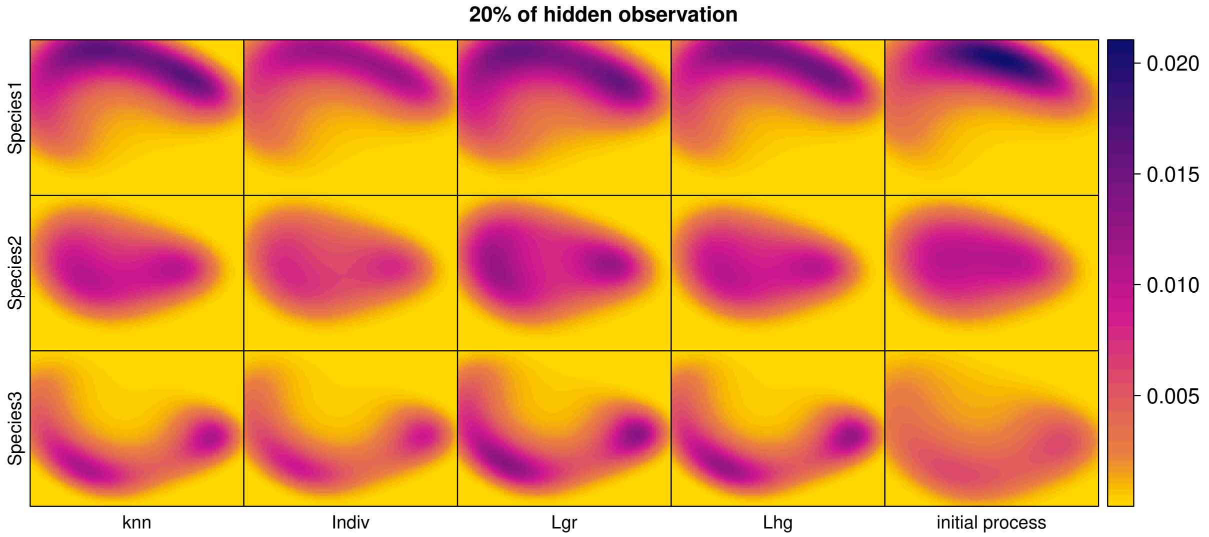


Figure 1: Predicted intensities obtained for the knn, individual, Loop grW and Loop grW initialization methods and the initial intensities from the process at 20% of hidden observations.The parameters of abundances and correlation are: $m1$=32, $m2$=42, $m3$=23; $\rho_{1-2}$=0.85, $\rho_{1-3}$=-0.09, $\rho_{2-3}$=0.20.

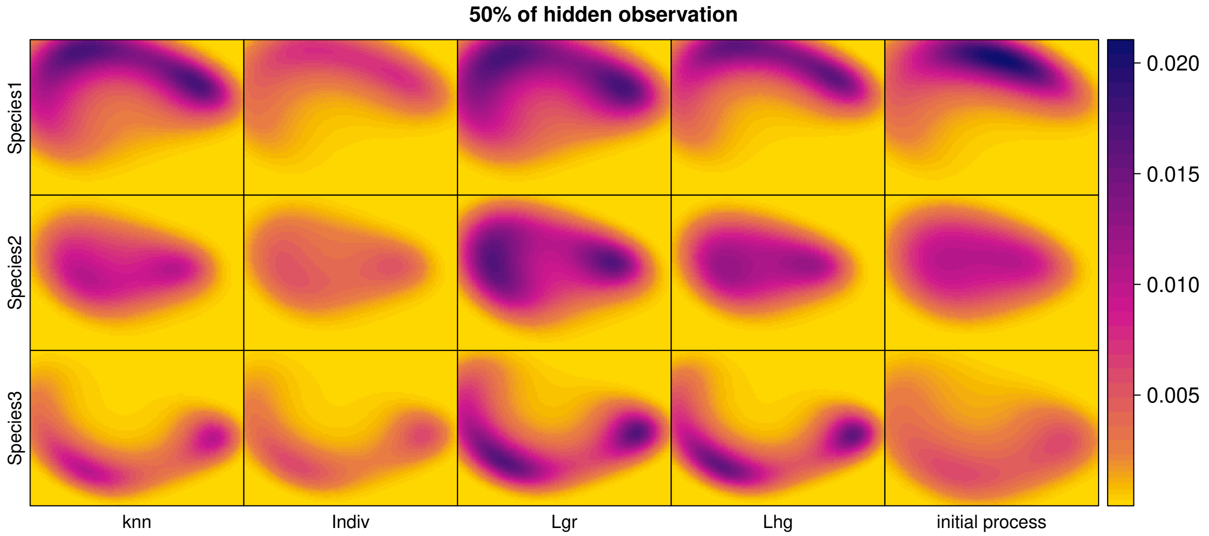


Figure 2: Predicted intensities obtained for the knn, individual, Loop grW and Loop grW initialization methods and the initial intensities from the process at 50% of hidden observations.The parameters of abundances and correlation are: $m1$=32, $m2$=42, $m3$=23; $\rho_{1-2}$=0.85, $\rho_{1-3}$=-0.09, $\rho_{2-3}$=0.20.

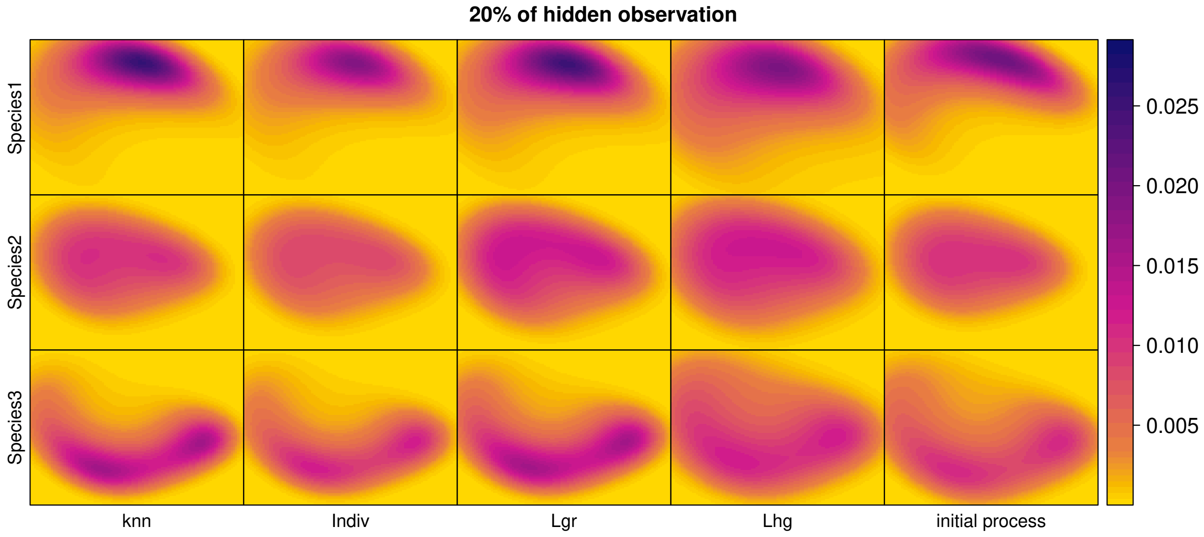


Figure 3: Predicted intensities obtained for the knn, individual, LoopngrW and Loop grW initialization methods and the initial intensities from the process at 20% of hidden observations. The parameters of abundances and correlation are:$m1$=33, $m2$=34, $m3$=35; $\rho_{1-2}$=0.85, $\rho_{1-3}$=-0.09, $\rho_{2-3}$=0.20.

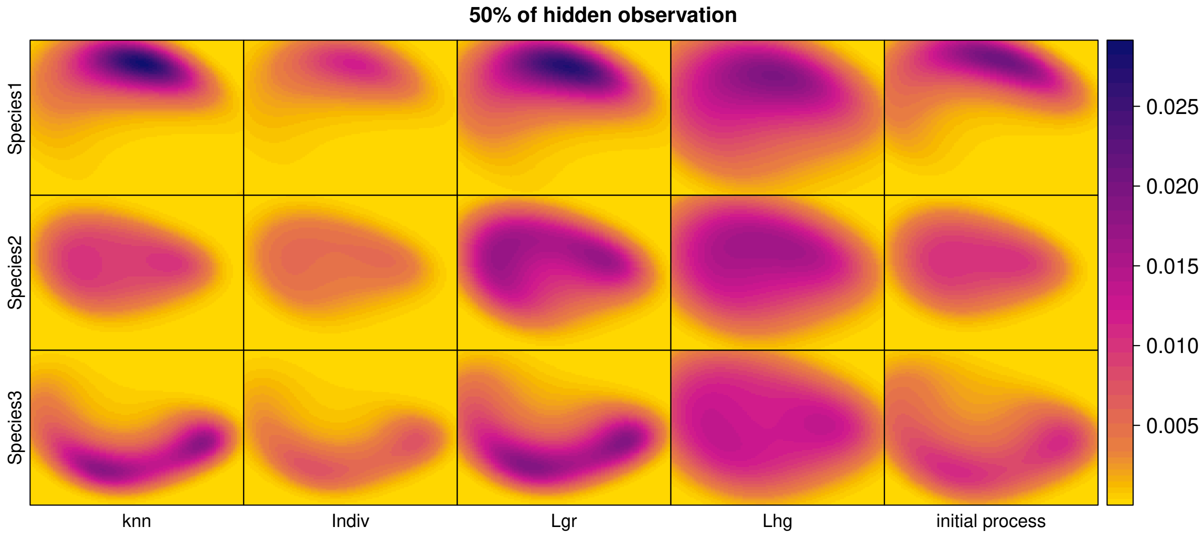


Figure 4: Predicted intensities obtained for the knn, individual, Loop grW and Loop grW initialization methods and the initial intensities from the process at 50% of hidden observations. The parameters of abundances and correlation are: $m1$=33, $m2$=34, $m3$=35; $\rho_{1-2}$=0.85, $\rho_{1-3}$=-0.09, $\rho_{2-3}$=0.20.

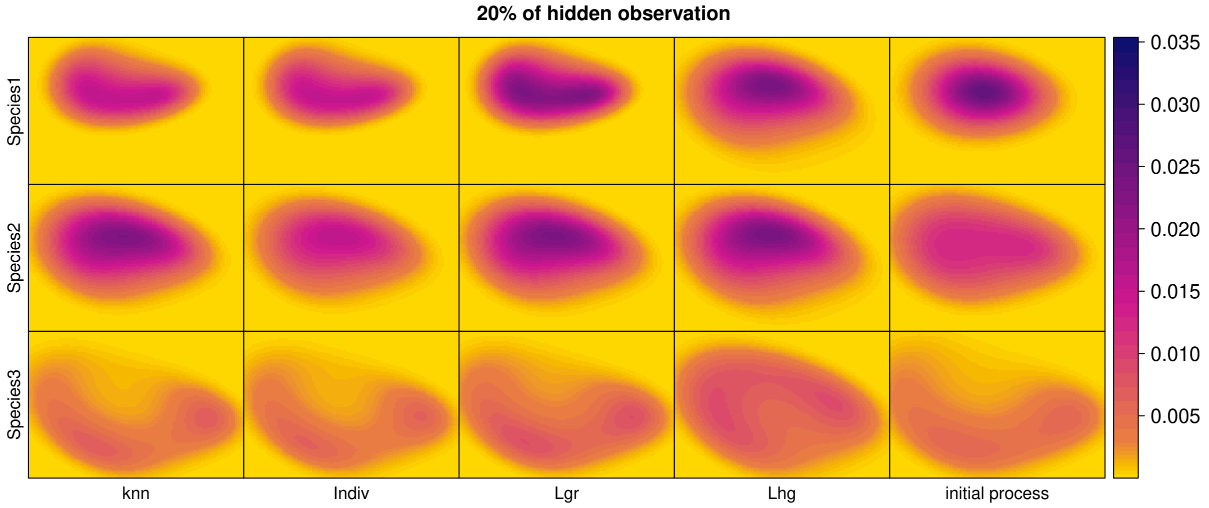


Figure 5: Predicted intensities obtained for the knn, individual, Loop grW and Loop grW initialization methods and the initial intensities from the process at 20% of hidden observations. The parameters of abundances and correlation are: $m1$=41, $m2$=46, $m3$=33; $\rho_{1-2}$=0.09, $\rho_{1-3}$=-0.42, $\rho_{2-3}$=0.20.

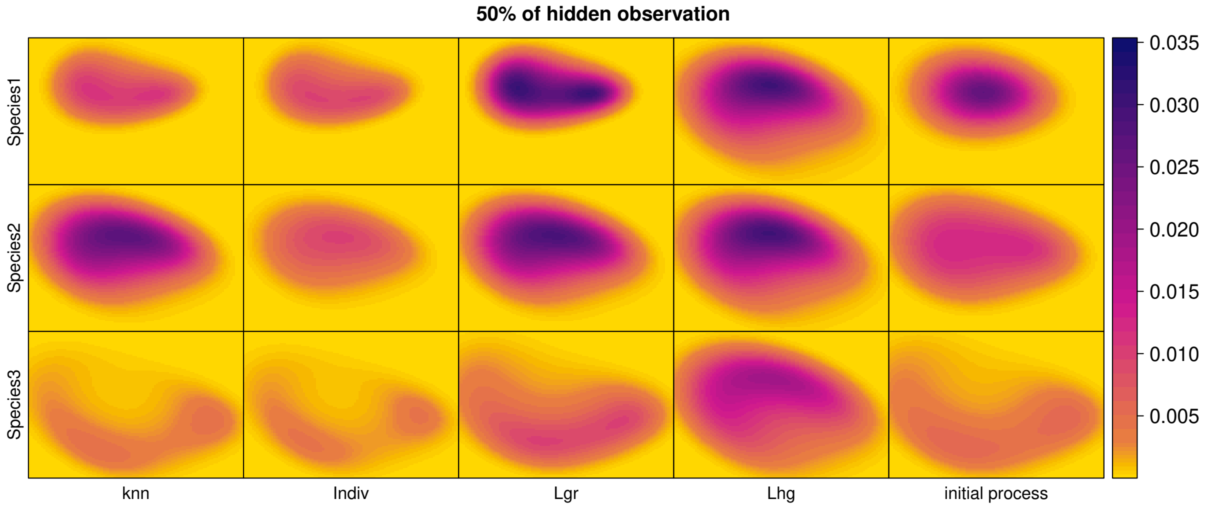


Figure 6: Predicted intensities obtained for the knn, individual, Loop grW and Loop grW initialization methods and the initial intensities from the process at 50% of hidden observations. The parameters of abundances and correlation are:$m1$=41, $m2$=46, $m3$=33; $\rho_{1-2}$=0.09, $\rho_{1-3}$=-0.42, $\rho_{2-3}$=0.20.

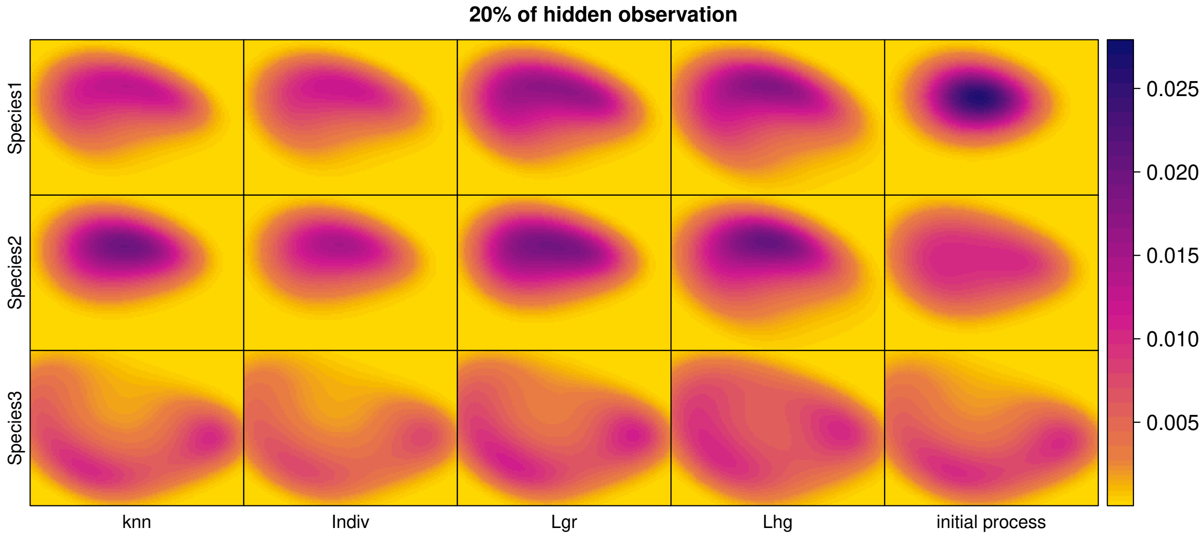


Figure 7: Predicted intensities obtained for the knn, individual, Loop grW and Loop grW initialization methods and the initial intensities from the process at 20% of hidden observations. The parameters of abundances and correlation are: $m1$=39, $m2$=37, $m3$=38; $\rho_{1-2}$=0.09, $\rho_{1-3}$=-0.42, $\rho_{2-3}$=0.20.

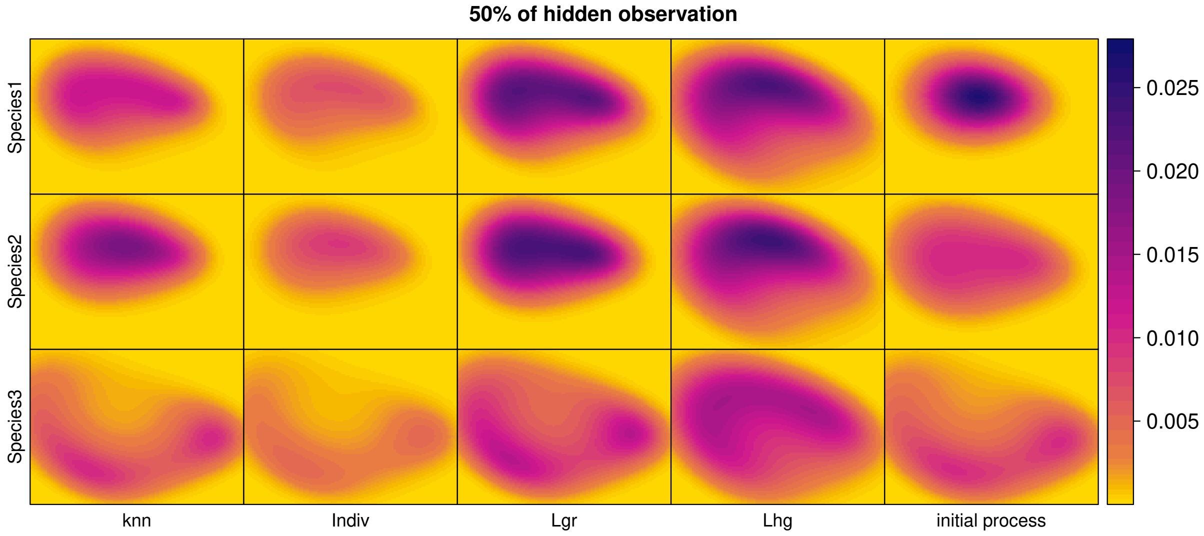


Figure 8: Predicted intensities obtained for the knn, individual, Loop grW and Loop grW initialization methods and the initial intensities from the process at 50% of hidden observations. The parameters of abundances and correlation are:$m1$=39, $m2$=37, $m3$=38; $\rho_{1-2}$=0.09, $\rho_{1-3}$=-0.42, $\rho_{2-3}$=0.20.

##### 0.1.3 Performances for the other methods not presented in the result section

In this section we present the results of measures of performances and predictions of the other methods tested but not shown in the results part. The measures of performances for those methods display worse values than for the individual method. Also, all those methods fail to predict the intensities starting from either at 20% of hidden or 50% of hidden by either overestimating or underestimating the shape of the prediction.

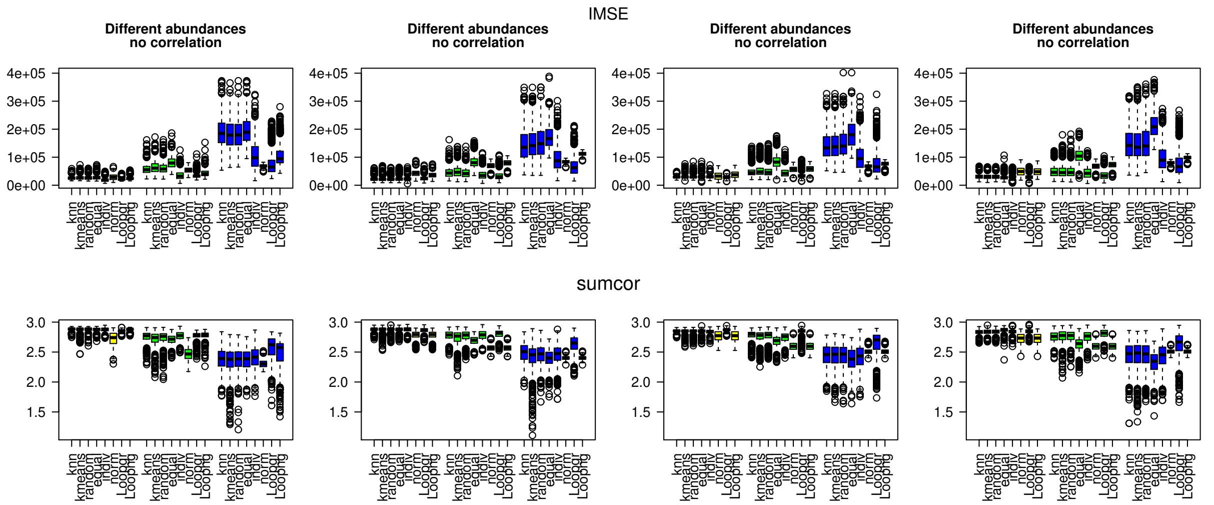


Figure 9: Relabeling measures of performance for all the initialization methods. Each color boxplot represents a different percentage of hidden observations: in yellow are the performances for 20% of hidden observations, in green for 50% and in blue for 80%.

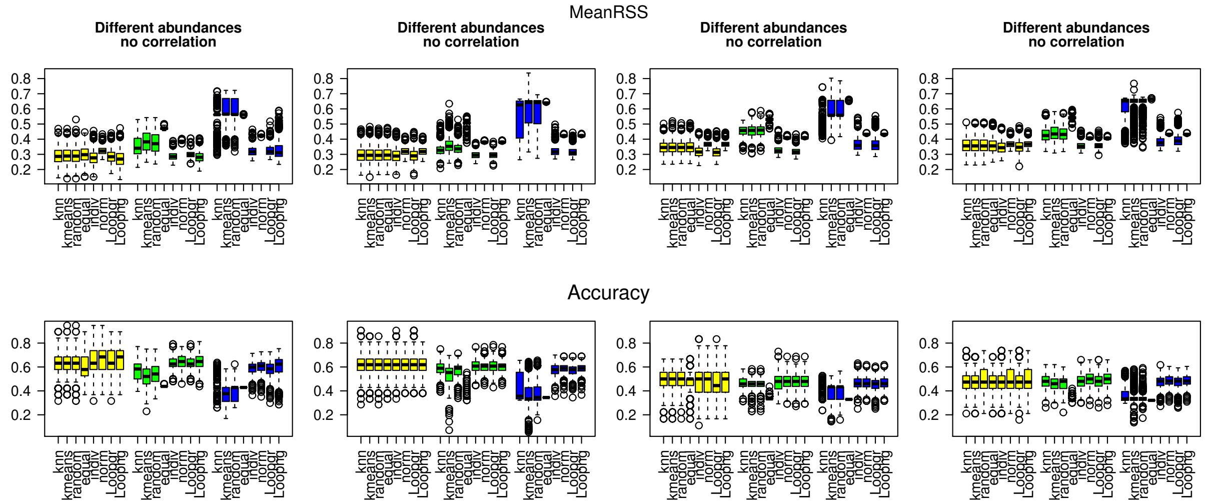


Figure 10: Prediction measures of performance for all the initialization methods. Each color boxplot represents a different percentage of hidden observations: in yellow are the performances for 20% of hidden observations, in green for 50% and in blue for 80%.

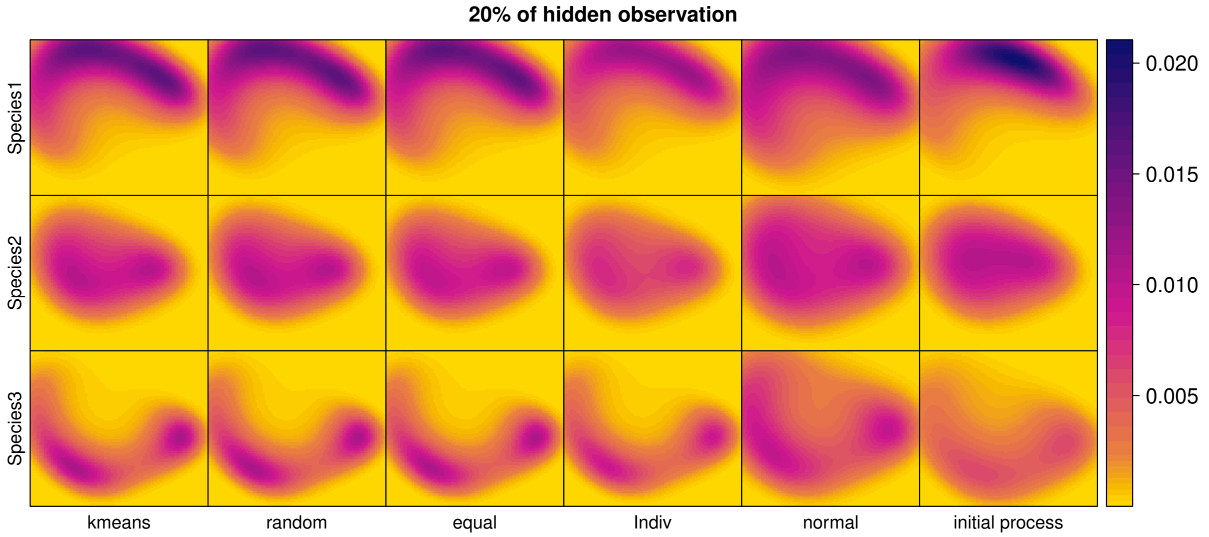


Figure 11: Predicted intensities obtained for the kmeans, random, equal, individual, normal initialization methods and the initial intensities from the process at 20% of hidden observations.The parameters of abundances and correlation are: $m1$=32, $m2$=42, $m3$=23; $\rho_{1-2}$=0.85, $\rho_{1-3}$=-0.09, $\rho_{2-3}$=0.20.

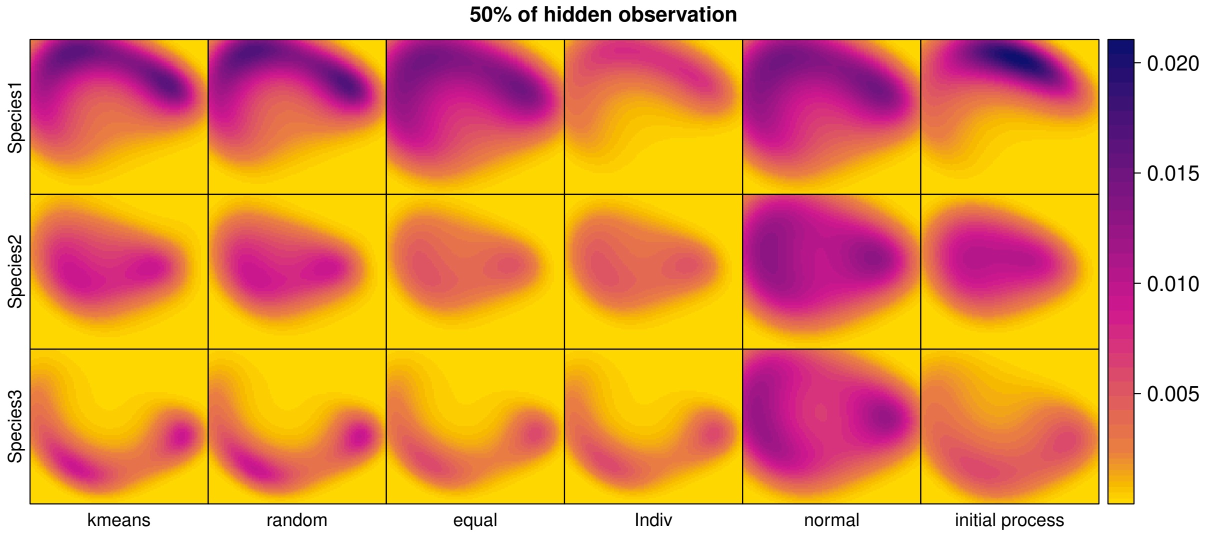


Figure 12: Predicted intensities obtained for the kmeans, random, equal, individual, normal initialization methods and the initial intensities from the process at 50% of hidden observations.The parameters of abundances and correlation are: $m1$=32, $m2$=42, $m3$=23; $\rho_{1-2}$=0.85, $\rho_{1-3}$=-0.09, $\rho_{2-3}$=0.20.

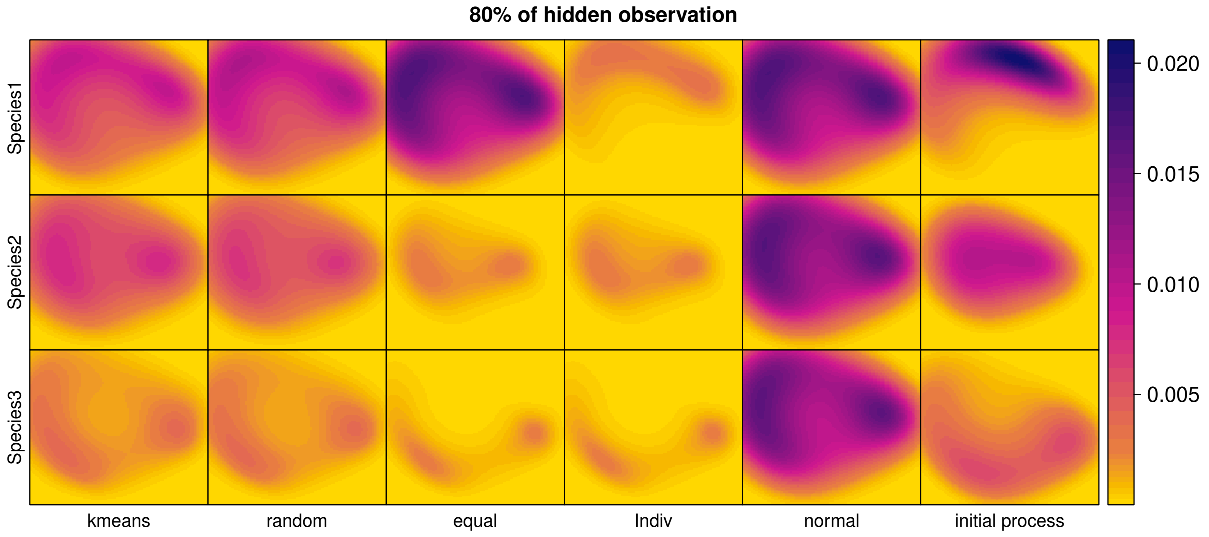


Figure 13: Predicted intensities obtained for the kmeans, random, equal, individual, normal initialization methods and the initial intensities from the process at 80% of hidden observations.The parameters of abundances and correlation are: $m1$=32, $m2$=42, $m3$=23; $\rho_{1-2}$=0.85, $\rho_{1-3}$=-0.09, $\rho_{2-3}$=0.20.

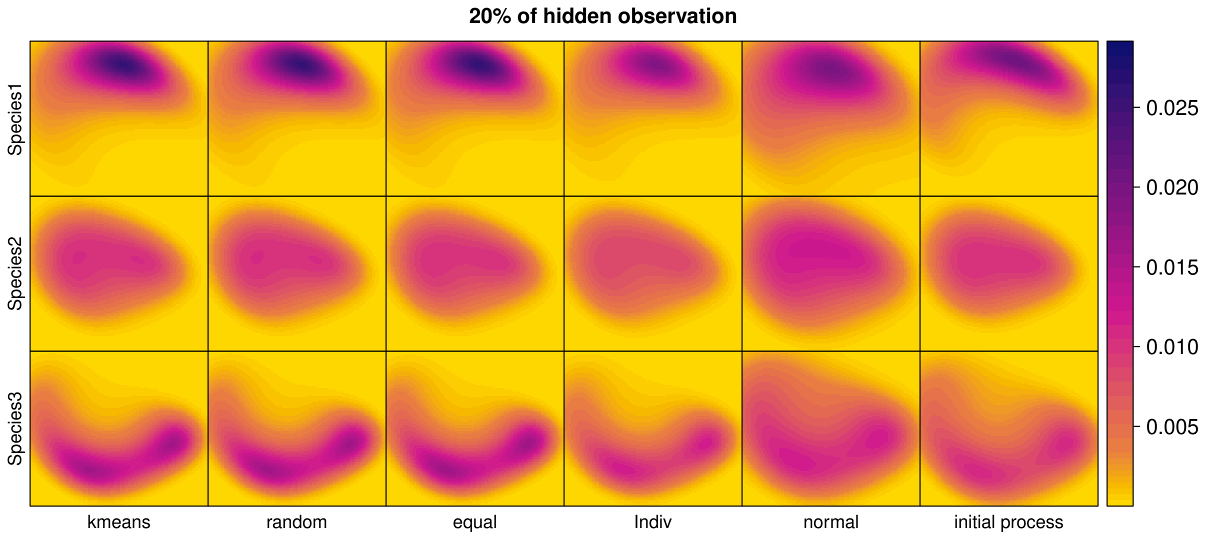


Figure 14: Predicted intensities obtained for the kmeans, random, equal, individual, normal initialization methods and the initial intensities from the process at 20% of hidden observations.The parameters of abundances and correlation are: $m1$=33, $m2$=34, $m3$=35; $\rho_{1-2}$=0.85, $\rho_{1-3}$=-0.09, $\rho_{2-3}$=0.20.

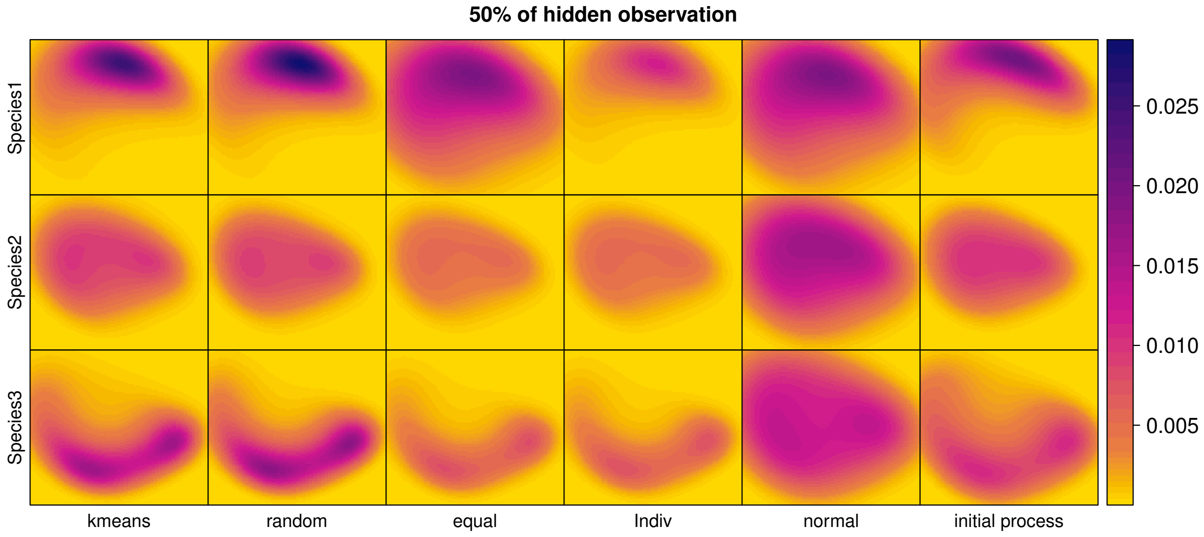


Figure 15: Predicted intensities obtained for the kmeans, random, equal, individual, normal initialization methods and the initial intensities from the process at 50% of hidden observations.The parameters of abundances and correlation are:$m1$=33, $m2$=34, $m3$=35; $\rho_{1-2}$=0.85, $\rho_{1-3}$=-0.09, $\rho_{2-3}$=0.20.

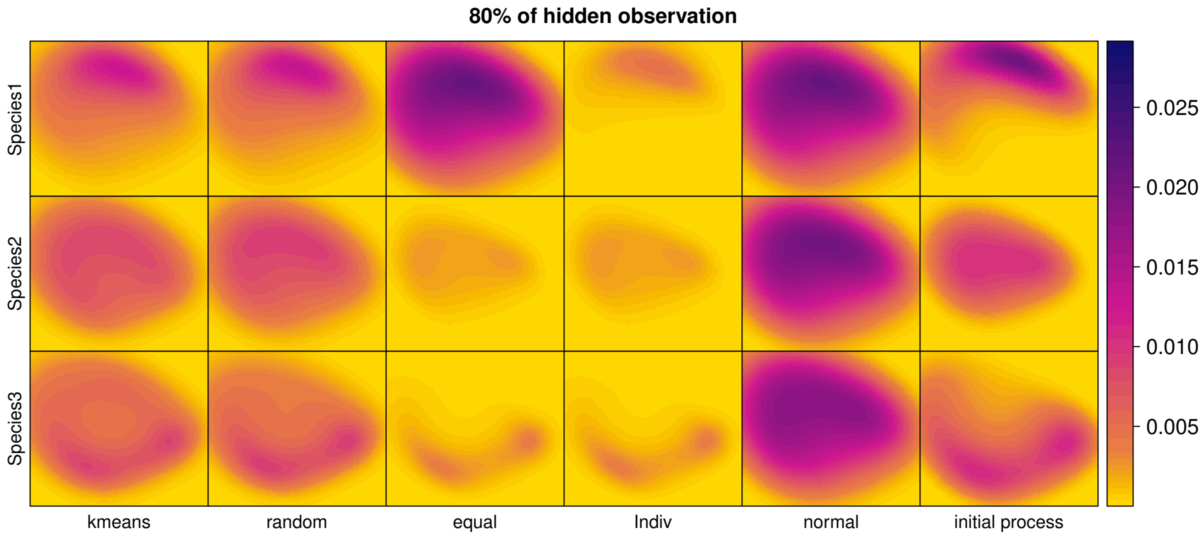


Figure 16: Predicted intensities obtained for the kmeans, random, equal, individual, normal initialization methods and the initial intensities from the process at 80% of hidden observations.The parameters of abundances and correlation are:$m1$=33, $m2$=34, $m3$=35; $\rho_{1-2}$=0.85, $\rho_{1-3}$=-0.09, $\rho_{2-3}$=0.20.

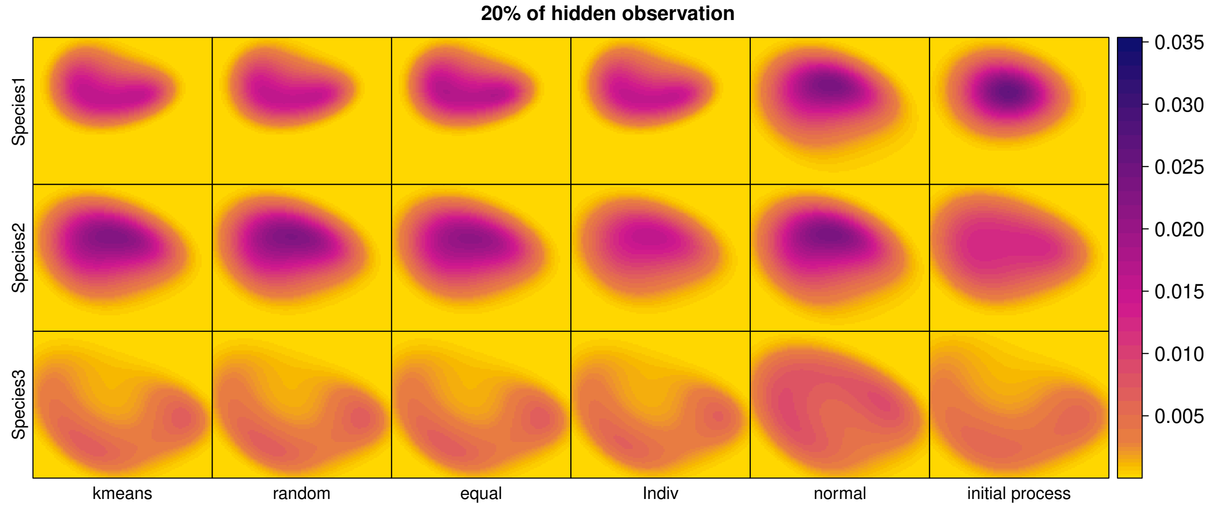


Figure 17: Predicted intensities obtained for the kmeans, random, equal, individual, normal initialization methods and the initial intensities from the process at 20% of hidden observations.The parameters of abundances and correlation are:$m1$=41, $m2$=46, $m3$=33; $\rho_{1-2}$=0.09, $\rho_{1-3}$=-0.42, $\rho_{2-3}$=0.20.

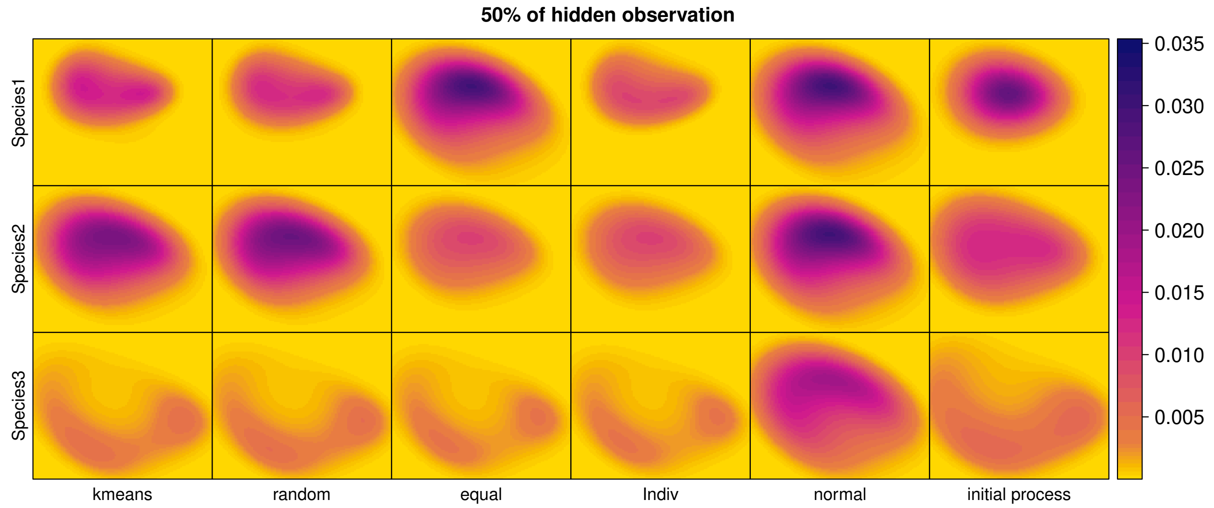


Figure 18: Predicted intensities obtained for the kmeans, random, equal, individual, normal initialization methods and the initial intensities from the process at 50% of hidden observations.The parameters of abundances and correlation are: $m1$=41, $m2$=46, $m3$=33; $\rho_{1-2}$=0.09, $\rho_{1-3}$=-0.42, $\rho_{2-3}$=0.20.

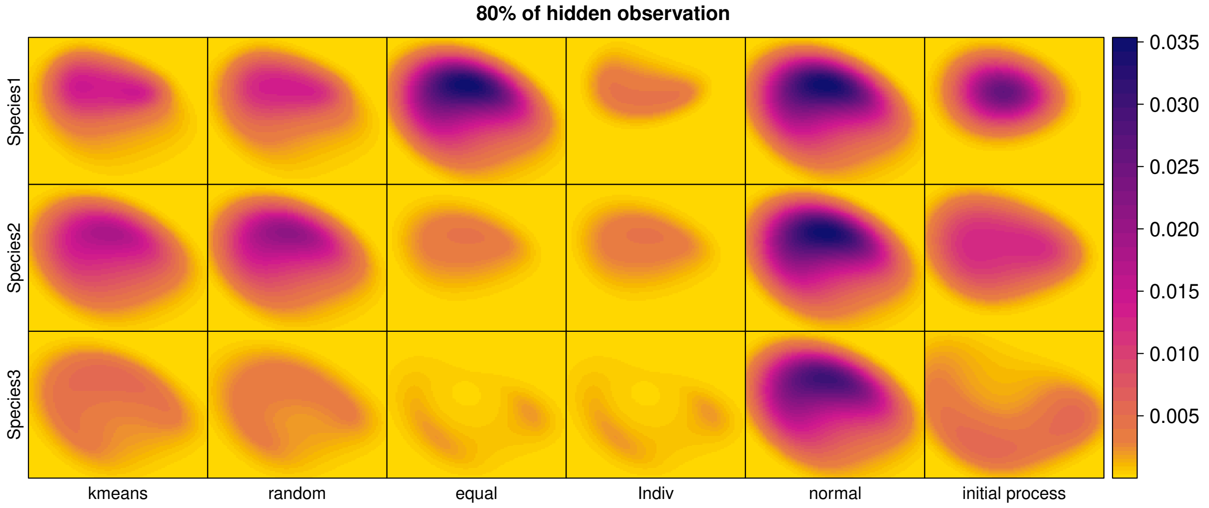


Figure 19: Predicted intensities obtained for the kmeans, random, equal, individual, normal initialization methods and the initial intensities from the process at 80% of hidden observations.The parameters of abundances and correlation are: $m1$=41, $m2$=46, $m3$=33; $\rho_{1-2}$=0.09, $\rho_{1-3}$=-0.42, $\rho_{2-3}$=0.20.

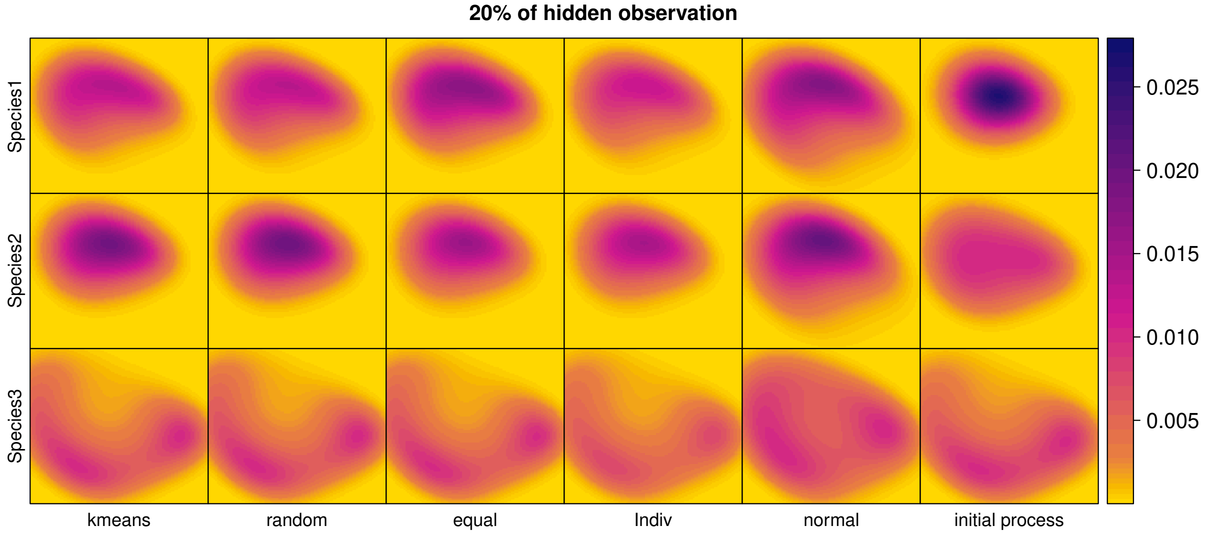


Figure 20: Predicted intensities obtained for the kmeans, random, equal, individual, normal initialization methods and the initial intensities from the process at 20% of hidden observations.The parameters of abundances and correlation are: $m1$=39, $m2$=37, $m3$=38; $\rho_{1-2}$=0.09, $\rho_{1-3}$=-0.42, $\rho_{2-3}$=0.20.

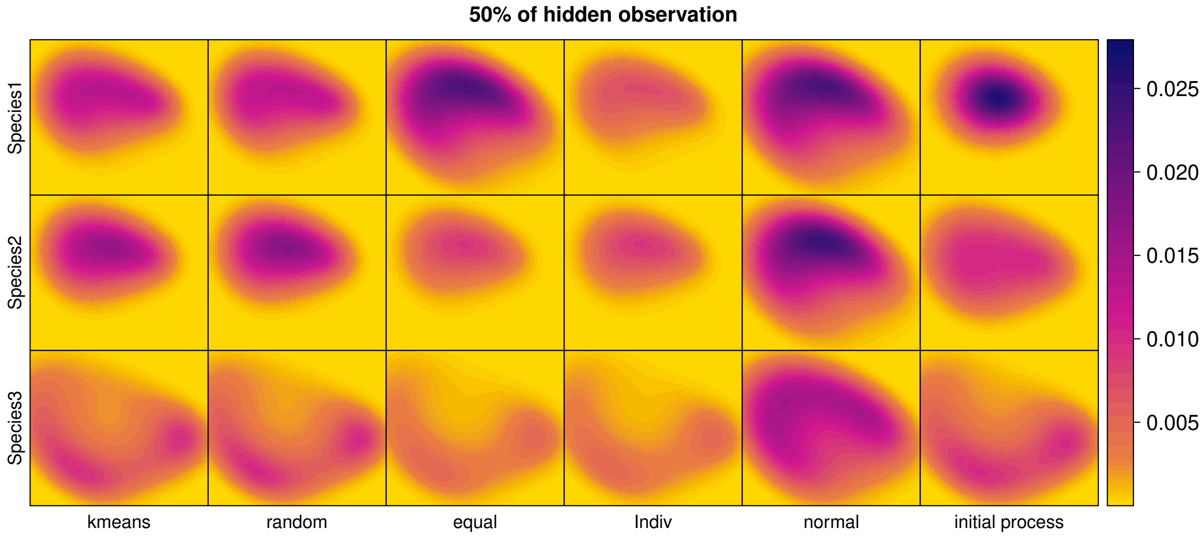


Figure 21: Predicted intensities obtained for the kmeans, random, equal, individual, normal initialization methods and the initial intensities from the process at 50% of hidden observations.The parameters of abundances and correlation are: $m1$=39, $m2$=37, $m3$=38; $\rho_{1-2}$=0.09, $\rho_{1-3}$=-0.42, $\rho_{2-3}$=0.20.

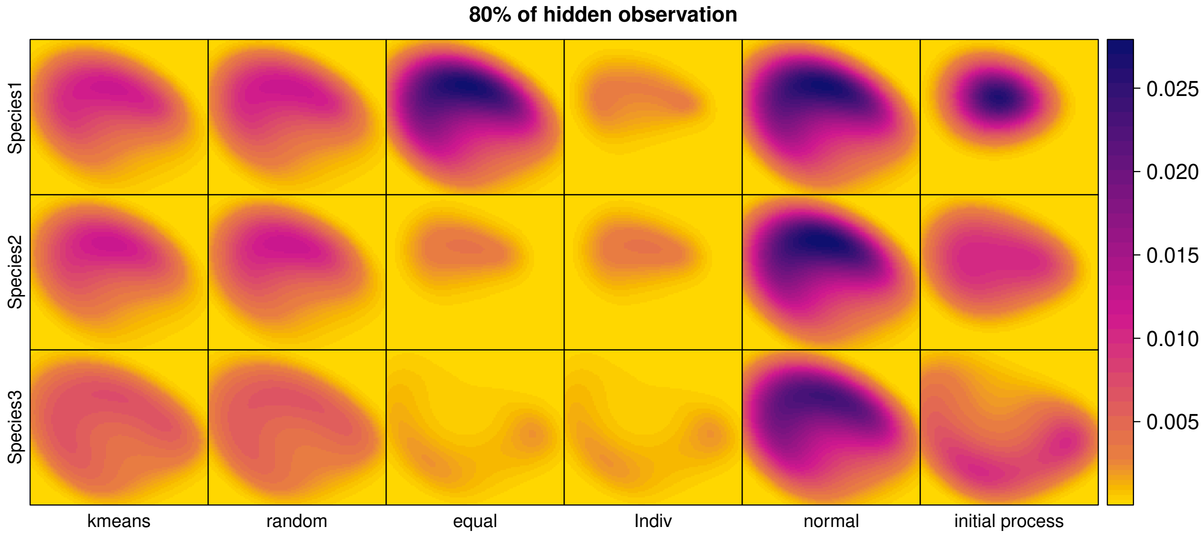


Figure 22: Predicted intensities obtained for the kmeans, random, equal, individual, normal initialization methods and the initial intensities from the process at 80% of hidden observations.The parameters of abundances and correlation are: $m1$=39, $m2$=37, $m3$=38; $\rho_{1-2}$=0.09, $\rho_{1-3}$=-0.42, $\rho_{2-3}$=0.20.

##### 0.1.4 Model parameters variation

Most of the results presented in the the results for the varying model parameters can be extended to all combination of abundances and correlation tested, especially for the knn and Loop hgW methods. For Loop grW method and for similar abundances not matter if the distribution are correlated or not, the $\delta_{\max}$ threshold doesn't vary in performances for all measures. Across the different combination of abundances and correlation tested, $\delta_{\min}$=0.1 show better performances. $\delta_{\text{step}}$, In the case where we have highly correlated distribution, the threshold 0.1 displays better performances for prediction than when the correlation is lower for IMSE and sumcor. Similarly for the threshold value 0.05 for MeanRSS and acc.

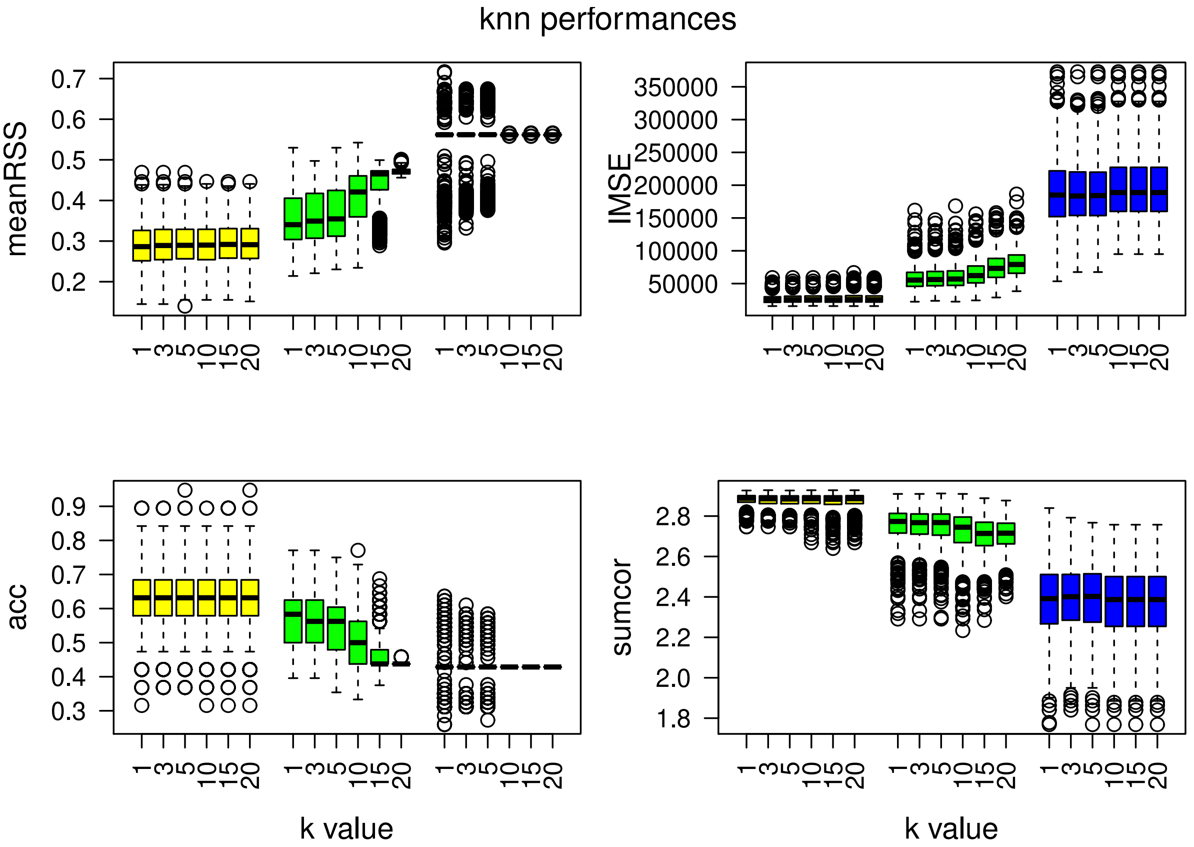


Figure 23: Model performances for the knn method. Each color boxplot represents a different percentage of hidden observations: in yellow are the performances for 20% of hidden observations, in green for 50% and in blue for 80%. The parameters of abundances and correlation are: $m1$=42, $m2$=31, $m3$=25; $\rho_{1-2}$=0.85, $\rho_{1-3}$=-0.09, $\rho_{2-3}$=0.20.

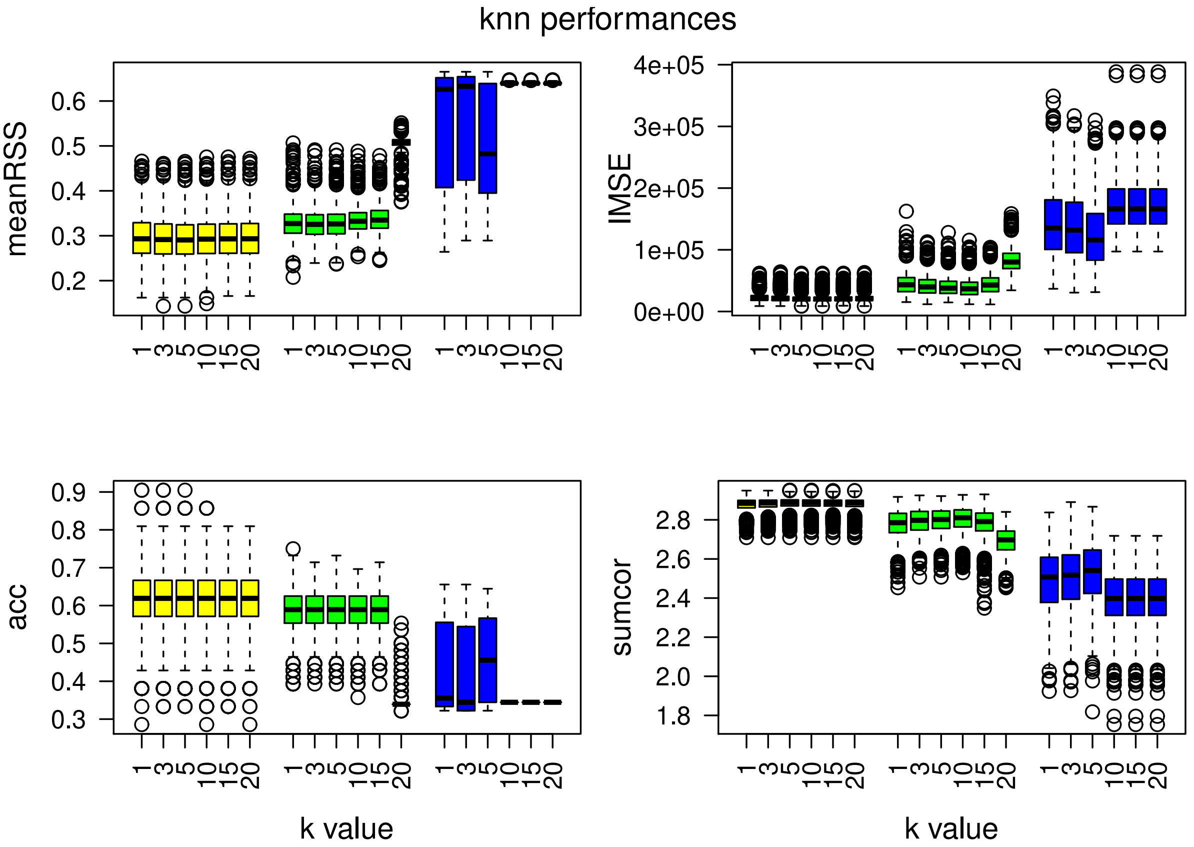


Figure 24: Model performances for the knn method. Each color boxplot represents a different percentage of hidden observations: in yellow are the performances for 20% of hidden observations, in green for 50% and in blue for 80%. The parameters of abundances and correlation are: $m1$=39, $m2$=37, $m3$=38; $\rho_{1-2}$=0.09, $\rho_{1-3}$=-0.42, $\rho_{2-3}$=0.20.

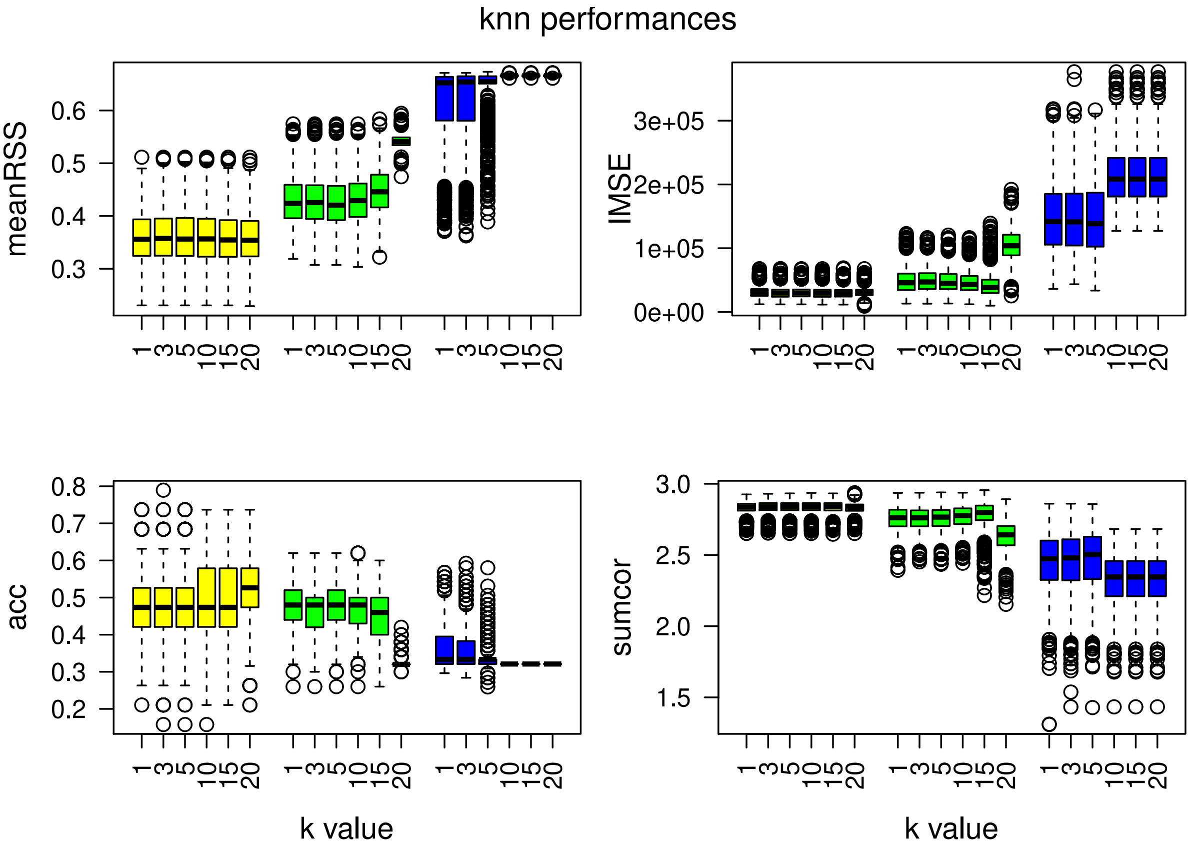


Figure 25: Model performances for the knn method. Each color boxplot represents a different percentage of hidden observations: in yellow are the performances for 20% of hidden observations, in green for 50% and in blue for 80%. The parameters of abundances and correlation are: $m1$=33, $m2$=34, $m3$=35; $\rho_{1-2}$=0.85, $\rho_{1-3}$=-0.09, $\rho_{2-3}$=0.20.

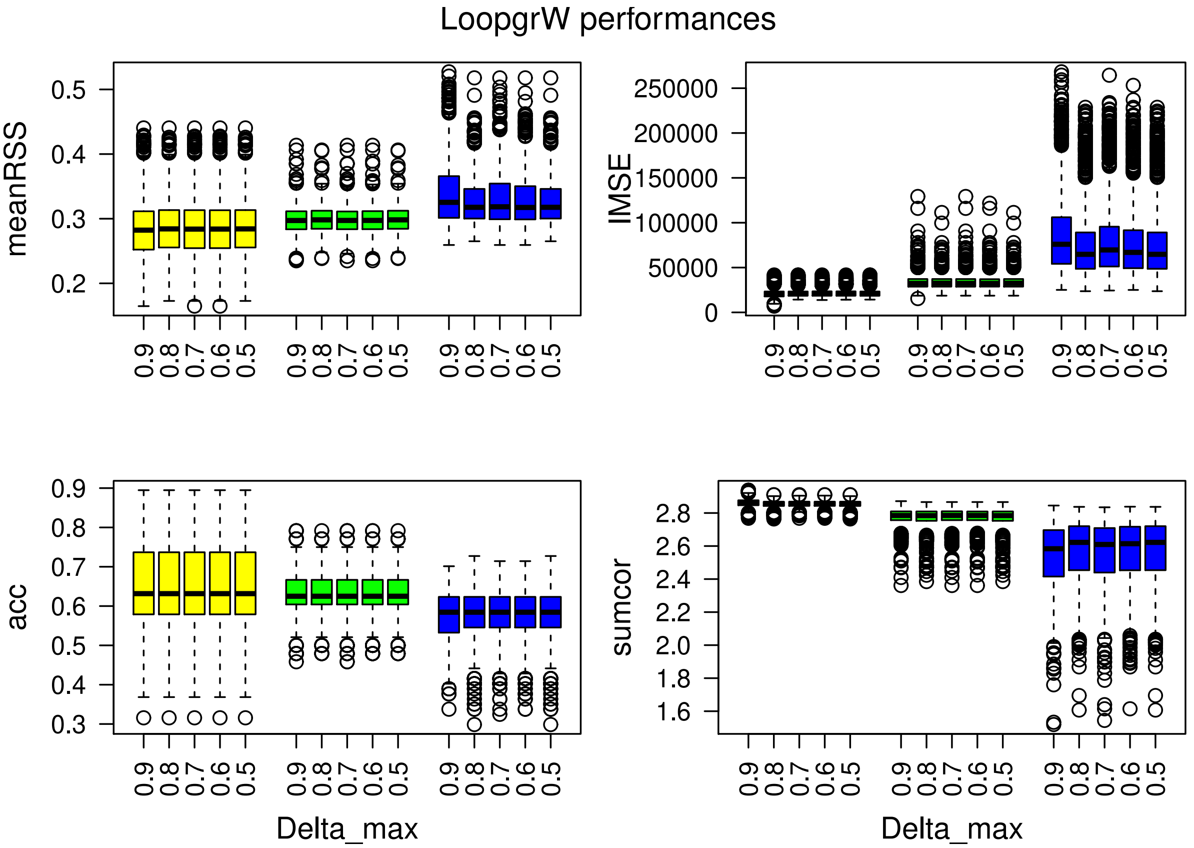


Figure 26: Model performances for the Loop grW method and different $w_{min}$. Each color boxplot represents a different percentage of hidden observations: in yellow are the performances for 20% of hidden observations, in green for 50% and in blue for 80%. The parameters of abundances and correlation are: $m1$=42, $m2$=31, $m3$=25; $\rho_{1-2}$=0.85, $\rho_{1-3}$=-0.09, $\rho_{2-3}$=0.20.

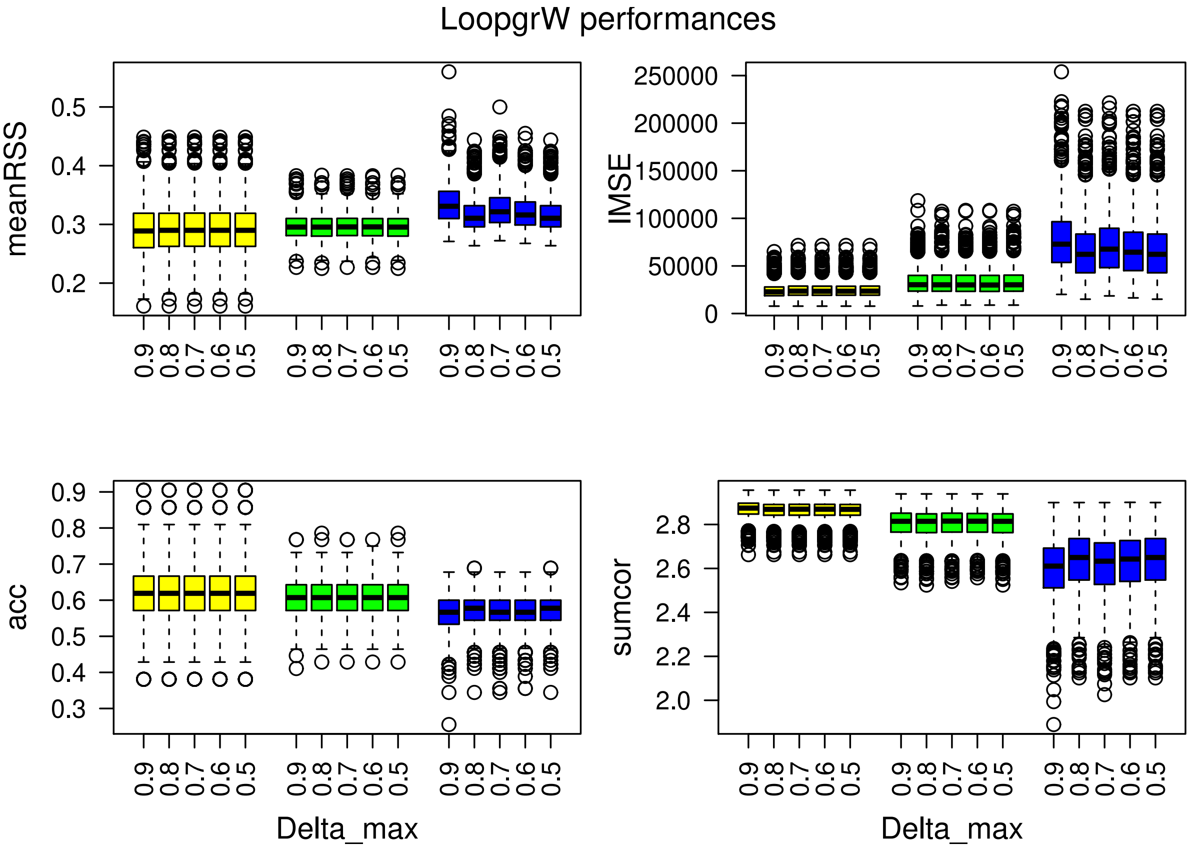


Figure 27: Model performances for the Loop grW method and different $w_{min}$. Each color boxplot represents a different percentage of hidden observations: in yellow are the performances for 20% of hidden observations, in green for 50% and in blue for 80%. The parameters of abundances and correlation are: $m1$=39, $m2$=37, $m3$=38; $\rho_{1-2}$=0.09, $\rho_{1-3}$=-0.42, $\rho_{2-3}$=0.20.

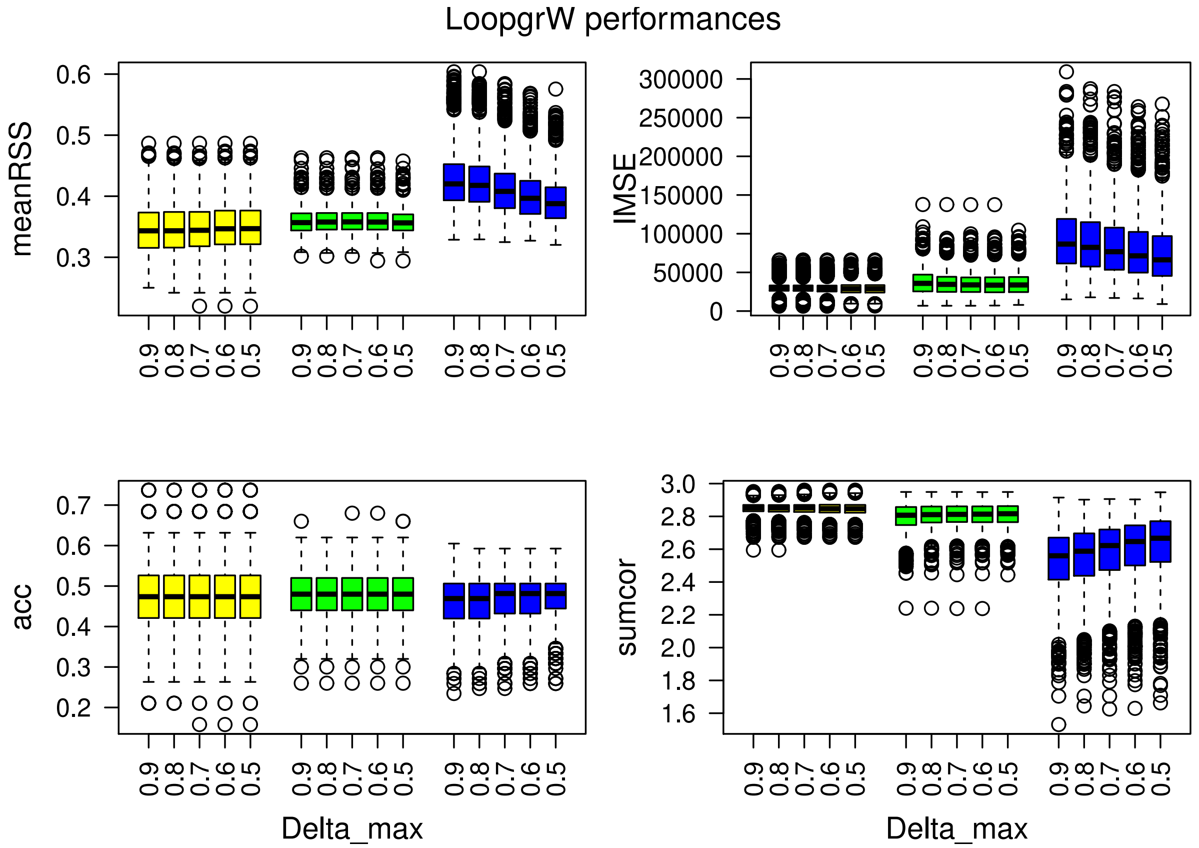


Figure 28: Model performances for the Loop grW method and different $w_{min}$. Each color boxplot represents a different percentage of hidden observations: in yellow are the performances for 20% of hidden observations, in green for 50% and in blue for 80%. The parameters of abundances and correlation are: $m1$=33, $m2$=34, $m3$=35; $\rho_{1-2}$=0.85, $\rho_{1-3}$=-0.09, $\rho_{2-3}$=0.20.

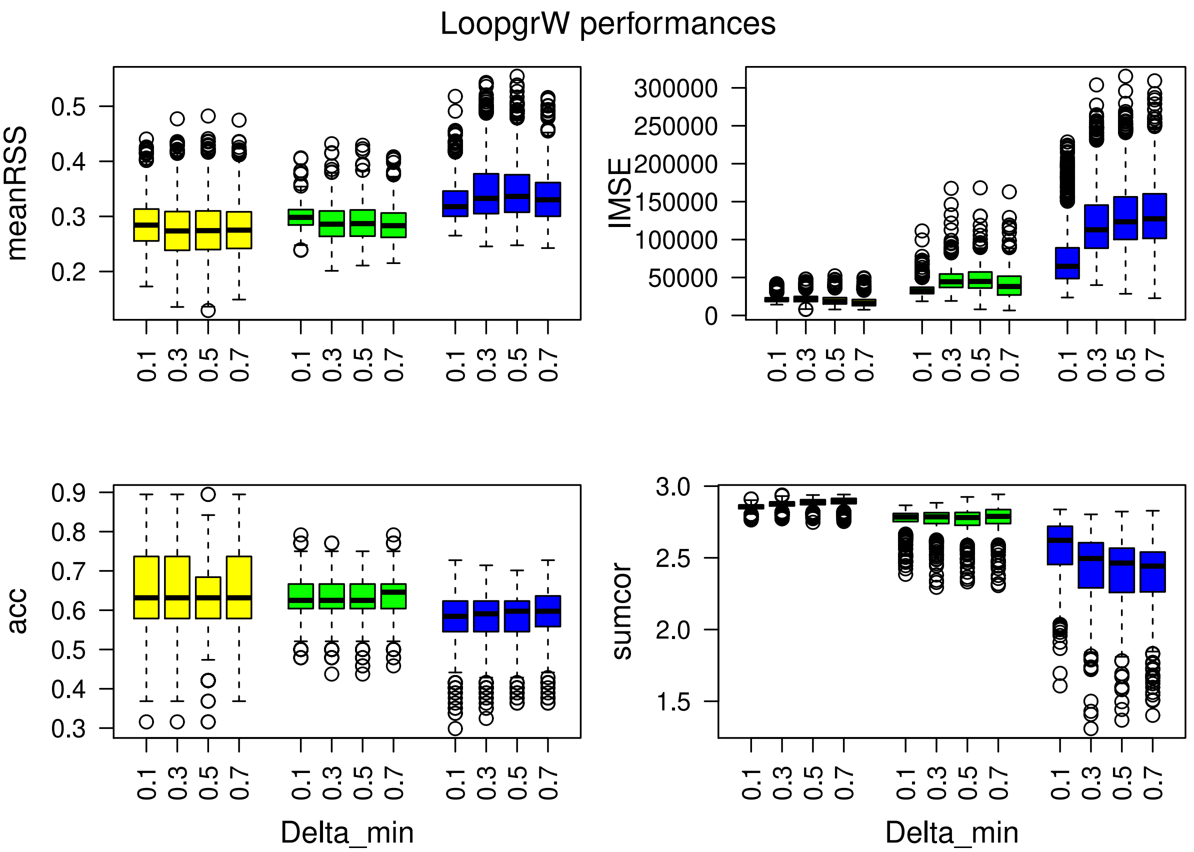


Figure 29: Model performances for the Loop grW method and different $\delta_{\min}$. Each color boxplot represents a different percentage of hidden observations: in yellow are the performances for 20% of hidden observations, in green for 50% and in blue for 80%. The parameters of abundances and correlation are: $m1$=42, $m2$=31, $m3$=25; $\rho_{1-2}$=0.85, $\rho_{1-3}$=-0.09, $\rho_{2-3}$=0.20.

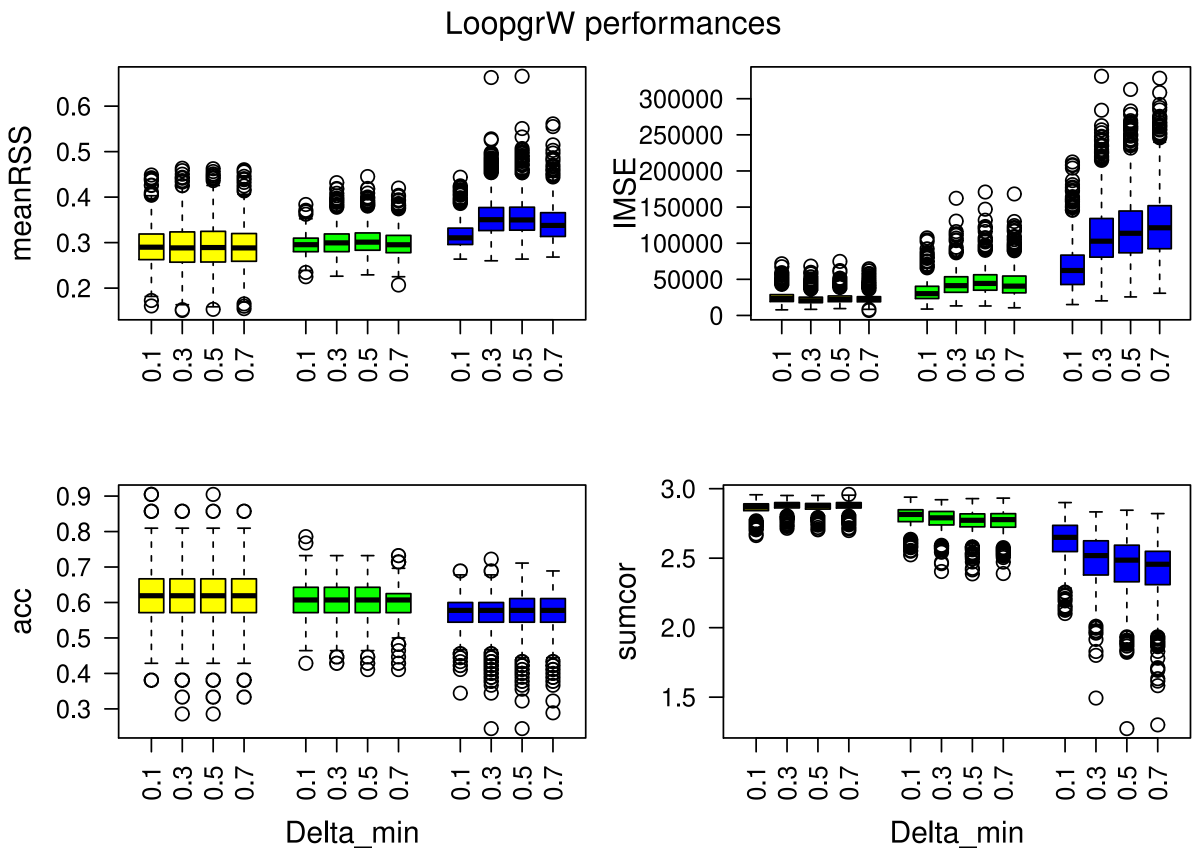


Figure 30: Model performances for the Loop grW method and different $\delta_{\min}$. Each color boxplot represents a different percentage of hidden observations: in yellow are the performances for 20% of hidden observations, in green for 50% and in blue for 80%. The parameters of abundances and correlation are: $m1$=39, $m2$=37, $m3$=38; $\rho_{1-2}$=0.09, $\rho_{1-3}$=-0.42, $\rho_{2-3}$=0.20.

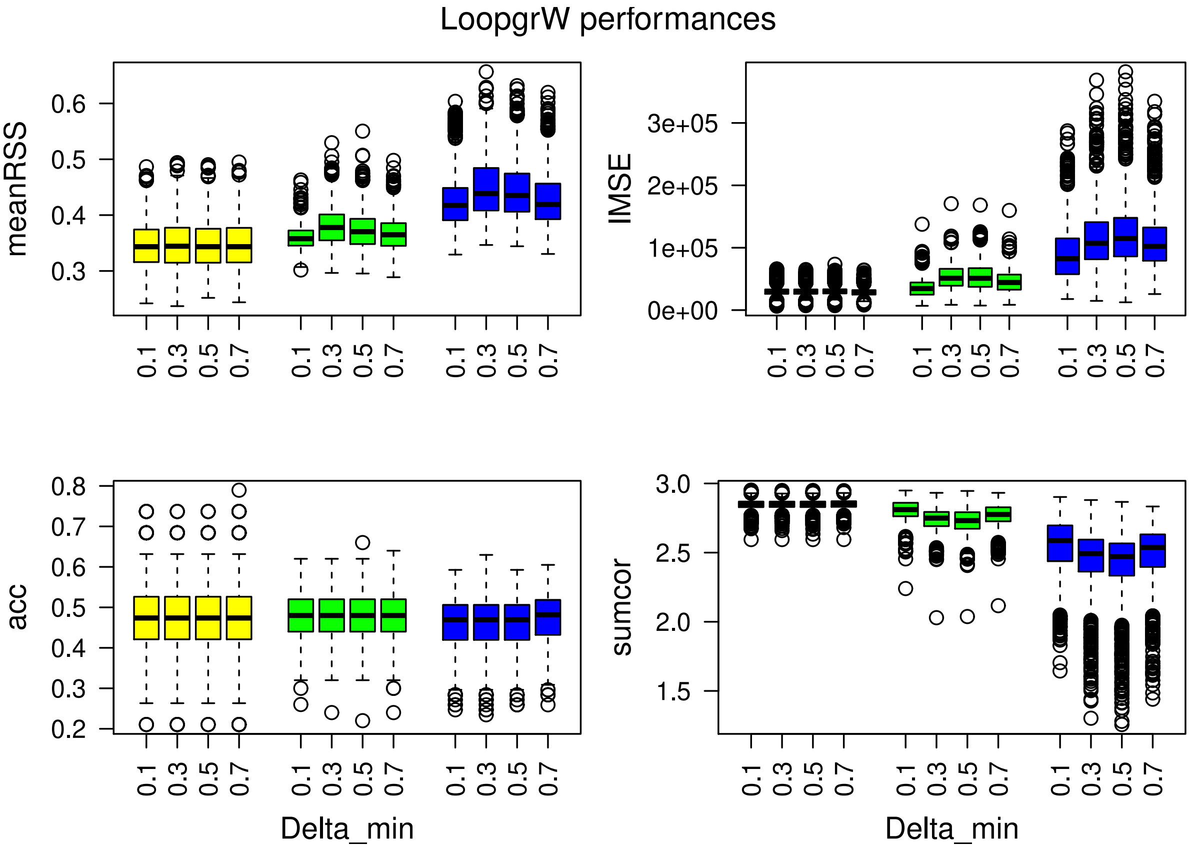


Figure 31: Model performances for the Loop grW method and different $\delta_{\mathrm{min}}$. Each color boxplot represents a different percentage of hidden observations: in yellow are the performances for 20% of hidden observations, in green for 50% and in blue for 80%. The parameters of abundances and correlation are: $m1$=33, $m2$=34, $m3$=35; $\rho_{1-2}$=0.85, $\rho_{1-3}$=-0.09, $\rho_{2-3}$=0.20.

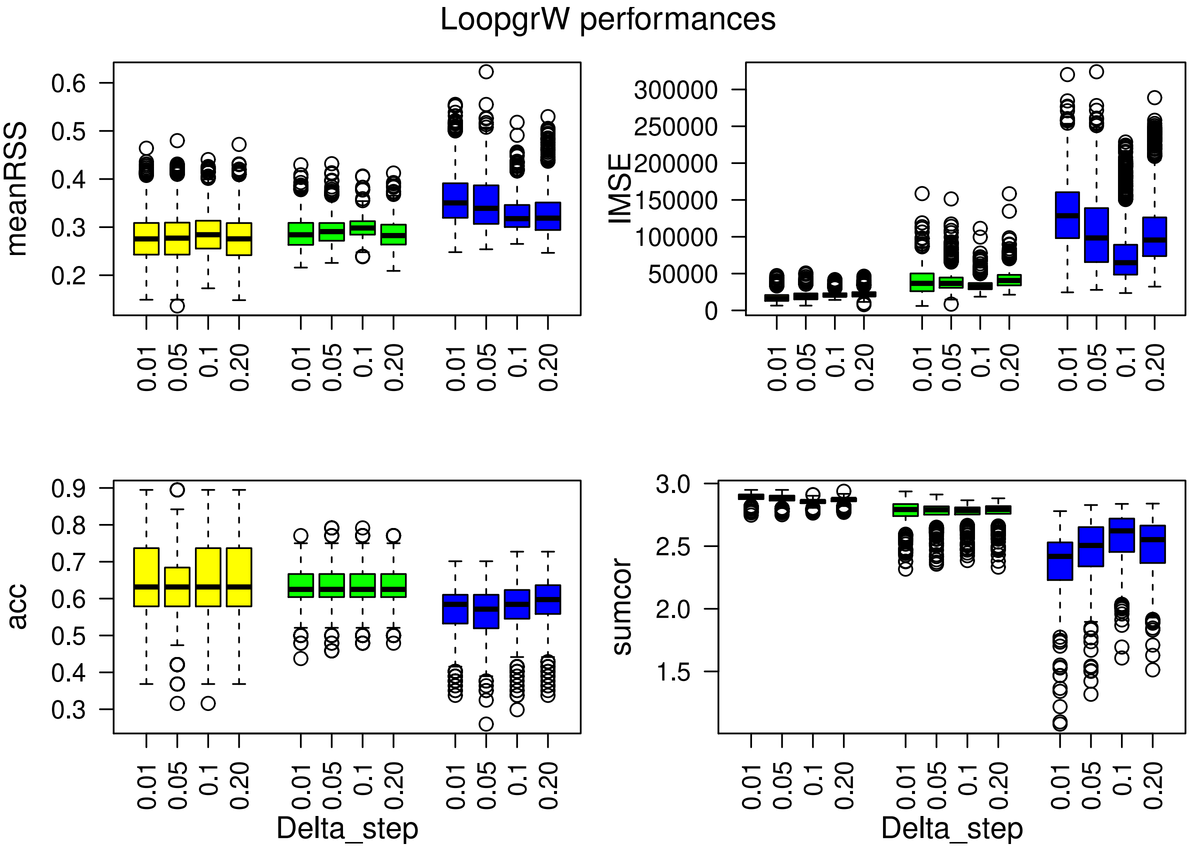


Figure 32: Model performances for the Loop grW method and different weight step $\delta_{\text{step}}$. Each color boxplot represents a different percentage of hidden observations: in yellow are the performances for 20% of hidden observations, in green for 50% and in blue for 80%. The parameters of abundances and correlation are: $m1$=42, $m2$=31, $m3$=25; $\rho_{1-2}$=0.85, $\rho_{1-3}$=-0.09, $\rho_{2-3}$=0.20.

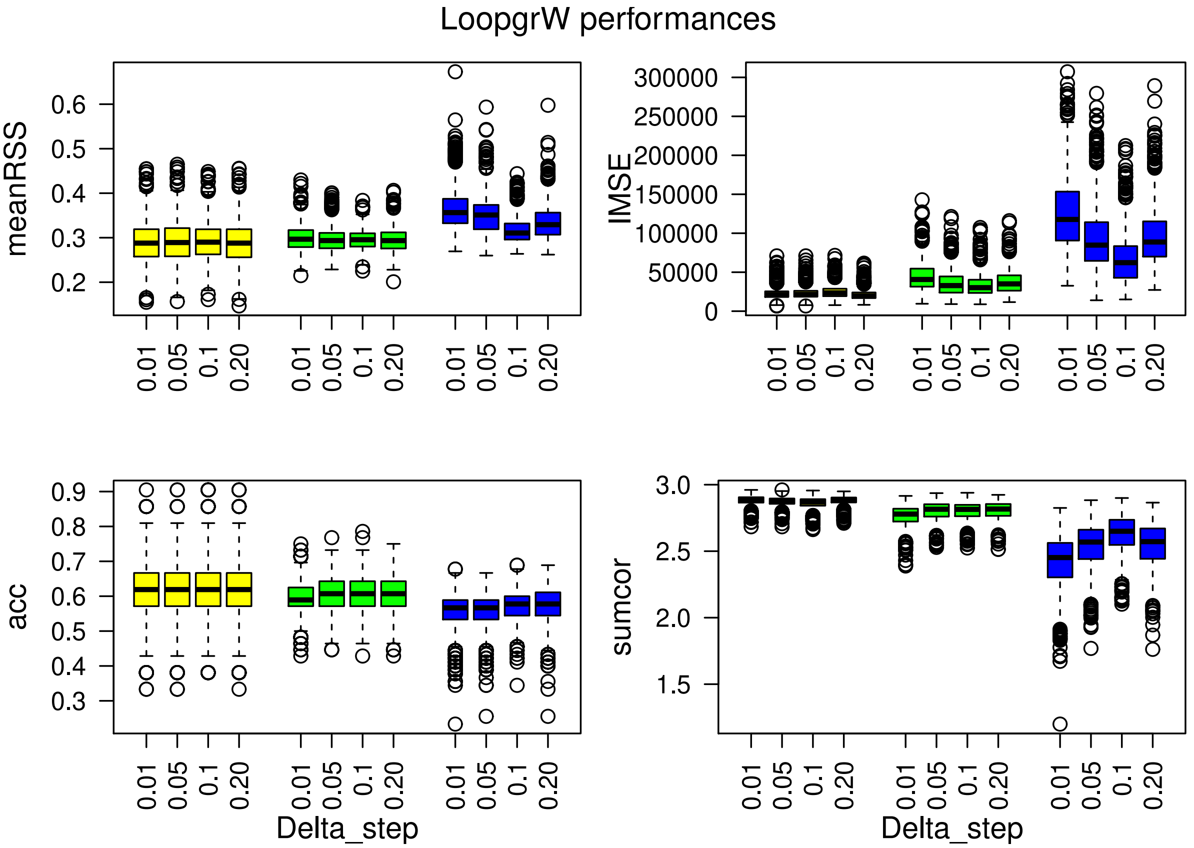


Figure 33: Model performances for the Loop grW method and different weight step $\delta_{\text{step}}$. Each color boxplot represents a different percentage of hidden observations: in yellow are the performances for 20% of hidden observations, in green for 50% and in blue for 80%. The parameters of abundances and correlation are: $m1$=39, $m2$=37, $m3$=38; $\rho_{1-2}$=0.09, $\rho_{1-3}$=-0.42, $\rho_{2-3}$=0.20.

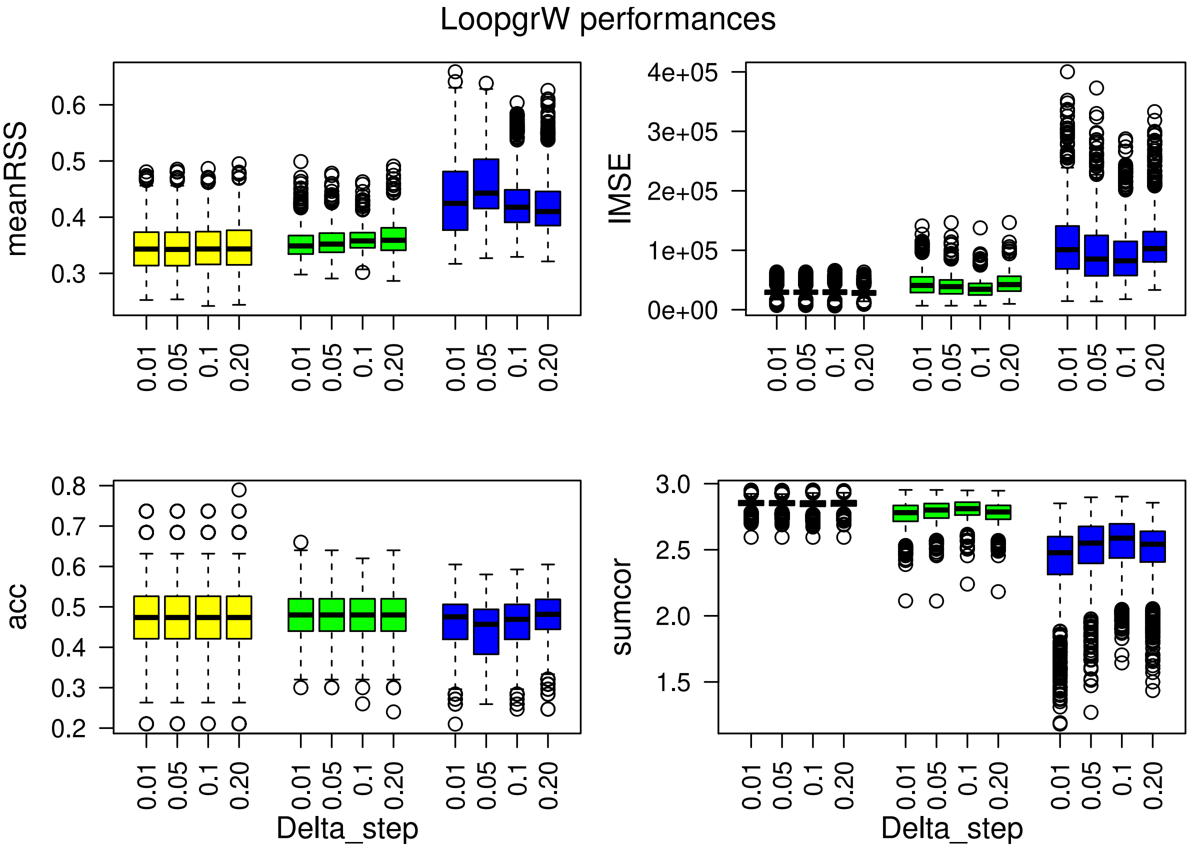


Figure 34: Model performances for the Loop grW method and different weight step $\delta_{\text{step}}$. Each color boxplot represents a different percentage of hidden observations: in yellow are the performances for 20% of hidden observations, in green for 50% and in blue for 80%. The parameters of abundances and correlation are: $m1$=33, $m2$=34, $m3$=35; $\rho_{1-2}$=0.85, $\rho_{1-3}$=-0.09, $\rho_{2-3}$=0.20.

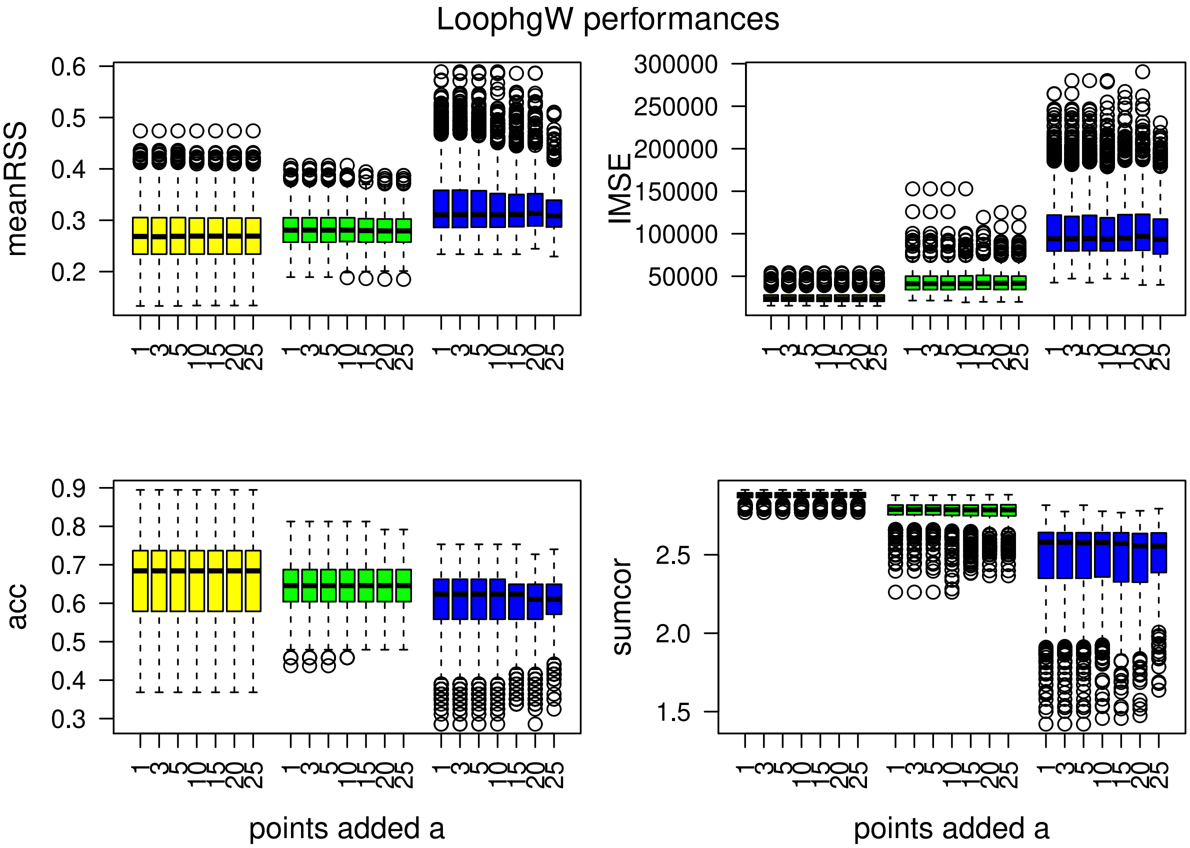


Figure 35: Model performances for the Lhg method. Each color boxplot represents a different percentage of hidden observations: in yellow are the performances for 20% of hidden observations, in green for 50% and in blue for 80%. The parameters of abundances and correlation are: $m1$=42, $m2$=31, $m3$=25; $\rho_{1-2}$=0.85, $\rho_{1-3}$=-0.09, $\rho_{2-3}$=0.20.

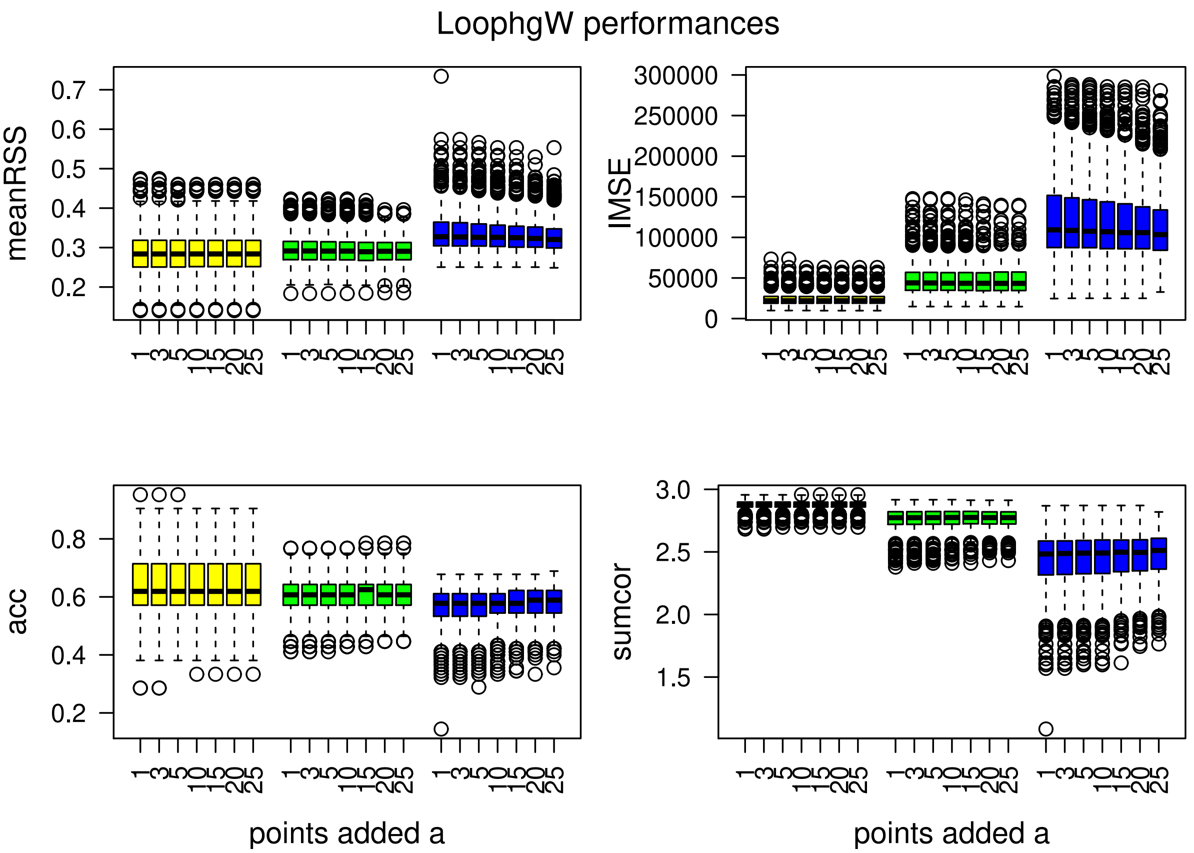


Figure 36: Model performances for the Lhg method. Each color boxplot represents a different percentage of hidden observations: in yellow are the performances for 20% of hidden observations, in green for 50% and in blue for 80%. The parameters of abundances and correlation are: $m1$=39, $m2$=37, $m3$=38; $\rho_{1-2}$=0.09, $\rho_{1-3}$=-0.42, $\rho_{2-3}$=0.20.

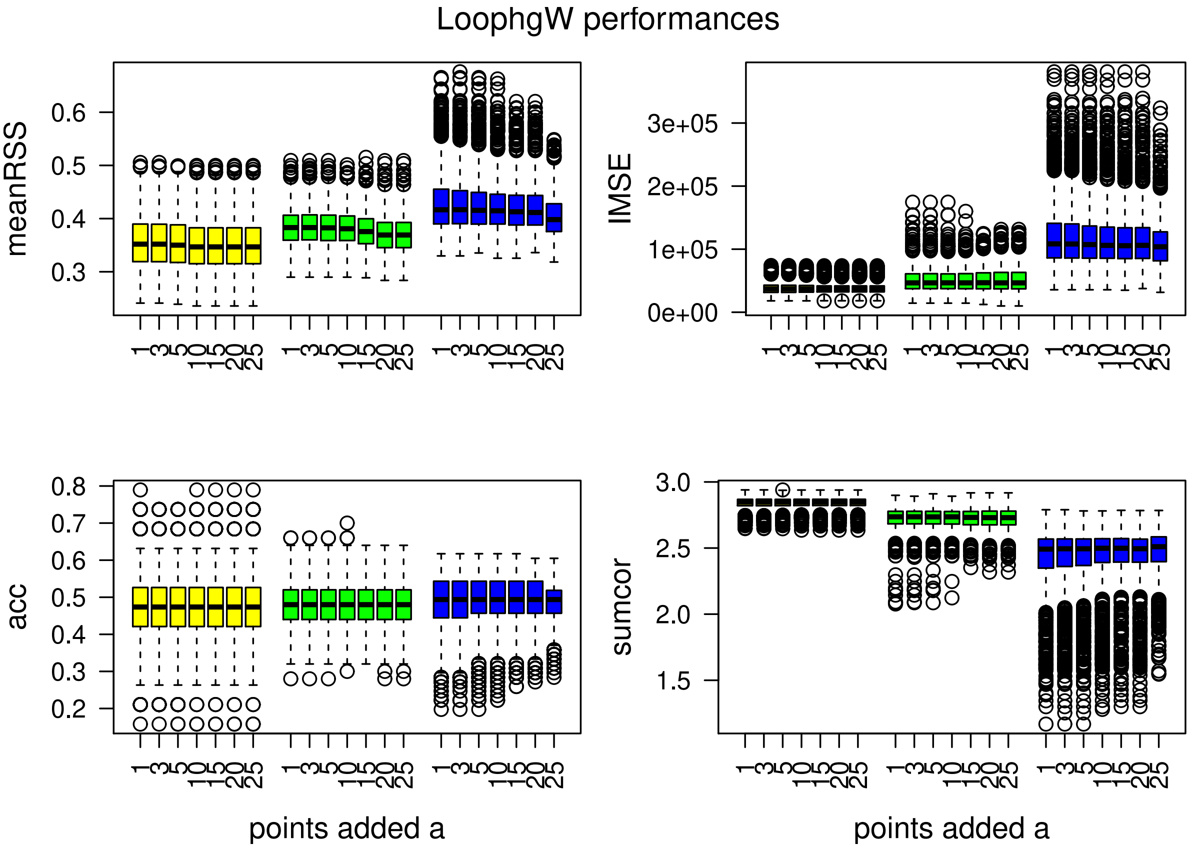


Figure 37: Model performances for the Lhg method. Each color boxplot represents a different percentage of hidden observations: in yellow are the performances for 20% of hidden observations, in green for 50% and in blue for 80%. The parameters of abundances and correlation are: $m1$=33, $m2$=34, $m3$=35; $\rho_{1-2}$=0.85, $\rho_{1-3}$=-0.09, $\rho_{2-3}$=0.20.
